# MAPR origins reveal a new class of prokaryotic cytochrome b_5_ proteins and possible role in eukaryogenesis

**DOI:** 10.1101/2021.11.17.468889

**Authors:** Daniel Tamarit, Sarah Teakel, Michealla Marama, David Aragão, Svetlana Y. Gerdes, Jade K. Forwood, Thijs J. G. Ettema, Michael A. Cahill

## Abstract

The multiple functions of PGRMC1, the archetypal heme-binding eukaryotic MAPR family member, include steroidogenic regulation, membrane trafficking, and steroid responsiveness. The interrelationships between these functions are currently poorly understood. Previous work has shown that different MAPR subclasses were present early in eukaryotic evolution, and that tyrosine phosphorylated residues appeared in the eumetazoan ancestor, coincident with a gastrulation organizer. Here we show that MAPR proteins are related to a newly recognized class of prokaryotic cytochrome-b_5_ domain proteins. Our first solved structure of this new class exhibits shared MAPR-like folded architecture and heme-binding orientation. We also report that a protein subgroup from Candidate Phyla Radiation bacteria shares MAPR-like heme-interacting tyrosines. Our results support bacterial origins for both PGRMC1 and CYP51A, that catalyze the meiosis-associated 14-demethylation of the first sterol lanosterol from yeast to humans. We propose that eukaryotic acquisition of a membrane-trafficking function related to sterol metabolism was associated with the appearance of MAPR genes early in eukaryotic evolution.

## 1. Introduction

Progesterone Receptor Membrane Component 1 (PGRMC1) is the archetypal member of the eukaryotic membrane-associated progesterone receptor (MAPR) family of cytochrome b_5_ (cytb_5_)-related proteins (Cahill, 2007). MAPR proteins are defined within the cytb_5_-superfamily by tyrosinate heme chelation with unique orientation of the heme (Kabe et al., 2016a; Kaluka et al., 2015; Thompson et al., 2007), and by the presence of a small insertion, the MAPR-specific inter-helical insertion region (MIHIR), between helices 3 and 4 of the canonical cytb_5_-domain (Cahill, 2007, 2017; Mifsud and Bateman, 2002).

Mammalian MAPR members include Neuferricin (NEUFC), Neudesin (NENF), PGRMC1 and PGRMC2. The latter diverged prior to the evolution of cartilaginous fish, and share many PGRMC properties. It remains unclear how they differ in function. All three MAPR families (PGRMC, NEUFC, NENF) were present in the Opisthikont ancestor of fungi and animals. All three families are also expressed in the nervous system (Hehenberger et al., 2020), and interact with cytochrome P450 (cyP450) enzymes (Ryu et al., 2017). Otherwise, NEUFC and NENF function is poorly understood: for discussion see Hehenberger et al. (2020).

PGRMC1 is by far the best characterized MAPR protein, and has diverse functions (Cahill et al., 2016). These include some that predate or are potentially ancient in eukaryotes, such as regulation of heme synthesis (Piel et al., 2016), cyP450 interactions (Ryu et al., 2017), and sterol metabolism (Cahill and Medlock, 2017). Some functions arose in eukaryotes, such as membrane trafficking (Cahill et al., 2016; Hampton et al., 2018), cell cycle regulation at the G_0_/G_1_ checkpoint (Cahill et al., 2016; Griffin et al., 2014; Peluso et al., 2014), and mitotic/meiotic spindle association (Juhlen et al., 2018; Luciano and Peluso, 2016; Terzaghi et al., 2016). The eukaryotic phylogenetic distribution of these properties remains unexplored. Other functions are specialized metazoan developments, including roles in e.g. fertility, embryogenic axon guidance, and membrane trafficking associated with synaptic plasticity (Cahill et al., 2016; Izzo et al., 2014).

MAPR cyP450-interactions (Ryu et al., 2017) conspicuously feature PGRMC1 regulation of lanosterol-14-demethylase, the most conserved eukaryotic cyP450 (CYP51A), to modify the first sterol (lanosterol) in yeast and mammals in an oxygen requiring reaction (Fig. 1) (Cahill, 2007; Hughes et al., 2007b). McGuire et al. recently reported that PGRMC1 binding to this and multiple cyP450s leads to their stabilization, elevating the protein levels of cyP450s which occurs even for heme-binding deficient PGRMC1 mutants. They argue, but do not demonstrate, that the pentacoordinate and tyrosinate chelated PGRMC heme iron ion is unlikely to act as an electron carrier in enzyme catalyzed reactions (McGuire et al., 2021).

**Fig. 1.**
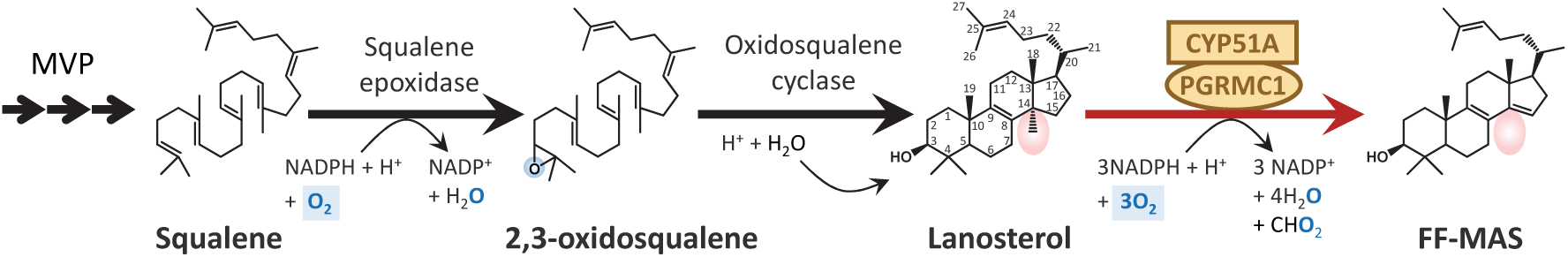
The production of FF-MAS from the mevalonate pathway. Enzymes involved in mammalian production and 14-demethylation of of the 4 methyl-sterol lanosterol. MVP: mevalonate pathway. The CYP51A/PGRMC1 reaction produces 14-dimethyl-14-dehydrolanosterol, also known as follicular fluid meiosis-activating sterol (FF-MAS). For CYP51A reaction stoichiometry see UniProt Q16850.

The isoprenoid precursors for lanosterol synthesis are produced by the mevalonate pathway, whose activity in eukaryotes is regulated by the SREBP/Insig/SCAP complex, which interacts with PGRMC1 (Cai et al., 2015; Suchanek et al., 2005). Thus, PGRMC1 is involved in the regulation of the mevalonate pathway, the modification of the first sterol, and it elicits responses to steroid/progesterone levels (Cahill, 2007; Cahill and Medlock, 2017; Neubauer et al., 2008).

PGRMC1 is also involved in endocytosis of low density lipoproteins (LDL) by the LDL receptor (Riad et al., 2020; Riad et al., 2018). Membrane traficking and sterol biology seem to have been ancient eukaryotic traits, whereas cholesterol transport via LDLs would be an evolutionary adaptation developed by multicellular animals. This both suggested that sterol biology is a central feature of PGRMC1 function, and reveals a gap in knowledge about how membrane trafficking and LDL receptor function arose.

We have recently shown that the evolutionary acquisition of phosphorylated PGRMC1 tyrosines 139 and 180 was coincident with the appearance of the gastrulation organizer and synapsed neurons in the common ancestor of eumetazoans (Hehenberger et al., 2020). Y139 is situated in the MIHIR, and represents one of the heptad repeat residues of a predicted coiled-coil protein interaction domain that shares similarity with coiled-coil motifs from various myosin proteins. Thereby, phosphorylation of Y139 would be predicted to disrupt coiled-coil interactions, and permit new interactions with phospho-tyrosine-binding SH2-domain proteins (Cahill, 2020; Hehenberger et al., 2020).

PGRMC1 is immunoprecipitated in protein complexes along with with components of the actin cytoskeleton (Salsano et al., 2020; Teakel et al., 2020), which is in turn involved in the membrane trafficking of exo- and endocytosis (Meunier and Gutierrez, 2016). It follows that evolutionarily ancient actin-cytoskeletal interactions of the PGRMC1 MIHIR sequence could be modulated by tyrosine phosphorylation that was acquired by eumetazoans at the same time as the gastrulation organizer developed (Cahill, 2020). Gastrulation involves induction by transcription factor Brachyury of a suite of actin-cytoskeletal genes that have been conserved since our common ancestor with the opisthokont *Capsaspora owczarzaki*, a single celled eukaryote that can switch between motile and sessile life cycle stages. A similar phenotypic switch is observed during early gastrulation when previously sessile epithelial cells undergo epithelial mesenchymal transition, and migrate within the embryo (Sebe-Pedros et al., 2016). Interestingly, PGRMC1 phosphorylation site mutants in cancer cells both altered the abundance of components of the actin cytoskeleton, and affected cell motility (Thejer et al., 2020a) by epigenetically altering gene expression (Thejer et al., 2020b).

PGRMC1 is induced in hypoxic human breast cancer cells at a time and place where cells switch to anaerobic metabolism (Neubauer et al., 2008), which was suggestive of PGRMC1 modulation of mitochondrial function. PGRMC1 localizes to mitochondria (Xu et al., 2011) where it associates with mitochondrial proteins (Salsano et al., 2020), modulates mitochondrial ferrochelatase (Piel et al., 2016), and is suggested to have co-evolved with a number of genes encoding mitochondrial proteins (Cahill and Medlock, 2017).

Experimental alteration of PGRMC1 phosphorylation status indeed dramatically affects mitochondrial form and function (Thejer et al., 2020a), simultaneously with altered genomic mutation rates and the status of genomic CpG epigenetic methylation (Thejer et al., 2020b). Since a MIHIR-containing MAPR gene appeared early in eukaryotes (Hehenberger et al., 2020), and PGRMC1 affects mitochondria, we set out to investigate a possible role of MAPR proteins in the origins of mitochondria and the eukaryotic cell.

Eukaryotic sterol biosynthesis arose primarily from a non-mitochondrially-inherited bacterial mevalonate pathway (MVP) (Castelle and Banfield, 2018; Hoshino and Gaucher, 2018). The isoprenoid squalene products of bacterial MVPs are cyclized by a squalene cyclase homolog into steroid-like six ringed structures called hopanoids (Barrantes and Fantini, 2016; Belin et al., 2018). Like mitochondrial cholesterol, hopanoids modulate bacterial membrane order and permeability to protons (Saenz et al., 2015). It has long been recognized that cholesterol decreases the permeability of the inner mitochondrial membrane to protons (Baggetto et al., 1992), thereby increasing the efficiency of electron transport chain activity, which is critical to mitochondrial function.

Eukaryotic lanosterol production involves the action of the novel enzyme squalene epoxidase, which produces 2,3-oxidosqualene that is then the substrate of squalene cyclase (Fig. 1). This eukaryotic-specific oxygen-consuming epoxidase reaction dictates that steroidogenesis is oxygen dependent (and hence may be regulated by oxygen tension), and is the reason that eukaryotic squalene cyclases produce the characteristic 3-hydroxysterols which bacterial hopanoids do not possess (Belin et al., 2018; Kirschvink and Kopp, 2008). As discussed above, the subsequent CYP51A/PGRMC1 catalyzed 14-demethylation of lanosterol also requires oxygen (Fig. 1). Hence, eukaryotic steroidogenesis can arguably be viewed as an adaptation of an ancestral hopanoid biosynthesis pathway which can modulate membrane lipid composition in the presence of atmospheric oxygen. This could potentially regulate a switch between aerobic and anaerobic metabolism.

The supply of cytoplasmically-synthesized sterols to the mitochondrion was probably important for eukaryogenesis before presumed proto-mitochondrial hopanoid pathway genes could be lost from the bacterial genome. With its mevalonate pathway and steroidogenic regulatory functions, sensitivity to sterol levels, and membrane trafficking functionality (Cahill and Medlock, 2017), a PGRMC1-like MAPR protein could have been ideally situated to enable this process. Here we investigated the origins of eukaryotic MAPR proteins, showing that the ancestral MAPR gene originated from a newly identified class of prokaryotic (cytb_5M_) cytb_5_-domain proteins. We were unable to conclusively identify the phylogenetic origins of the ancestral eukaryotic proto-MAPR gene, or whether a MAPR gene was present in the LECA.

## 2. Materials and methods

### 2.1. Selection of sequences

To search for MAPR-related proteins in bacteria for Fig. A1A, initial NCBI BLASTp analyses (https://blast.ncbi.nlm.nih.gov/Blast.cgi) (Johnson et al., 2008) were performed using the following MAPR proteins (Accession and FASTA files obtained from UNIPROT): Q9UMX5 (human NENF), Q8WUJ1 (human NEUFC), O00264 (human PGRMC1), O15173 (human PGRMC2), Q9XFM6 (*Arabidopsis thaliana* plant MAPR protein), and Q12091 (*Saccharomyces cerevisiae* yeast MAPR protein), or the single human cytrochrome b5 sequence (P00167). The NCBI non-redundant protein sequence data base was searched, restricting phylogentic taxa using the “Organism” input to the eukaryotes (taxid:2759), Bacteria (taxid:2) or Archaea (taxid:2157). Hits were selected as proteins with schematic graphical alignment across the cytb_5_ domain as provided by the BLASTp “Color key for alignment scores” output. Typical values were >40% identity, E<10. For Fig. A1A new eukaryotic sets of MAPR and cytb_5_ sequences were generated with BLASTp query sequences of NP_006658_1 (PGRMC1) and P04166 (rat cytb_5_). Three queries were made for each sequence, with “Organism” set to 1) Eukaryotes excluding Opisthokonta (taxid:33154) (the clade containing yeasts and animals), 2) Opisthokonta excluding metazoans (taxid:33208), and 3) Metazoans exluding Chordata (taxid:7711). For each query the top ten BLASTp hits were used to generate sets of 90 MAPR and cytb_5_ proteins. Default BLASTp parameters were used unless otherwise specified: Expect threshold “10”, Word size “6”, Max matches in a query range “0”, Matrix “BLOSUM62”; Gap Costs: “Existence: 11, Extension: 1”, and Conditional adjustments “compositional score matrix adjustment”.

Another round of sequence similarity searches was performed with the hope to capture a more diverse sequence dataset and ensure that all representatives of cytb_5MY_ were found. Separate BLASTp searches were performed against NCBI’s NR database using reference sequences for Clade 1 (KUO41884, NP_037481_1 and NP_006658_1), Clade 2 (P04166 and pdb_1CXY_A) and cytb_5MY_ (EKD84736, KKQ43725, KKR05718, KKR31495, KKS94036, KKU89055, OGE21801, OHA80798, OHA83705, OIP98122, and PJA14771). Each one of these BLASTp searches was performed using a maximum number of target sequences set to 10,000, and a maximum e-value of 1e-5. These sequences were grouped into a non-redundant fasta file of 2470 sequences. CD-Hit (Fu et al., 2012) was used to reduce redundancy of these sequences using 5 different similarity thresholds, namely 50%, 65%, 70%, 80% and 90%.

Reduced stringency BLASTp was performed with standard NCBI BLASTp settings except: 10,000 max sequences, expect threshold 50, Word size 3 (Organism: bacteria, taxid:2) or Word size 2 (Organism: archaea database, taxid:2157) (Fig. A3). Word size 3 and 2 are the available alternative options on the NCBI BLASTp server. Data sets were iteratively refined to yield multiple hundreds of aligned sequences with sufficient gap-free residues for NJ tree building as described case by case in Fig. A3.

For BLASTp analysis of cytb_5MY_-resembling proteins in eukaryotic sequences (Supplementary Appendix B) the accession numbers from Fig. A1C were individually entered as successive query sequences to the NCBI BLASTp server, with Organism restricted to Eukaryota (taxid:2759), otherwise employing default BLASTp settings. The top 100 best BLASTp hits per analysis were downloaded as FASTA files and subjected to MAFFT analysis as described below including control reference sequences NP_006658.1 (PGRMC1, clade-1), NP_037481.1 (NENF, clade-1), KUO41884.1 (Hadesarchaea archaeon YNP_N21 cytb_5M_, clade-1), P04166.2 (rat cytochrome b_5_, clade-2) and 1CXY_A (*E. Vacuolata* Cytochrome B558, clade-2). The 116 resulting sequences were subjected to MAFFT and refining of the data set by deleting gapped sequences to obtain sufficient gap-free alignments for tree building (between 112 and 116 sequences per analysis: see Supplementary Appendix B).

BLASTp of non-cytb_5MY_ candidate phylum radiation (CPR) bacterial cytb_5_-like proteins against eukaryotes were performed using query sequences (IMG identifiers) Ga0075854_11113, Ga0075854_11513, Ga0301032_10388 and Ga0301032_10385, with organism restricted to Eukaryota (taxid:2759) and all other parameter default values. BLASTp searches for cytb_5M_ proteins in CPR were performed separately using the query sequences KUO41884.1 and KKR30394.1 with Organism limited to bacteria candidate phyla (taxid:1783234). Resulting hits with homology over the cytb_5_-domain and E value <4 were downloaded. All hits from KKR30394 were also detected by KUO41884.

Taxon-restricted CYP51A sequences were obtained through BLASTp searches using UniProt Q16850 (human CYP51A) as search string with organism restricted in separate searches to 1) Opisthokonta (taxid:33154) exclude: Bilateria (taxid:33213) (“Eukaryotic”), 2) Archaebacteria (taxid:2157), 3) Alphaproteobacteria (taxid:28211) (“alphaproteobacteria”), and 4) Bacteria (taxid:2), exclude Alphaproteobacteria (taxid:28211) (“other bacteria”). Additionally, the eukaryotic sequences obtained by this search were aligned using MAFFT L-INS-I and the obtained alignment was used as query for a taxon-unrestricted Psiblast sequence similarity search (Altschul et al., 1997) against NR, with 100,000 maximum target sequences and an e-value of 1e-5. Out of these sequences, 4859 with e-values lower than 1e-50 were selected, and reduced with CD-Hit using a 70% similarity threshold, resulting in 533 sequences.

### 2.2. Phylogenetic analyses

#### 2.2.1. Phylogenetic reconstruction of cytb5 sequences

All NCBI BLASTp hit CSV files were downloaded and hits combined in Microsoft Excel. Non-redundant FASTA sequences were retrieved using Batch Entez, and subjected to MAFFT alignment by the L-INS-i method (Katoh et al., 2019; Kuraku et al., 2013; Yamada et al., 2016) using the Computational Biology Research Consortium (CBRC) server at https://mafft.cbrc.jp/alignment/server/index.html. For reduced stringency BLASTp, sequences which branched outside of MAPR (clade-1) and rat cytb_5_ (clade-2) reference sequences on the preliminary guide tree were discarded prior to iterative elimination of aligned sequences and NJ tree-building. Sequences with gaps were deleted manually by inspecting the MAFFT FASTA alignment in AliView (Larsson, 2014) and removing gapped sequences via the “Refine dataset” function of the CBRC server to iteratively generate sequence data sets with sufficient gap free sites for tree-building. The sequence selection was inverted, followed by sequence realignment with MAFFT L-INS-i until sufficient gap-free sites for tree building were iteratively obtained. Inferred phylogenetic trees were constructed using the NJ method (all gap-free sites) and each of JTT, WAG and Poisson substitution models with bootstrap selected and 1,000 resamplings, and were processed using Archaeopteryx software through the forester.jar program (Han and Zmasek, 2009), as described on the CBRC server.

Another round of phylogenetic analyses was performed on the extended set of homologs for cytb5 sequences, and the 5 datasets generated through sequence redundancy reduction with CD-Hit (see above). The resulting datasets were each aligned using MAFFT L-INS-I and T-Coffee’s regressive algorithm (Garriga et al., 2019), using the heads-and-tails strategy whereby each alignment is performed in the forward and reverse sense of the sequences, and the four resulting alignments were then merged using MergeAlign (Collingridge and Kelly, 2012). Finally, the merged alignments were trimmed using the - gappyout option in trimAl (Capella-Gutierrez et al., 2009). Phylogenetic reconstruction was then performed with IQ-Tree 2.0 (Minh et al., 2020) using ModelFinder (Kalyaanamoorthy et al., 2017) with all variations of the models JTT, WAG, and LG, plus the mixture models generated by LG+C10.C60, with multiple rate heterogeneity (none, +G4, +R2, +R4, +R8) and frequency (none, +F) parameters, and adding the models LG4M and LG4X. Finally, PMSF (Wang et al., 2018) approximations of the chosen models (LG+C60+F+R8 for the 50% and 65% datasets, LG+C50+F+R8 for the 70% and 90% datasets, and LG+C40+F+R8 for the 80% dataset) were performed for all alignments to obtain trees with 100 non-parametric bootstrap pseudorreplicates, which were then analysed both using the classical Felsenstein criterion, and the Transfer Bootstrap Expectation (TBE) (Lemoine et al., 2018) variant. One more round was performed on sequence belonging only to Clade 1. The same procedure as above was followed, but more parameter variations were explored. Alignments were obtained with MAFFT L-INS-I. Trimming was performed to remove alignment positions with either over 75% or 50% gaps. Phylogenetic reconstructions were performed using LG+C60+R4+F+PMSF, WAG+C60+R4+F+PMSF or the model chosen by ModelFinder among JTT, WAG and LG with C10, C20, C40 and C60 mixtures, plus LG4M, LG4X, UL2 and UL3, in all combinations with extended rate heterogeneity (none, +G4, +G8, +R4, +R8) and frequency (none, +F) parameters), resulting in the WAG+C60+R6 and WAG+C60+R6+F mixture models being chosen for the PMSF approximation of the 50% and 65% reduced datasets, respectively). The 65%, 70% and 90% reduced datasets were also aligned following the heads-and-tails strategy with MAFFT L-INS-I, trimmed at a 75% gap threshold, and used for phylogenetic reconstruction under the LG+C60+R4+F+PMSF model. Critical TBE thresholds were calculated for specific branches following Lemoine et al. (2019), as (1 – ((p-x)/(p-1)), where t is the number of sequences at the lighter side of the bipartition, and x is the number of sequences of the clade of interest (13).

#### 2.2.2. Phylogenetic analyses of CYP51A

The 533 CYP51A sequences obtained as above were aligned using MAFFT L-INS-I and trimmed with trimAl removing all columns containing at least 50% gaps. IQ-Tree 2.0 was used for phylogenetic reconstruction, under the model LG+G4+F, with 1000 replicates for SH-like approximate likelihood ratio tests and Ultrafast bootstraps.

### 2.3. Crystal structure

To achieve protein expression, the codon optimised ORF was subcloned into pGEX-4T-1-H expression vector to create pGEX4T1_KUO41884 (Fig. A2), transformed into *BL21(DE3) pLysS* cells (Novagen) (9F-, *ompT, hsdS*B(rB-mB-), *gal, dcm* (DE3) pLysS (CamR)), and cultured in an expression base media (1 % Tryptone, 0.5 % yeast extract) containing 30 mL of 2.852 M NaCl and ampicillin (100 μg/ml) until the OD_600_ reached 0.6. Protein expression was induced using 1 mM isopropyl 1-thio-D galactopyranoside (IPTG) and cells were incubated overnight at 37°C at 80 rpm. Cells were harvested by centrifugation at 6,000 rpm at 18°C for 30 minutes. Cell pellets were resuspended in GST (glutathione-S-transferase) cell lysis buffer (50 mM Tris(hydroxymethyl)aminomethane, 125 mM NaCl, pH 7.4). All protein purification steps were performed at room temperature. The soluble whole cell lysate was filtered through a 0.45 μM syringe filter. The soluble cell extracts were injected using a superloop at 2 mL/minute into a GST column equilibrated with GST cell lysis buffer. Proteins were eluted using 10 mM Glutathione. The N-terminal GST tag was cleaved with 100 μL of Tobacco etch virus (TEV) protease. Size exclusion chromatography was performed to further purify proteins using AKTA FPLC using an S200 26/600 filtration column (GE Healthcare), following Khandokar et al. (2017). The protein eluted as a single homogeneous peak, and at a volume consistent with the expected molecular weight of a monomer. The protein was stored at -20°C prior to use for crystallisation.

Crystals were obtained through sparse matrix screening and the hanging drop vapor diffusion method following Khandokar et al. (2017). Crystals in the space group *P*6_3_ diffracted to 1.9Å at the Australian Synchrotron microcrystallography beamlines (Aragao et al., 2018; Cowieson et al., 2015). The diffraction data was integrated in Mosflm (Battye et al., 2011), scaled, and reduced in AIMLESS (Evans, 2011; Evans and Murshudov, 2013), and the structure determined by molecular replacement using PDB ID 1J03 in Phaser (McCoy et al., 2007), REFMAC (Vagin et al., 2004), PHENIX (Adams et al., 2010), and COOT (Emsley et al., 2010). The final structural model has been refined to an Rwork and Rfree of 0.22 and 0.27 respectively, no Ramachandran outliers, and good stereochemistry (see Table A2). LIGPLOTs (Laskowski and Swindells, 2011), structural superposition (Krissinel and Henrick, 2004), and DALI (heuristic PDB search) protein structure comparison by alignment of distance matrices (Holm and Laakso, 2016) were performed as described.

### 2.4. Other protein analysis

The degree of conservation of cytb_5_-domain residues in the structure of PGRMC1 was visualized using the Consurf server (http://consurf.tau.ac.il/) (Ashkenazy et al., 2016). Sequence Logo Plots of amino acids in respective protein clades were generated using WebLogo (http://weblogo.berkeley.edu/logo.cgi) (Crooks et al., 2004). Non-PGRMC1 residues from the 4X8Y structure (Kabe et al., 2016a) were not included in the PGRMC1 structural data for Fig. 3.

### 2.5. Gene cluster analyses

Conserved gene cluster analysis is based on the observation that functionally related genes are often collocated on the chromosomes in prokaryotes, preserving similar gene context across phylogenetically diverse organisms (Overbeek et al., 1999; Pellegrini et al., 1999). During evolution proteins that function together in a pathway or structural complex evolve in a correlated fashion, and tend to be either preserved collectively, thus ensuring that the pathway or complex remains fully functional, or be eliminated all together. Methods of chromosomal gene context analysis have proved to be valuable for delineation of evolutionary patterns between organisms, as well as for protein function prediction (Mavromatis et al., 2009; Overbeek et al., 1999; Pellegrini et al., 1999).

To find all genomes containing cytb_5MY_ genes, we first downloaded all 7645 CPR bacterial genomes from NCBI. Using the alignment employed to reconstruct the phylogeny in Fig A3B, we performed an HMM search against all CPR genomes using HMMER 3.1b2 (hmmer.org). All candidate sequences were joined with those from the original alignment, aligned with MAFFT-auto, trimmed with trimAl to remove all columns with over 90% gaps and used for fast phylogenetic reconstruction using Fasttree2 (Price et al., 2010). Genomes containing genes clustering with the cytb_5MY_ sequences were selected for further analysis, and annotated using InterProScan v5.48-83.0 (Jones et al., 2014). Taxonomic identification of these genomes was performed against the dereplicated genomes provided by Jaffe et al. (2020) using fastANI (Jain et al., 2018) with default parameters or, if no results were found, by using a fragment length threshold of 1000 bp. Gene synteny analyses were performed using the genoPlotR package (Guy et al., 2010) and the Gene Neighborhood Viewer and Chromosomal Cassette Viewer tools (Mavromatis et al., 2009) of the Integrated Microbial Genomes (IMG) database (Chen et al., 2019).

## 3. Results

### 3.1. MAPR related to new prokaryotic cytb_5M_

Preliminary BLASTp searches for the presence of MAPR-related proteins in prokaryotes were conducted using eukaryotic MAPR proteins and human cytb_5_ as queries. We combined the significant hits from separate BLAST searches into a non-redundant list of 176 proteins, and generated an alignment including the reference sequences for human PGRMC1, human NENF, and rat cytb_5_. We inferred a preliminary distance-based phylogenetic tree, which produced two distinct eukaryotic clades (Fig. A1A), both of which were subtended by prokaryotic sequences. Notably, clade-1 contained MAPR proteins and the BLASTp hits obtained with those queries, and clade-2 contained eukaryotic cytb_5_, and its corresponding BLASTp hits. These results suggest that the eukaryotic cytb5 protein families are polyphyletic and originated from at least two separate prokaryotic lineages. An alternative explanation could be that we have theoretically sampled MAPR-like and cytb_5_-like proteins from a single bacterial homolog, with apparent distinctiveness being an artefact of our BLASTp search strategy using MAPR and eukaryotic cytb_5_ sequence search strings.

To address this possibility we first separately interrogated the archaeal and bacterial sequence data bases by BLASTp with selected prokaryotic sequences from each of clades 1 and 2 from the prior analysis to generate independent sets of cytb_5_ sequences. MAFFT alignment again produced inferred trees with clade-1 and -2 tree topology, with bootstrap confidence values (BCV) >94% for both archaeal and bacterial trees (see Supplementary Appendix A). BLASTp with string sequences from one clade again did not detect members of the other. Combining all of the sequence from above with additional MAPR sequences yielded similar results when aligning 814 sequences (Fig. A1B), consistent with the cytb_5_–domain proteins in clades-1 representing a new type of distinct prokaryotic cytb_5_ proteins. MAPR and some bacterial proteins formed a sub-cluster distinct from 348 other prokaryotic clade-1 sequences with BCV 87% (Fig. A1B). PDF tree images, FASTA sequence alignments, and xml tree files for preliminary analyses and the panel of Fig. A1A are available in Supplementary Appendix A.

We had still not eliminated the possibility that separate clades were due to BLASTp search string bias. To further investigate this issue, we employed reduced-stringency BLASTp analysis by using ‘word size’ (the number of adjacent residue identity required for alignment) parameters of 3 and 2, so that queries from one clade detected sequences from the other clade (Fig. A2). The results continued to generate two discrete clades connected by a long branch (Fig. A3). We therefore detected no evidence that the two eukaryotic clades detected represented a sampling artefact selected by a biased choice of BLAST query sequences from a broad continuum of prokaryotic cytb_5_ proteins. . Instead, these results support the existence of two separate types of prokaryotic cytb_5_ domain, which gave rise to eukaryotic MAPR proteins (clade-1) and cytb_5_ proteins (clade-2), respectively. We propose the name cytochrome b_5<u>M</u>_ (cytb_5M_) for the newly recognized “<u>M</u>APR-like” prokaryotic cytb_5_-domain clade-1 proteins.

### 3.2. Bacterial cytb_5MY_ cluster with MAPR

Next, we considered the clade-1 bacterial sequences that clustered closer to MAPR than to other prokaryotic cytb_5M_ proteins (Fig. A1B,C). Three contained MIHIR residues, while a clade of 11 sequences did not. The first three belonged to two 65 and 14 kb-long delta-proteobacterial contigs (KPK16282 and KPK52515) and one acidobacterium chromosome (ANM31058). They were nested within eukaryotic sequences in the tree, thus potentially representing misclassified sequences or horizontally transferred genes. The top eukaryotic BLASTp hits for two MIHIR-containing sequences were plants (KPK16282.1, KPK52515.1; not shown), and for the other (ANM31058.1) were all either choanoflagellates or animals (not shown). We investigated the top BLASTp similarity of the proteins encoded by the two contiguous genes in their respective contigs, and observed that multiple top hits belonged to the same broad taxa, consistent with their classification being correct. Hence, these MIHIR-containing bacterial sequences likely represent HGT originating from eukaryotic plants and holozoans.

The other 11 MAPR-clustering bacterial sequences lacked MIHIR sequences, but contained the cognate equivalents of PGRMC1 heme-chelating Y113, and heme hydrogen bond donors Y107, K163 and K164 (Kabe et al., 2016a) (Fig. A1C), as opposed to the cytb_5_–domain bis-his axial heme ligation of hitherto described bacterial and eukaryotic cytb_5_ proteins (Kabe et al., 2016a; Liu et al., 2014). We designate this subgroup of cytb_5M_ proteins with “<u>M</u>APR-like tyrosines (<u>Y</u>)” as cytb_5<u>MY</u>_. All cytb_5MY_ sequences were found to originate from CPR bacteria (Table A1). The top 100 BLASTp hits of all cytb_5MY_ sequences from Fig. A1B against eukaryotic sequences returned only proteins with MIHIR sequences (Supplementary Appendix B), indicating that eukaryotic genomes do not encode cytb_5MY_ proteins.

To further investigate cytb_5MY_ phylogenetic relationships, we performed additional sequence similarity searches using the 11 cytb_5MY_ and additional reference sequences as queries (see Methods). We then reduced the resulting 2486 sequences under 5 different levels of redundancy, and subjected each one of these into more rigorous phylogenetic analyses. The obtained maximum likelihood phylogenies corroborated the previous topology, revealing two major separate eukaryotic clades (containing MAPR and cytb_5_ sequences, respectively), and supporting the affiliation of CPR-bacterial cytb_5MY_ with the eukaryotic MAPR sequences, albeit under different topologies and without satisfying branch support (Fig. A4). This is important because if cytb_5MY_ formed a sister group to eukaryotic MAPR proteins it could imply that MAPR proteins originated from a cytb_5MY_ protein. However, if cytb_5MY_ proteins are topologically nested within the MAPR family, it would imply that CPR bacteria obtained a MAPR gene from eukaryotes by HGT. We were unable to discriminate between these models.

In four of the obtained phylogenies, cytb_5MY_ branched within eukaryotes, while the fifth displayed cytb_5MY_ as sister to eukaryotes. For each phylogenetic reconstruction, 100 non-parametric bootstrap pseudorreplicates were obtained, for which both Felsenstein Bootstrap Proportion (FBP) and Transfer Bootstrap Expectation (TBE) (Lemoine et al., 2018) values were calculated to assess the robustness of the obtained trees.

We performed additional phylogenetic reconstructions using the same 5 reduced datasets, but restricting the analysis to clade-1 sequences only (Fig. 1, Supplementary Appendix D). In these trees, we observed two major topologies with respect to the positioning of the cytb_5MY_ sequences: as sister to MAPR group (Fig. 1A), or nested within it, possibly as sister to the NEUFC clade (Fig. 1B) (see also Supplementary Appendix D). In general, these trees included high TBE support for many relevant branches, and none of the trees can be considered to accurately reflect cytb_5MY_ relationships. TBE values reflect the average number of transfers at either side of the bipartition, thus indicating the consistency of a clade but not necessarily of all its members (Lemoine et al. 2018). For that reason, we calculated for each tree the threshold TBE value that would support a given bipartition without allowing the transfer of the 13 cytb_5MY_ sequences.

With this consideration, several trees on the sequence datasets reduced at 50% and 90% redundancy levels included a TBE-supported topology with cytb_5MY_ as sister to all eukaryotic MAPR sequences. However, trees reconstructed from the sequence dataset reduced at 65% redundancy level supported the alternative, derived position as sister to the NEUFC eukaryotic sequences. From these results we conclude that cytb_5MY_ may derive from a HGT event from a eukaryote into a CPR bacterium, followed by loss of the MIHIR sequences and further propagation to other CPR species through HGT. However, an alternative scenario that a CPR cytb_5MY_ protein gave rise to eukaryotic MAPR proteins cannot be excluded by our tree-building exercise.

Furthermore, the relationship between the eukaryotic MAPR protein families indicates that multiple events of duplication occurred early in the evolution of eukaryotes and diversified in various eukaryotic groups (Hehenberger et al., 2020). However, it is not possible to conclude from this analysis whether a MAPR ancestral protein was present in the LECA, or whether it was acquired later during eukaryotic evolution. The analysis is limited by the relatively short homologous region of the cytb5 domain, the substantial evolutionary distances separating the sequences, and the skewed taxonomic distribution of the domain.

Both MAPR and cytb_5MY_ proteins are derived cytb_5M_ proteins. However, the primitive MAPR/cytb_5MY_ structural and functional characteristics remain uncertain. We suggest these proteins to be sufficiently distinct to warrant reference to prokaryotic cytb_5M_ and cytb_5MY_ proteins, with the continued use of the MAPR terminology for eukaryotes (although formally, as clade-1 members, MAPR and cytb_5MY_ proteins cladistically belong to the cytb_5M_ clade).

### 3.3. Sequence differences between clades

Next, we analyzed the nature of conserved sequence differences between cytb_5_, cytb_5M_, cytb_5MY_, and MAPR proteins. Sequence logos (Fig. 3A) revealed altered frequencies of amino acid usage between clades (ΔC1:ΔC2, Fig. 3A). This included a surface loop between PGRMC1 G83-R88 (Kabe et al., 2016a) (loop-1 in Fig. 3), which is at least two residues larger in all clade-1 than clade-2 proteins. G83, site of a pronounced change of polypeptide backbone direction (Kabe et al., 2016a), is strongly conserved in clade-1 (Fig. 3A). A second loop involving rat-cytb_5_ heme-binding (Rodriguez-Maranon et al., 1996) was strongly conserved in clade-2 (loop-2 in Fig. 3). Additionally, relative to cytb_5M_, MAPR and cytb_5MY_ proteins exhibit conserved cognates of PGRMC1 Y107, Y113, K163 and Y164 (Fig. A1C), all of which are involved in heme interaction (Kabe et al., 2016a). The residues differential between clade-1 and clade-2, or cytb_5M_ and cytb_5MY_/MAPR proteins are interspersed along the primary PGRMC1 sequence (Fig. 3A, top), yet form a contiguous surface extending from the heme-binding pocket to the surface loop, which rests upon conserved F81 and G83 (Fig. 3B,C). Residues more similar between MAPR and cytb_5MY_ are clustered around the heme-binding pocket (purple in Fig. 3A,B). In contrast, the residues conserved among all proteins in the analysis (conserved between both cytb_5_-domain clades) predominantly occupy the protein interior (Fig. 3D).

### 3.4. Clade-1 protein structure

We next solved the first crystal structure of a prokaryotic cytb_5M_ clade-1 protein from the archaeon Hadesarchaea YNP_N21 (KUO41884.1) (Fig. A1B) (attempts to crystalize multiple cytb_5MY_ proteins were unsuccessful: not shown), to compare this with the structures of representative proteins from each major group of clades-1 and -2, revealing overall shared similarity of the clade-1 proteins (Fig. 4). The HP and HS heme iron chelation sites of rat cytb_5_ (Fig. 4E) are strongly conserved in clade-2, with loop-2 extending from the conserved HP site (Fig. 3A). Hadesarchaea-cytb_5M_ lacks the MAPR heme-binding tyrosines yet binds heme in a MAPR-like orientation. It shares one heme chelating histidine (H61) with conventional (clade-2) cytb_5_ proteins (Fig. 4E, Fig. A5), which is conserved in cytb_5M_ proteins (Fig. 3A). Heme-binding residues differ between cytb_5M_, cytb_5MY_ and MAPR proteins within clade-1 (Fig. 3A). Excluding known MAPR proteins (Cahill, 2007), the published structures most closely resembling Hadesarchaea-cytb_5M_ belong to clade-2 and exhibit clade-2-like heme-binding (Fig. A6, Table A3), confirming the novelty of our archetypal clade-1 cytb_5M_ structure. Altogether, the requirements for dissimilar heme-binding and folded architecture underlie the presence of two discrete prokaryotic cytb_5_ domain-related clades in Fig. 2, one of which gave rise in eukaryotes to classical cytb_5_ (clade-2), and the other to MAPR proteins (clade-1).

**Fig. 2.**
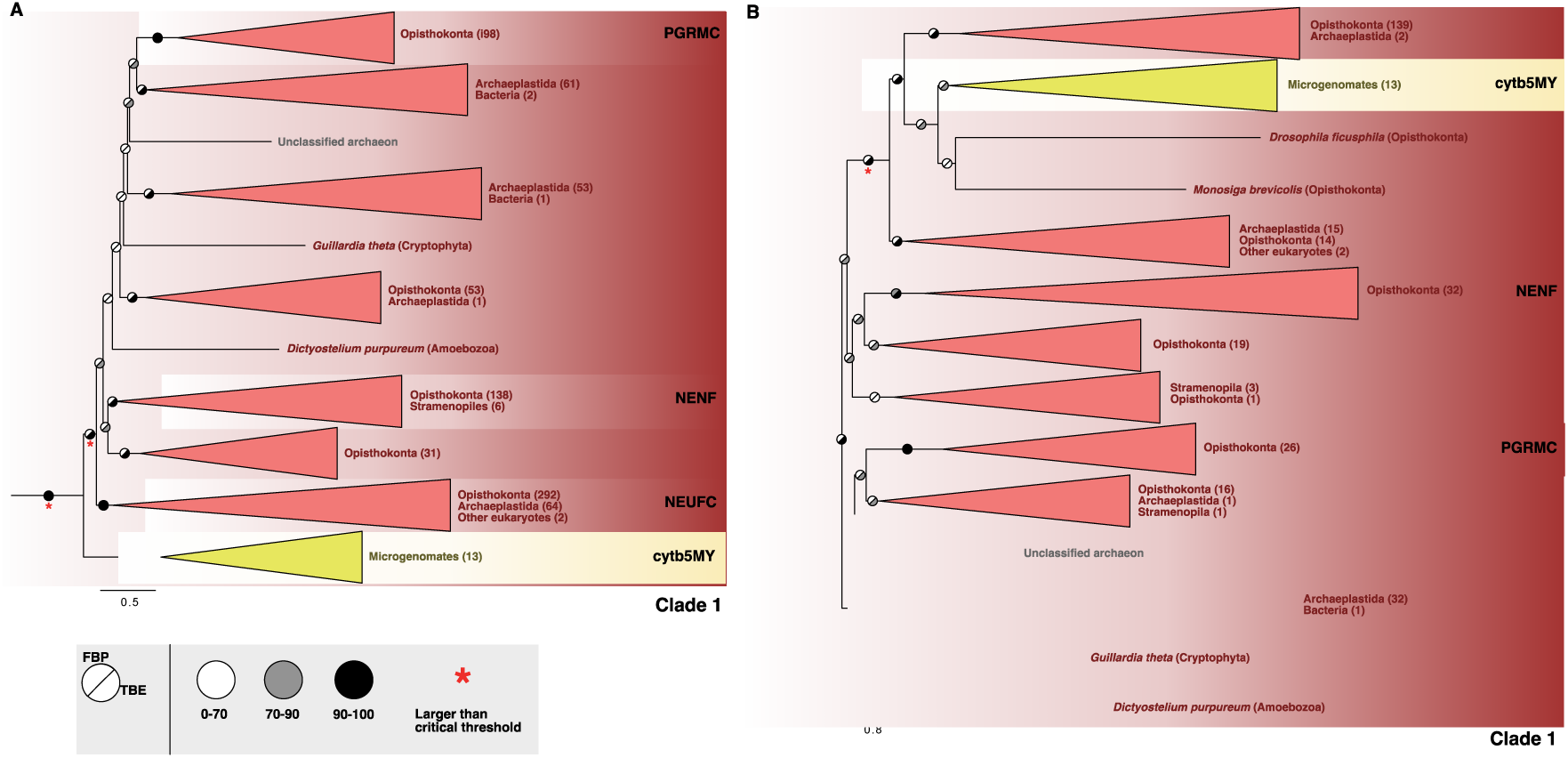
Eukaryotic cytochrome b5 domain proteins are polyphyletic. Maximum-likelihood phylogenetic reconstruction using IQ-Tree2 under LG+C60+R4+F+PMSF model of cytb5 sequences reduced at 90% (A) and 65% (B) redundancy. Both trees were rooted on the branch leading to Clade 1 bacterial sequences. Branch symbols indicate Felsenstein Bootstrap Proportions (FBP; upper left half) and Transfer Bootstrap Expectation (TBE; lower right half) interpretations of 100 non-parametric bootstrap pseudorreplicates. Critical thresholds for TBE were calculated as explained in the methods. Additional phylogenetic reconstructions under these and other levels of redundancy, alternative alignment and trimming strategies, and alternative evolutionary models, are all summarized in Supplementary Appendix D. Sequences, alignments and trees are included in Supplementary Appendix C.

**Fig. 3.**
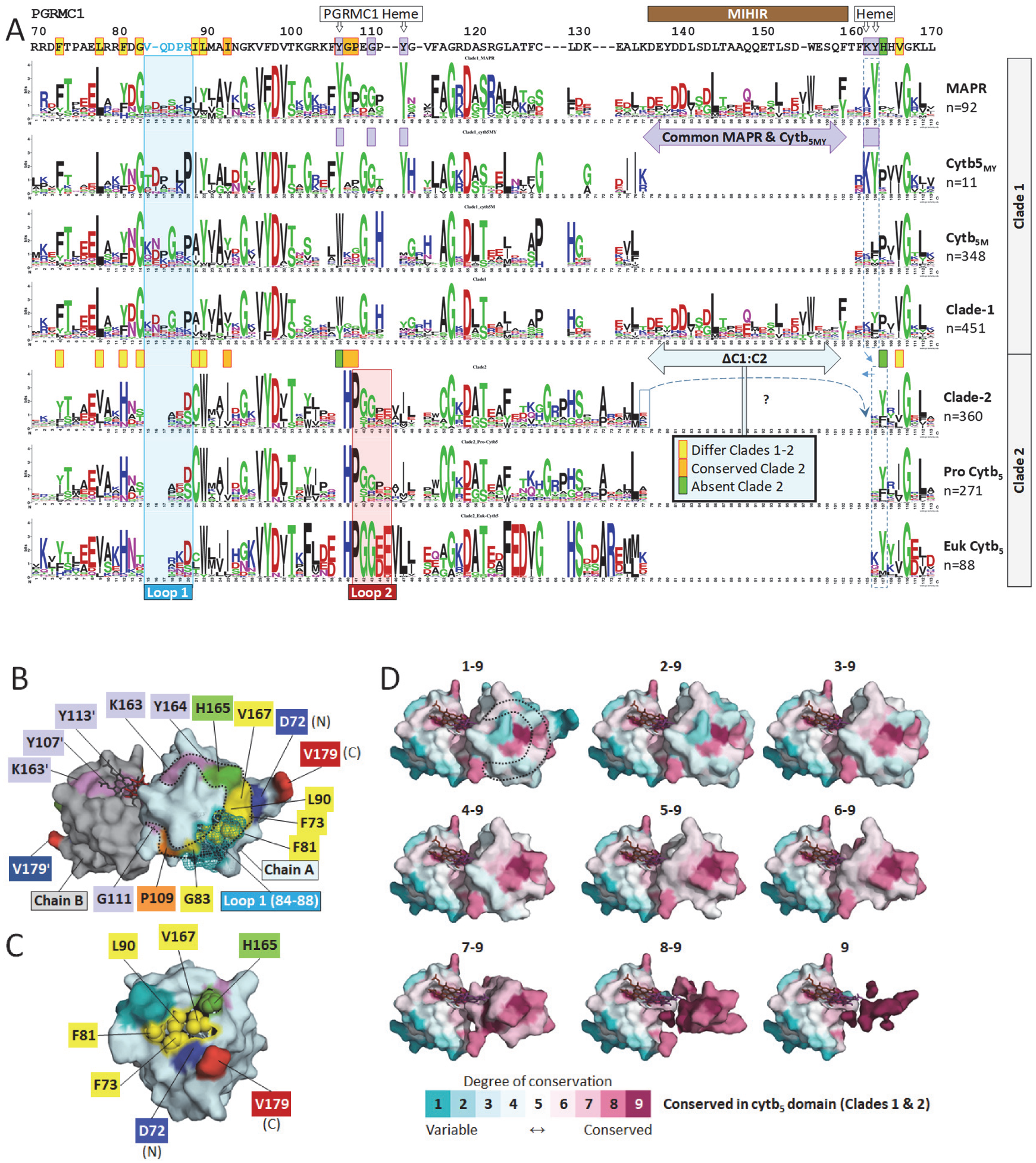
A conserved clade-1 surface. (A) Sequence logos of the alignment of Fig. A1A. The box and “?” in clade-2 show a potentially incorrect inter-clade alignment, in which case the adjacent position (H165) is absent from clade-2. “Loop-1” and “loop-2” are shown. (B) A conserved clade-1 surface. Loop 1 is shown as open mesh to display conserved underlying F81 and G83. (C) Rotation through 90° relative to “B”, showing conserved exposed surface residues as filled spheres. (D) Residues conserved between all cytb_5_-domain sequences in Fig. A1A (both clades) constitute the protein interior, as generated by Consurf (Ashkenazy et al., 2016), based upon 4X8Y.

**Fig. 4.**
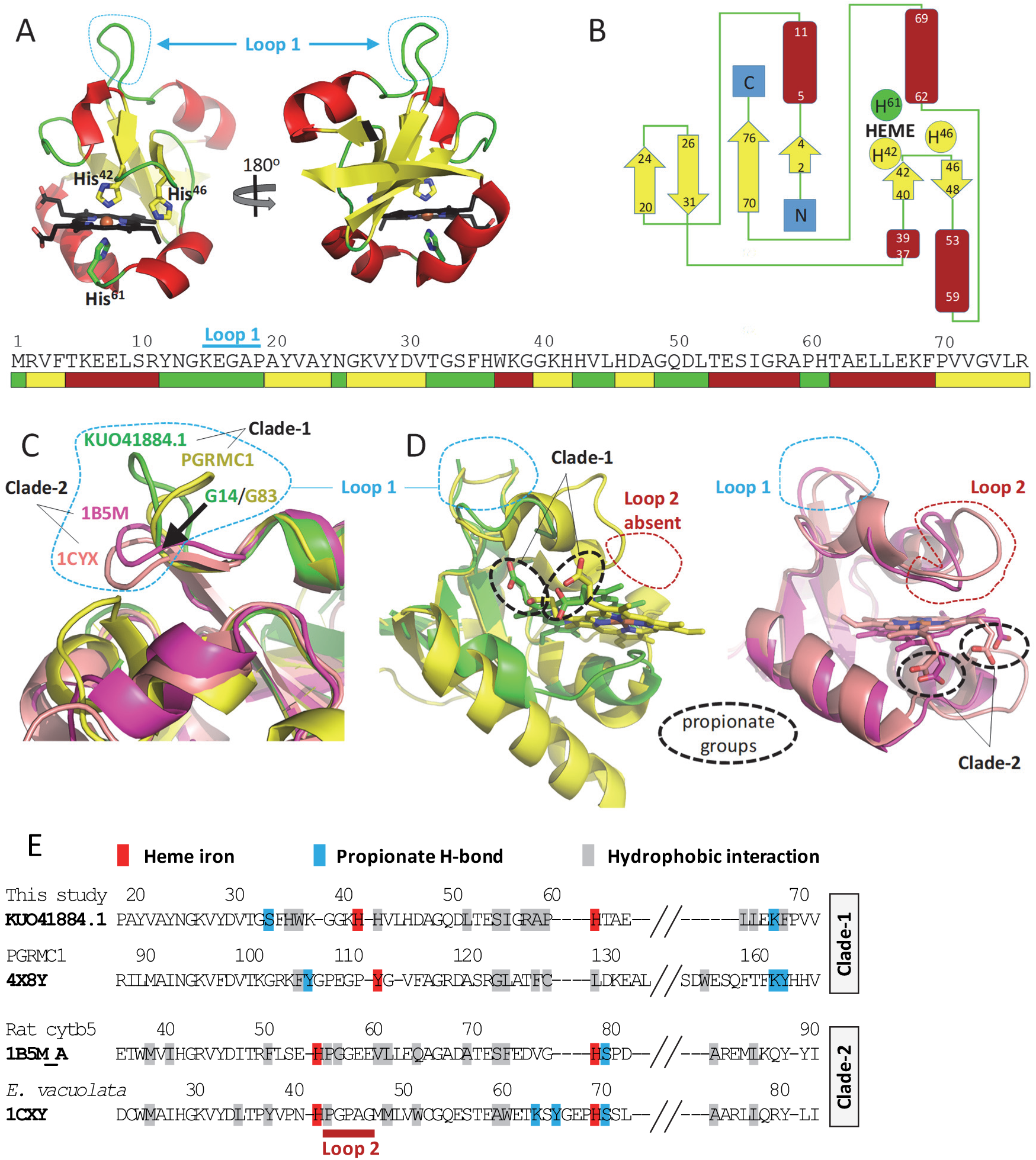
The crystal structure of Hadesarchaea cytb_5M_. (A) Hadesarchaea YNP_N21 (NCBI KUO41884.1) cytb_5M_ structure. (B) Topological organization of the structure from A). Numbering follows KUO41884.1. (C) Alignment of loop-1 regions of bacterial cytb_5_ (1CXY, *Ectothiorhodospira vacuolata*) (Kostanjevecki et al., 1999), rat mitochondrial cytb_5_ (1B5M_A) (Rodriguez-Maranon et al., 1996), human PGRMC1 (4X8Y) (Kabe et al., 2016a), and KUO41884.1. The position of KUO41884.1 G14 and PGRMC1 G83 is arrowed at the base of the clade-1-specific loop (PGRMC1 84-88). (D) Heme orientation differs by typically 90 degrees between clades-1 and -2. (E) Contacts with heme. The four proteins are shown in the alignment from Fig. A1A, showing heme-interacting residues. See Fig. A5 for interaction details.

### 3.5. Genomic context of CPR cytb_5MY_ clade-1 proteins

To compare the genomic context of CPR cytb_5M_ and cytb_5MY_ genes we investigated the distribution of both protein types in CPR bacterial genomes. To recover all genomes containing cytb5_MY_ sequences regardless of sequence redundancy, we performed another phylogenetic analysis with the sequences shown in Fig. 1AB and all hits with e-value lower than 1e-5 after an HMM search against all 7645 CPR genomes found in NCBI (as of July 5^th^ 2021) (Fig A6). This tree revealed a total of 36 genomes which contained cytb5_MY_ proteins. To visualise the presence of cytb_5MY_ in CPR bacteria, we mapped these 36 genomes onto a phylogenomic reconstruction performed by (Jaffe et al., 2020) (Fig. 5A). This analysis revealed that these genomes belong to multiple distinct phyla and that the presence of cytb_5MY_ is highly punctuated in CPR bacteria, indicative of horizontal gene transfer as their main mode of evolution.

**Fig. 5.**
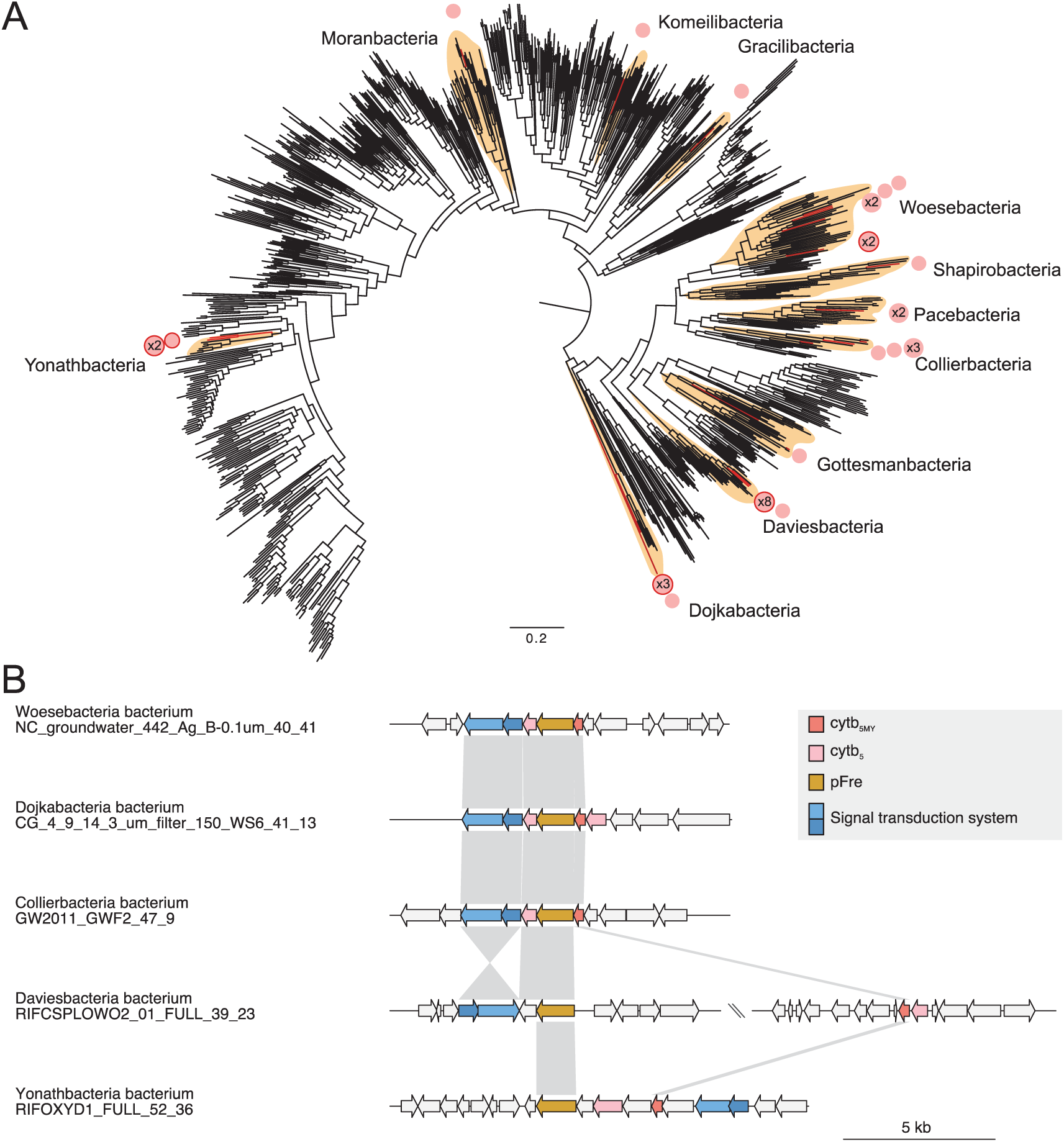
Phylogenetic distribution and CPR genomic context of cytb_5MY_ genes. (A) Phyletic distribution of cytb_5MY_, mapped onto a maximum likelihood tree obtained from a concatenated alignment of 16 ribosomal proteins by Jaffe et al. (2020). Genome assignment was performed through ANI value calculation: circles indicate assignment with an ANI value larger than 70%; circles are outlined if they indicate an assignment larger than 99%), and larger if they indicate assignment of more than one genome. Only the taxonomy of groups with cytb_5MY_-containing genomes is indicated. (B) 5-kb gene neighbourhoods of cytb_5MY_ and pFre genes of selected genomes. Each arrow represents a gene and they are colored based on assigned function according to the legend. Connecting lines are drawn between homologous genes found as best reciprocal Blastp hits with e-values under 1e-5.

We then investigated the gene neighbourhoods of cytb_5MY_ in these 36 genomes and found that in most (29), the cytb_5MY_ genes colocalized with putative ferric reductase (pFre) genes containing a ferric reductase-like transmembrane domain (Fig. 5B, Fig. A7). Other genes found to colocalise with cytb_5MY_ were those containing other cytb5-domain proteins (28), and two-component inducible signal transduction systems (22) (Fig. 5B, Fig. A7). The predicted phylogenetic tree topology for the cytb5MY genes in Fig. A7 is identical to Fig. A8, which expands the cytb5MY section of Fig A3B. While the synteny was often lost, a putative operonic structure of these genes followinig the same disposition was found for multiple distant lineages (e.g., Fig. 5B). Alternative configurations were found in multiple genomes, although these were often highly similar, such as a group of 8 Daviesbacterial and 3 Yonathbacterial genomes with over 99% average nucleotide identity (ANI) values, respectively (Fig. 5A).

Using the Integrated Microbial Genomes and Microbiomes (IMG) platform, we observed that besides cytb_5MY_ and pFre, cytb_5MY_ -containing operons often involved two novel types of atypical cytb_5_-domain-like proteins which we denote as cytb_5_-like type A (CBLA) and B (CBLB) proteins, additional to the cytb_5MY_ protein and a two-component signal transduction element (Fig. 5B). The CBLA and CBLB proteins were distantly related to cytb_5_-domain protein clades-1 and -2, respectively, with barely detectable homology (Fig. A9). Because the deepest branches of the tree of Fig. A7 contain a two component element, a pFre gene, two cytb5-related genes as well as a cytb5MY gene, this probably represents the ancestral CPR state. All proteins in this putative ancestral operon contain one or more predicted transmembrane helices – hence, we speculate these could potentially form a membrane-associated protein complex.

Pfamscan (Mistry et al., 2007; Mistry et al., 2021) identified the ferric reductase domain (PF01794) in 1427 of the 7645 CPR genomes, 185 of which contained genes with cytb_5_ domain (PF00173) within 10 kb of the pFre genes (164 of which were distinct from the 39 genomes described above). Yet colocalization of pFre with cytb_5_-domain genes, including cytb_5MY_, was not observed outside of CPR bacteria in the IMG database, suggesting that a putative transmembrane protein complex containing pFre and cytb_5_-like proteins performs an unknown CPR-specific function.

### 3.6. CYP51A is of bacterial origin

Having considered the prokaryotic origins of MAPR proteins, we next considered the origins of the eukaryotic MAPR-containing enzyme pathway responsible for synthesizing 14-demethylated lanosterol. Fig. 1 shows the conversion of squalene, the product of the mevalonate pathway (MVP), to the first sterol lanosterol, and subsequent decarboxylation to 14-dimethyl-14-dehydrolanosterol, also known as follicular fluid meiosis-activating sterol (FF-MAS) (Mitsche et al., 2015). The respective MVP (Castelle and Banfield, 2018; Hoshino and Gaucher, 2018) and squalene cyclase enzymes (Barrantes and Fantini, 2016; Rajamani and Gao, 2003) are of bacterial origin (Frickey and Kannenberg, 2009). PGRMC1-like proteins bind to and regulate CYP51A to catalyze subsequent lanosterol-14-demethylation in opisthokonts such as yeast and mammals (Hand et al., 2003; Hughes et al., 2007b). The evolutionary origin of CYP51A, however, has thus far remained unclear.

We performed taxon-restricted maximum-likelihood phylogenies of CYP51A, which revealed a closer relationship between eukaryotic CYP51A and bacterial rather than archaeal proteins with an ultrafast bootstrap support of 91% and an SH-like approximate likelihood ratio test support value of 98.7% (Fig. 6). A larger-scale taxonomically-unrestricted sequence similarity search against the NR database retrieved ca. 3500 eukaryotic, ca. 1000 bacterial (of which 30 from Alphaproteobacteria) and only 4 archaeal sequences with an e-value lower than 1e-50. A maximum-likelihood phylogeny of a reduced but phylogenetically diverse dataset obtained from these sequences resulted in a monophyletic eukaryotic clade with no particular affiliation to any specific bacterial group (Fig. 6). Altogether, these results indicate that CYP51A derived from a bacterial gene, indicating that both the MVP (Hoshino and Gaucher, 2018) and genes for the subsequent conversion of squalene to FF-MAS (Fig. 1) may have been inherited via HGT from one or more bacterial donors. This would mean that steroidogenesis was imposed upon the emerging eukaryotic cell by bacteria.

**Fig. 6.**
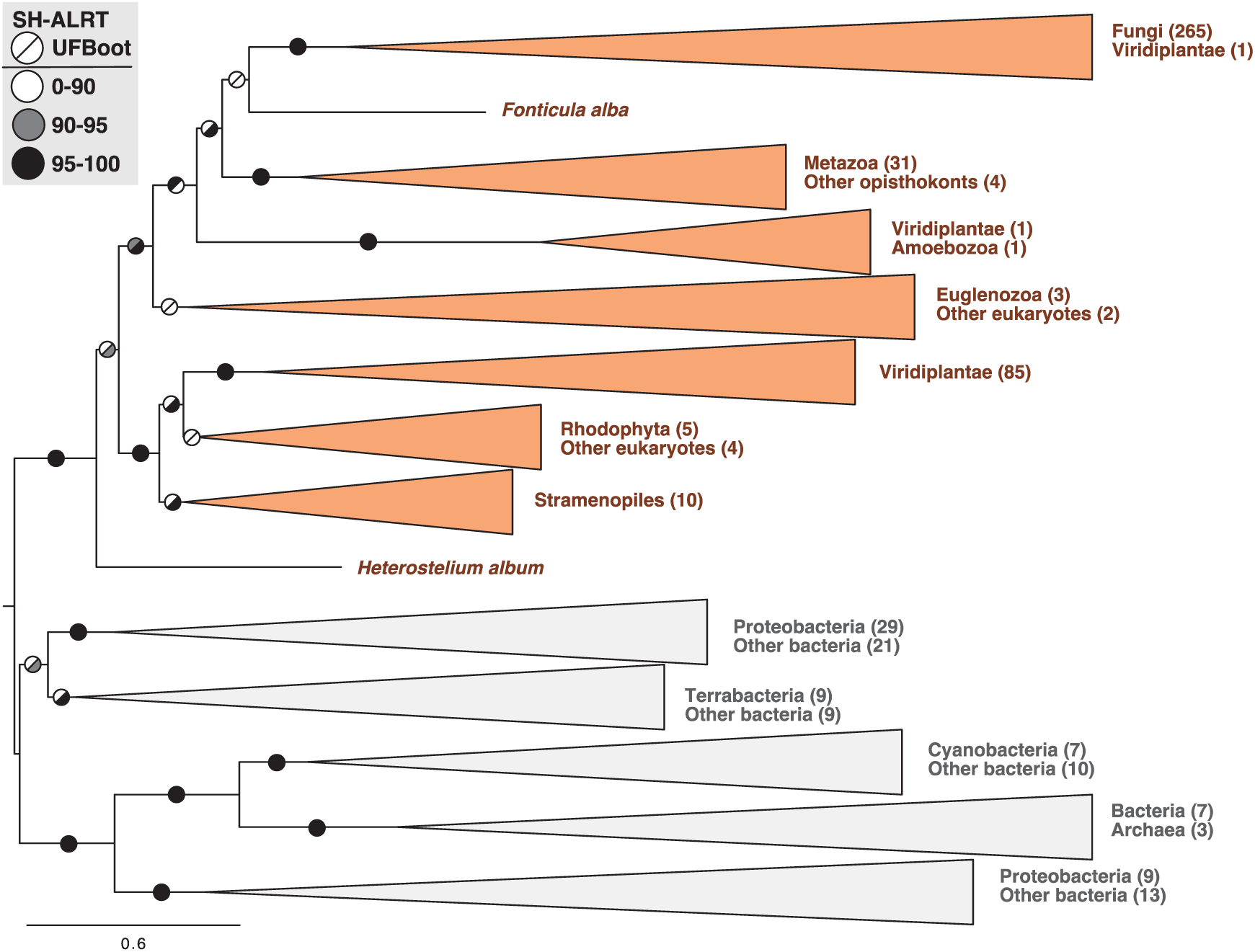
CYP51A is of bacterial origin. Maximum-likelihood phylogenetic reconstruction using IQ-Tree2 under the LG+G4+F model. Branch symbols represent SH-like Approximate Ratio Test (upper left) and Ultrafast bootstrap (lower right), both using 1000 pseudorreplicates. Input alignment contained 533 sequences and 488 sites (Supplementary Appendix F).

## 4. Discussion

Here, we provide the first description of distinct clades of MAPR-related cytb_5M_ clade-1 and eukaryotic cytb_5_-like clade-2 prokaryotic proteins. We show that the distinctive cytb_5_-related MAPR proteins and conventional eukaryotic cytb_5_ proteins arose from separate ancestral eukaryotic cytb_5_-domain genes. MAPR proteins may have arisen from a newly recognized distinct class of CPR cytb_5MY_ proteins, already possessing the residues necessary for the unique tyrosinate-based heme chelation of MAPR proteins. Conversely, cytb_5MY_ protein may alternatively have arisen by HGT of a eukaryotic MAPR protein into CPR bacteria. We cannot confidently discriminate between these opposing models. The cytb_5_ domain is rather small, permitting relatively few informative sites for phylogenetic analyses to reconstruct their evolutionary relationships. The relationship between MAPR and cytb_5MY_ proteins remains therefore uncertain, the latter perhaps being either a sister group to all MAPR proteins or alternatively having arisen from the NEUFC clade within MAPR. While genome annotators currently refer to e.g. “Cytochrome b5-like Heme/Steroid-binding domain-containing protein” to proteins of both clades, we suggest that “Heme/Steroid-binding” should only characterize MAPR-like clade-1.

The newly identified class of MAPR-like cytb_5MY_ proteins are unique to CPR bacteria, which typically have small genomes characteristic of symbionts (Brown et al., 2015; Castelle and Banfield, 2018; Castelle et al., 2018). Some eukaryotic MVP enzymes resemble those from CPR (Castelle and Banfield, 2018; Hoshino and Gaucher, 2018). Therefore, a CPR cytb_5MY_ gene may have contributed to a MAPR-influenced eukaryotic steroidogenic pathway. Genomic organization of CPR cytb_5MY_ genes suggests an ancestral role related to undefined yet regulated redox reactions, which have no known eukaryotic counterparts.

The presence of a cytb_5MY_/pFre/cytb_5_-like operon co-locating with a two-component inducible system (Fig. 5A) in CPR but not other prokaryotes strongly suggests the existence of an inducible CPR-specific process which requires cytb_5MY_ proteins. Regulated bacterial ferric-reductase-like operons are unusual because in prokaryotes these enzymes are largely constitutively expressed, catalyzing the reduction of free flavins which in turn could transfer electrons to a variety of substrates (e.g. various ferric siderophores are known substrates for the ‘classic’ ferric reductases) (Schroder et al., 2003). While eukaryotic ferric reductases are specific for Fe^3+^, in prokaryotes they are merely flavin reductases: reducing diverse flavins, not Fe^3+^ directly. Hence, we can predict neither flavin-specificity, nor the specificity for the terminal pFre reduction substrate: which could be Fe^3+^, copper, other siderophores (Schroder et al., 2003), or entirely different compounds.

The signature tyrosinate heme chelation of MAPR proteins allows them to bind ferric/Fe^3+^ heme tightly, but ferrous/Fe^2+^ heme weakly (Kaluka et al., 2015; Thompson et al., 2007), potentially discharging their heme upon reduction (“one-shot”, or stoichiometric reactivity), with heme-chaperone and conditional status-monitoring implications (Cahill and Medlock, 2017; Kabe et al., 2016b). This specialized functionality may be involved at CPR pFre/cytb_5MY_ loci, speculatively involving Fe^3+^ reduction via cytb_5_-domain proteins, with cytb_5MY_–released Fe^2+^-heme perhaps acting as siderophore (Schroder et al., 2003). Involvement of MAPR proteins in similar biology remains unreported, however the heme-transport function of PGRMC2 (Galmozzi et al., 2019) is conceivably related.

The ancestral MAPR protein had evolved a eukaryotic-specific MIHIR motif, and together with CYP51A was catalysing the oxidative 14-demethylation of lanosterol (Fig. 1) early during eukaryotic evolution, at least before the diversification of fungi and metazoans (Hand et al., 2003; Hughes et al., 2007b). Although cytb_5MY_ proteins may have evolved through HGT from a eukaryotic NEUFC gene to CPR bacteria, a bacterially-inherited MAPR-dependent steroidogenic pathway (Fig. 1) could carry profound implications for early eukaryotic evolution.

Eukaryotic membranes differ from those of archaea, such that a bacterial contribution to their evolution is believed to have been substantial. Cytoplasmically synthesized sterols and membrane lipids required the evolution of new transport pathways to reach the mitochondrion. Through regulation of SREBP1 and SREBP2 activity, PGRMC1 regulates the synthesis of fatty acids and sterols respectively (Cai et al., 2015; Lee et al., 2018; Shimano and Sato, 2017; Suchanek et al., 2005), the core constituents of eukaryotic cell membranes. It is possible that MAPR-mediated membrane trafficking (Cahill et al., 2016; Riad et al., 2020) contributed to the properties of eukaryotic cell membrane whose lipids resemble bacterial more than archaeal membranes, or alternatively, to the trafficking of cytoplasmically synthesized novel lipids to modify mitochondrial membrane properties. This line of reasoning could be highly informed if the NEUFC and NENF MAPR proteins were better functionally charactized, yet that is not the case (Hehenberger et al., 2020).

Four main features hitherto distinguished MAPR from other previously described cytb_5_ proteins: tyrosinate heme-chelation, heme orientation, a MIHIR, and membrane-trafficking functionality. Our analysis, the resemblance of the MIHIR to an actin-cytoskeleton interaction motif (Hehenberger et al., 2020), and the association of PGRMC1 with actin cytoskeletal components (Salsano et al., 2020; Teakel et al., 2020), together show that heme orientation was inherited by MAPR from a protein within the prokaryotic clade-1, whereas the MIHIR appears to have been a eukaryotic invention which may be associated with membrane trafficking. MAPR and cytb_5MY_ proteins also share tyrosinate heme chelation, however the primitive state of the two protein classes remains unclear, as discussed above.

We hypothesise that the original MIHIR-containing MAPR protein may have initially been associated with the heme-related and redox-modulated modification of sterol-containing membranes and their trafficking, perhaps to modulate mitochondrial function. A potential non-vesicular route for mitochondrial sterols is via endoplasmic reticulum (ER)-mitochondrial contacts (EMC): communication mediators between ER and mitochondrial compartments. PGRMC1 is also associated with EMC (Cho et al., 2017). In a preprint, Sabbir and colleagues report that PGRMC1 ablation disrupts the ordered association of mitochondria with the ER at EMC/mitochondrial-associated membranes (MAMs) (Sabbir et al., 2020). Because of the deep evolutionary conservation of the MAPR/CYP51A1 steroidogenic reaction, this MAPR function is possibly related to mitochondrial regulation of an early eukaryote. However, this conjectural hypothesis requires further study.

Sterols are involved in endocytosis at multiple levels in yeast and animals, both in facilitating membrane properties that enable receptor activation, and by participation in an actin-independent post-internalization process (Heese-Peck et al., 2002). Additionally to catalyzing the CYP51A reaction, PGRMC1 is involved in sensing progesterone and progestogen sterol levels (Cahill et al., 2016; Cahill and Medlock, 2017; Ruan et al., 2012), associates with sigma-2 receptor/TMEM97 in sterol transport via endocytosis (Cahill and Medlock, 2017; Riad et al., 2020; Riad et al., 2018; Xu et al., 2011), and regulates sterol/lipid homeostasis via interaction with the SREBP/Insig/SCAP complex, where it is also involved in transcriptional regulation of SREBP-1 (Cahill and Medlock, 2017; Lee et al., 2018; Suchanek et al., 2005).

An evolutionary ancient role of MAPR-regulated sterol synthesis and mitochondrial oxygen consumption status seems plausible. Whereas bacterial hopanoid synthesis does not require oxygen (Saenz et al., 2015), molecular O_2_ is required for steroidogenesis: in particular squalene oxidase epoxidation of squalene (Belin et al., 2018; Kirschvink and Kopp, 2008) and the first lanosterol modification to the 4-methyl sterol FF-MAS by the highly conserved PGRMC1-regulated CYP51A reaction (Stromstedt et al., 1996; Takishita et al., 2017) (Fig. 1). We could therefore model early eukaryotic steroidogenesis as a modulator of oxic/anoxic metabolic responses.

Accordingly, the synthesis of yeast cholesterol-like ergosterol from lanosterol requires oxygen. The fungal Sre1 (the yeast ortholog of SREBP) is cleaved to activate steroidogenesis in a process regulated by hypoxia and the sensing of 4-methyl sterols by Scp1 (the yeast ortholog of SCAP). Changes in 4-methyl sterol levels can lead to an Scp1-dependent proteolysis of Sre1, in a mechanism that has been proposed (based upon presence in mammals and two fungi) to possibly be conserved among unicellular eukaryotes (Hughes et al., 2007a; Hughes and Espenshade, 2008). The fungal Sre1 hypoxic response is accompanied by upregulated increased carbohydrate catabolism and globally reduced transcription and translation in a monumental metabolic switch by the coordination of multiple hypoxic pathways whose interconnectedness remains poorly understood (Burr and Espenshade, 2018).

Interestingly, these dramatic metabolic changes are reminisicent of PGRMC1-dependent effects on mammalian cell culture cells (Thejer et al., 2020a; Thejer et al., 2020b), however, any mechanistic link remains uninvestigated. Both the *Schizosaccharomyces pombe* Dap1 MAPR protein (a PGRMC-like protein (Hehenberger et al., 2020)) and the CYP51A homolog are transcriptionally-induced by hypoxia (Hughes et al., 2007b). PGRMC1 is also induced in the hypoxic zone of human tumors (Neubauer et al., 2008), possibly representing conserved ancestral MAPR hypoxic functionality. That could be related to a proposed role of mitochondria in O_2_ detoxification during early eukaryotic evolution (Imachi et al., 2020; Kurland and Andersson, 2000) (either pre- or post-LECA), and to the profound effects of PGRMC1 on mitochondria (Thejer et al., 2020a), and on glycolytic metabolism (Sabbir, 2019; Thejer et al., 2020b).

MAPR genes are present in at least plants, opisthokonts, *Guillardia*, *Dictyostelium* and Stramenopiles (Hehenberger et al., 2020) (Fig. 2) (there has been no systematic phylogenetic survey), which suggests an early MAPR origin in eukaryotes. Many aspects of modern meiosis reflect functions shared by these groups (Loidl, 2016), which may extend to MAPR proteins and steroids. Yeast meiotic membrane fusion requires sterols (Aguilar et al., 2010). Mammalian PGRMC1 is attached to the kinetochore microtubules of meiotic/mitotic spindles (Juhlen et al., 2018; Luciano and Peluso, 2016), and to metaphase centromeres (Juhlen et al., 2018; Luciano et al., 2010; Terzaghi et al., 2016). The products of the reaction catalyzed by PGRMC1 and CYP51A are either FF-MAS (Fig. 1) or dihydro-FF-MAS, both 4-methylsterol inducers of meiosis (Fakheri and Javitt, 2011; Mitsche et al., 2015). Similar 4-methylsterols activate meioisis in plants as well as animals (Darnet and Schaller, 2019) and are induced by hypoxia in yeast (see above), suggesting the operation of an ancient and conserved mechanism. Mammalian PGRMC1 mediates a progesterone-induced block of meiotic progression (Guo et al., 2016; Luciano and Peluso, 2016). Yeast *Saccharomyces cerevisiae* Dap1 (PGRMC1 ortholog) mutation also leads to cell cycle defects (Hand et al., 2003). As tantalizingly as these PGRMC attributes align with possible ancestral eukaryotic biology, their phylogenetic distribution has not been investigated, and so functional extrapolation to an early eukaryotic ancestral MAPR gene remains speculative.

Whereas post-cholesterol steroid hormones evolved in vertebrates, the synthetic pathway from lanosterol to cholesterol involves production of diverse bioactive sterols (Mitsche et al., 2015) which sequentially appeared during evolution, where CYP51A/PGRMC1 catalyze the first step after lanosterol is produced (Fig. 1). The closest related bacterial proteins to eukaryotic CYP51A (Fig. 6) suggest that HGT of both the ancestral MAPR and CYP51A genes to an early eukaryote from genomes other than the proto-mitochondrial alphaproteobacterium to metabolize a lanosterol-like hopanoid and regulate its transport may have occurred early in eukaryotic evolution.

The MAPR-dependent first sterol modification (Fig. 1) may have originally been catalyzed by bacterial enzymes to manipulate an associated early eukaryotic cell, and drive a metabolic switch that is still operative in humans. These considerations refocus the context of highly conserved PGRMC1 biology across the spectrum of heme/lipid metabolism and transport, and affected diseases such as metabolism and glucose regulation, cancer, fertility, and neuropathologies such as Alzheimer’s disease (Cahill et al., 2016; Cahill and Medlock, 2017; Izzo et al., 2020; Lee et al., 2018; Peluso and Pru, 2014; Riad et al., 2020; Riad et al., 2018; Sabbir, 2019; Xu et al., 2011).

## CRediT authorship contribution statement

**Daniel Tamarit**: Data curation, Formal analysis, Funding acquisition, Investigation, Methodology, Writing - review & editing.

**Sarah Teakel**: Formal analysis, Writing - review & editing.

**Michealla Marama**: Formal analysis, Writing - review & editing.

**David Aragão**: Methodology, Resources, Writing - review & editing.

**Svetlana Y. Gerdes**: Data curation, Formal analysis, Investigation, Methodology, Writing - original draft.

**Jade K. Forwood**: Funding acquisition, Methodology, Resources, Supervision, Writing - review & editing.

**Thijs J. G. Ettema**: Funding acquisition, Methodology, Resources, Supervision, Writing - review & editing.

**Michael A. Cahill**: Conceptualization, Formal analysis, Funding acquisition, Investigation, Methodology, Project administration, Resources, Writing - original draft.

## Acknowledgements

This research was supported by Charles Sturt University (CSU) School of Biomedical Sciences (SBMS) internal awards to M.A.C. and PhD scholarship support to S.T; by the Swedish Research Council (VR grants 2015-04959 to T.J.G.E., and 2018-06609 to D.T.), by the European Research Council (ERC consolidator grant 817834 to T.J.G.E.). Computation was performed on resources provided by the Swedish National Infrastructure for Computing through the Uppsala Multidisciplinary Center for Advanced Computational Science (UPPMAX) under projects SNIC-2020-5-37 (to T.J.G.E. and Felix Homa) and SNIC-2020-15-113 (to D.T.). This research was partly undertaken on the macromolecular crystallography beamlines at the Australian Synchrotron, part of ANSTO. Elisabeth Hehenberger (Department of Botany, University of British Columbia, Vacouver, Canada) provided protist MAPR sequences. Aya Narunsky (The Nir Ben-Tal Group at the Department of Biochemistry and Molecular Biology of Tel-Aviv University) assisted with Consurf results. Amy Medlock (University of Georgia), George John (CSU), and Mike Dyall-Smith (University of Melbourne) commented upon draft manuscripts.

Supplementary Information is linked to the online version of the paper.

## Research data for this article

The structure of Hadesarchaea YNP_N21 cytb_5M_ (KUO41884.1) has been deposited in the Protein Data Bank with accession number 6NZX. The PDB validation report is included as Supplementary Appendix G. Other data and materials are supplied as supplemental data or are available upon request.

---

**Fig. A1.**
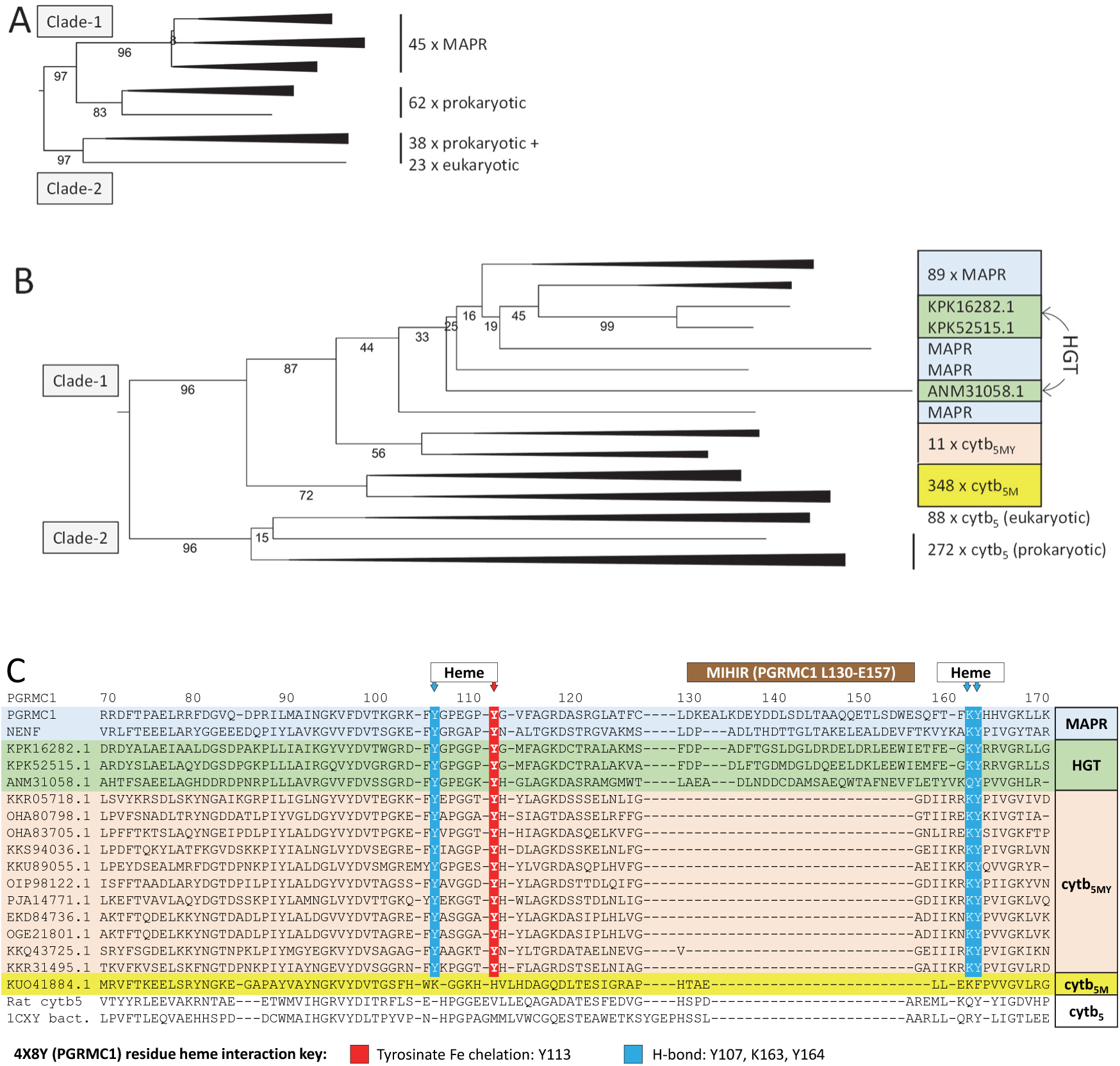
BLASTp hits from MAPR or cytochrome b5 protein search strings form two discrete clades and MAPR proteins are related to prokaryotic cytb_5MY_. (A) Neighbour-Joining 1,000 bootstrap maximum likelihood phylogenetic tree under the Poisson substitution model, constructed from the alignment of 176 protein sequences by MAFFT. (B) Neighbour-Joining phylogenetic tree under the Poisson substitution model, constructed as per (A)from the alignment of 814 protein sequences.Sequences from preliminary analyses are combined with independent eukaryotic MAPR and cytb_5_ sequences (see methods) in a 65 gap-free site alignment. 11 proteins clustering closer to MAPR than other bacterial proteins are indicated (cytb_5MY_). See Supplementary Appendix A for FASTA and tree files of all preliminary investigations, as well as the combined tree depicted in this panel. (C) MAFFT alignment of MAPR with bacterial HGT and cytb_5MY_ proteins. Heme-interacting residues and the MIHIR in PGRMC1 are indicated. The heme interaction key follows the PGRMC1 4X8Y structure (Kabe et al., 2016a). Alignment with representative clade-2 cytb_5M_ sequence (KUO41884.1, Hadesarchaea cytb_5M_), and eukaryotic (NP_085075.1, rat cytb5B) or bacterial (1CXY_A *Ectothiorhodospira vacuolata*) clade-1 proteins is provided below.

**Fig. A2.**
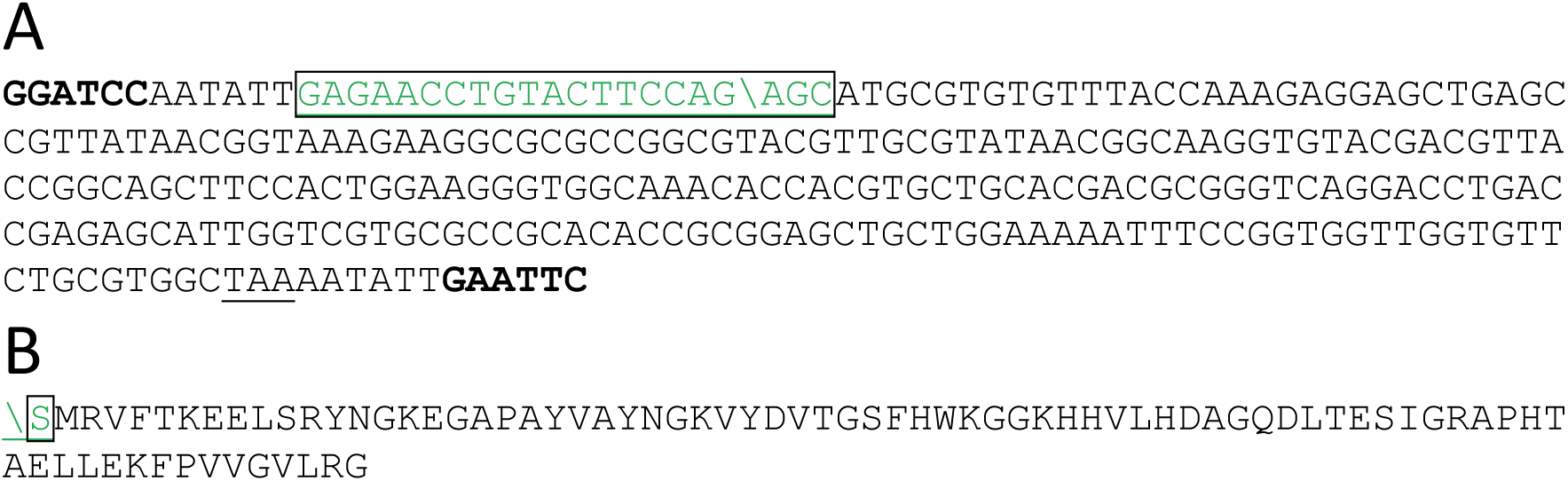
Sequences of pGEX4T1_KUO41884 and the crystallised protein. (A) KUO41884.1 sequence was cloned into the BamH1 and EcoRI sites (bold) of pGEX4T1 with added Tobacco Etch Virus (TEV) cleavage site (boxed, cleavage site in translated protein marked by \). The stop codon is underlined. (B) Sequence of the crystallized protein, corresponding to KUO41884.1 with additional N-terminal S residue after TEV cleavage.

**Fig. A3.**
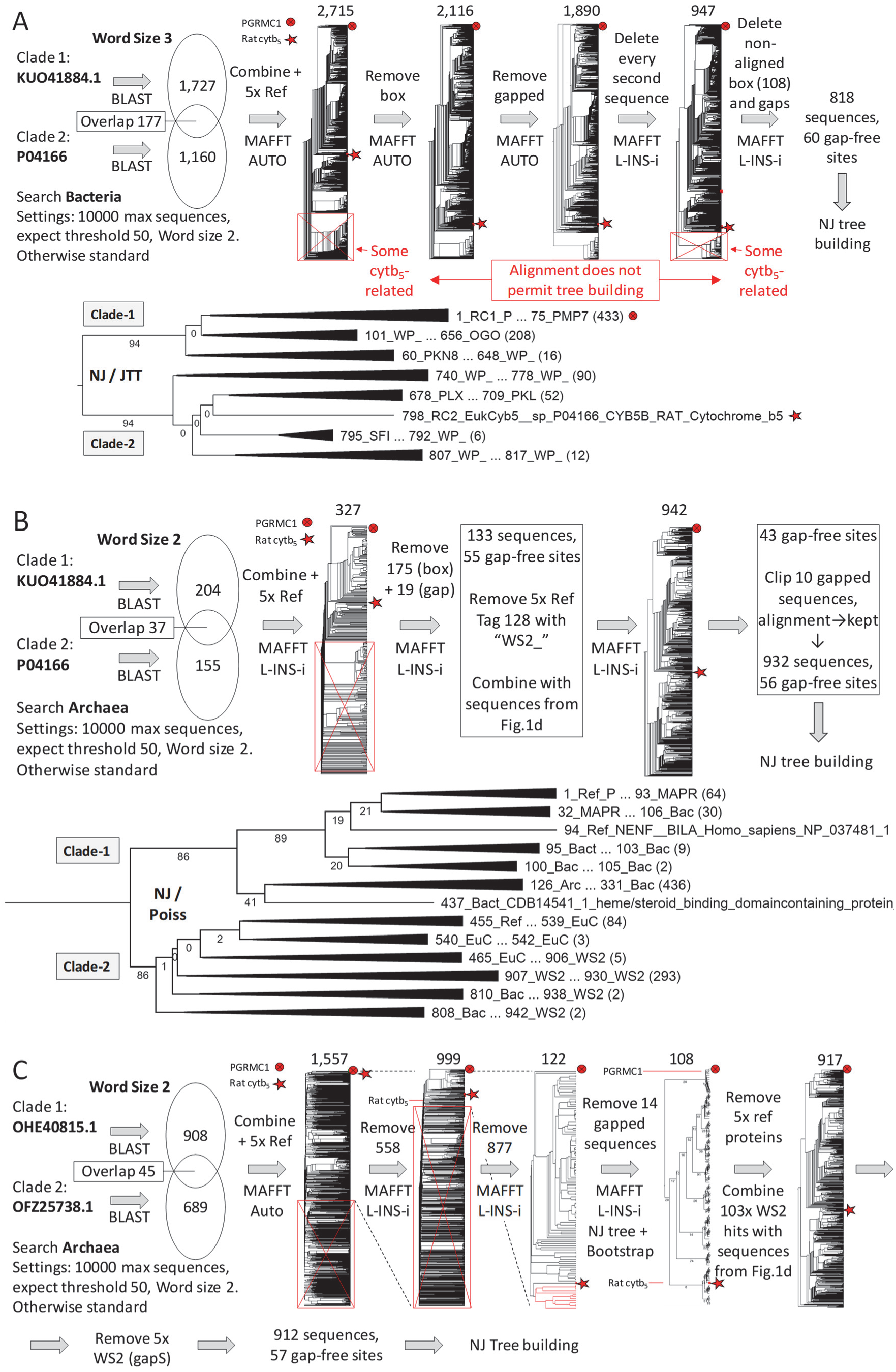
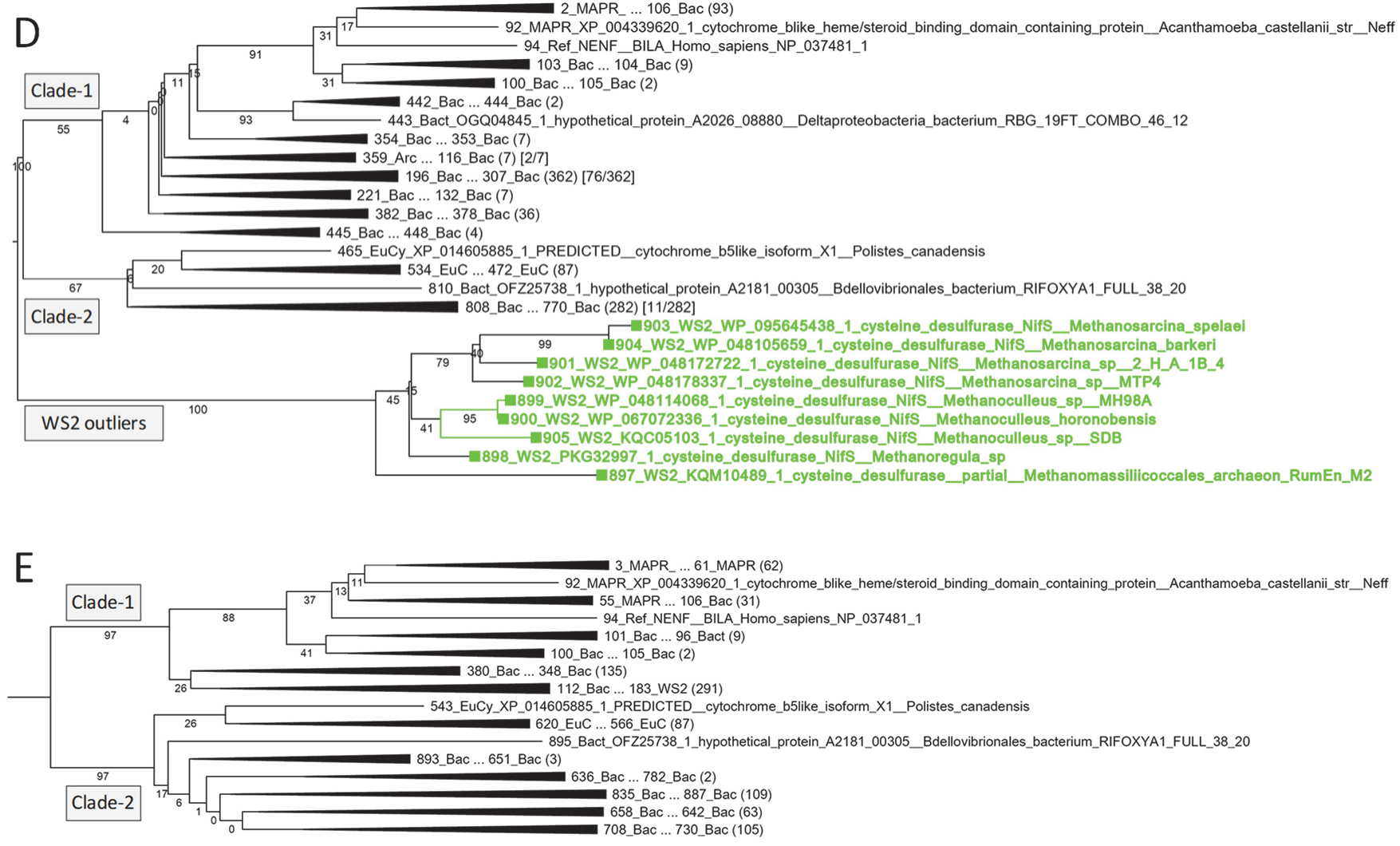
Reduced stringency BLAST does not detect a sequence continuum between clades 1 and 2. (A) Schematic of reduced stringency BLASTp analysis using Word Size 3 (see Methods) against the bacterial data base. The indicated sequences from clade-1 and clade-2 were used as BLASTp search strings, and combined with three clade-1 (PGRC1 NP_006658.1, NENF NP_037481.1, KXH77621.1) and two clade-2 (CYB5B_RAT P04166, and WP_103012903.1) reference sequences, tagged “>RC2_” and >RC2_” respectively. Iterative processing of the data as shown led to 818 gap-free sequences which were used for NJ treebuilding. Bootstrap support for separate clades 1 and 2 was 94% for JTT (depicted), 89% (WAG), and 93% (Poisson). Of the 159 sequences in clade-2 of this tree (excluding two clade-2 reference sequences), only 4 (WP_093087318.1, KKQ27723.1, KXK26085.1, GBD33942.1) were detected by both the clade-1 (KUO41884.1) and clade-2 (P04166) BLAST query sequences. (B) Using the same BLASTp search strings as panel a, the smaller archaeal data base was searched with Word Size 2 (WS2), followed by iterative MAFFT data refinement as schematically depicted. 103 WS2 hits were combined with the sequences from Fig. A1A and iteratively refined to provide a robust tree architecture of 932 sequences used for NJ tree building. Bootstrap support for the existence of separate clades 1 and 2 was 85% (JTT), 76% (WAG), and 86% (Poisson, depicted). Of the 34 WS2 sequences present in clade-2, 10 (AQS28389.1, RLI96648.1, RME54213.1, PIN98782.1, RME31891.1, RME52388.1, PIN95630.1, PIZ83262.1, OIO41921.1, RME53069.1) were also BLASTp hits for clade-1 search string KUO41884.1. (C) Word size 2 BLASTp with independent query sequences to panel b. Because of the relatively low hit number of panel b the analysis was repeated using the indicated two outlying clade-1 and clade-2 sequences from Fig. A1A. Iterative data refinement 103 WS2 hits were combined with sequences from Fig. A1A to yield ultimately 912 sequences with 57 gap-free sites for tree building. (D) NJ/Poisson tree (1,000 bootstrap) of the process from panel c, including nine WS2 sequences with 100% bootstrap support as outliers to clades 1 and 2 (green). Bootstrap support for clades 1 and 2 is below 70% for JTT, WAG and Poisson NJ trees (depicted), however a group of cytb_5_-related proteins form an outlying distinct clade with 100% bootstrap support. Of the 11 WS2 sequences present in clade-2, none were also BLASTp hits for clade-1 search string OHE40815.1. Of the 9 outlying WS2 sequences, one (KQM10489) was also a BLASTp hit for clade-1 search string OHE40815.1. (E) Identical tree to panel d, with nine WS2 outlying sequences removed prior to MAFFT alignment. This tree contains 78 WS2 sequences that fall within clades 1 or two, but none that cluster between those clades. Bootstrap support for clades 1 and two is 93% (JTT), 89% (WAG) and 97% (Poisson, depicted). 11 proteins were detected by both clade-1 and clade-2 search querie sequences. No proteins intermediate in sequence between clades 1 and 2 were found. For identities of all cytb-L proteins detected in CPR bacteria see Supplementary Appendix C.

**Fig. A4.**
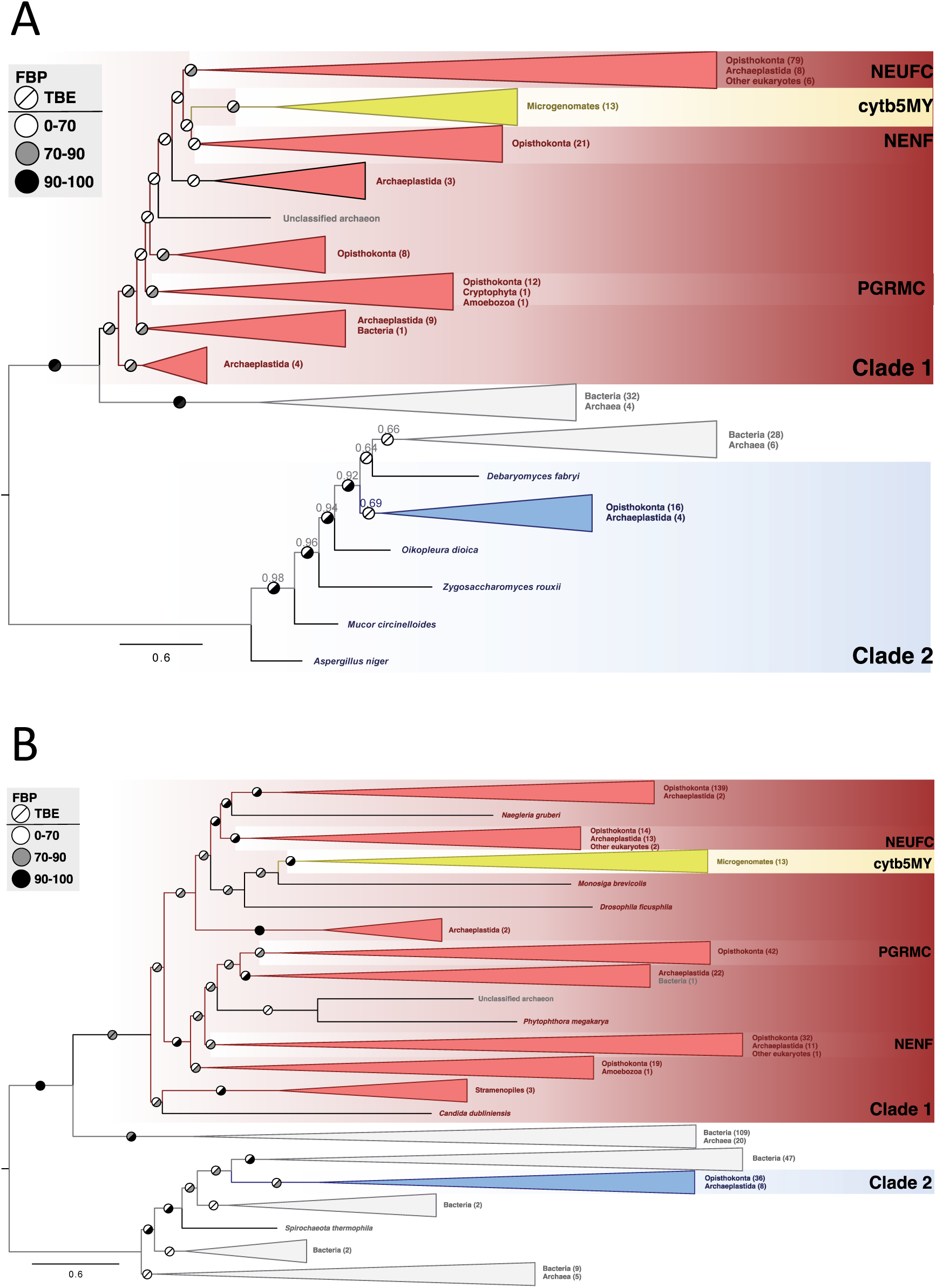
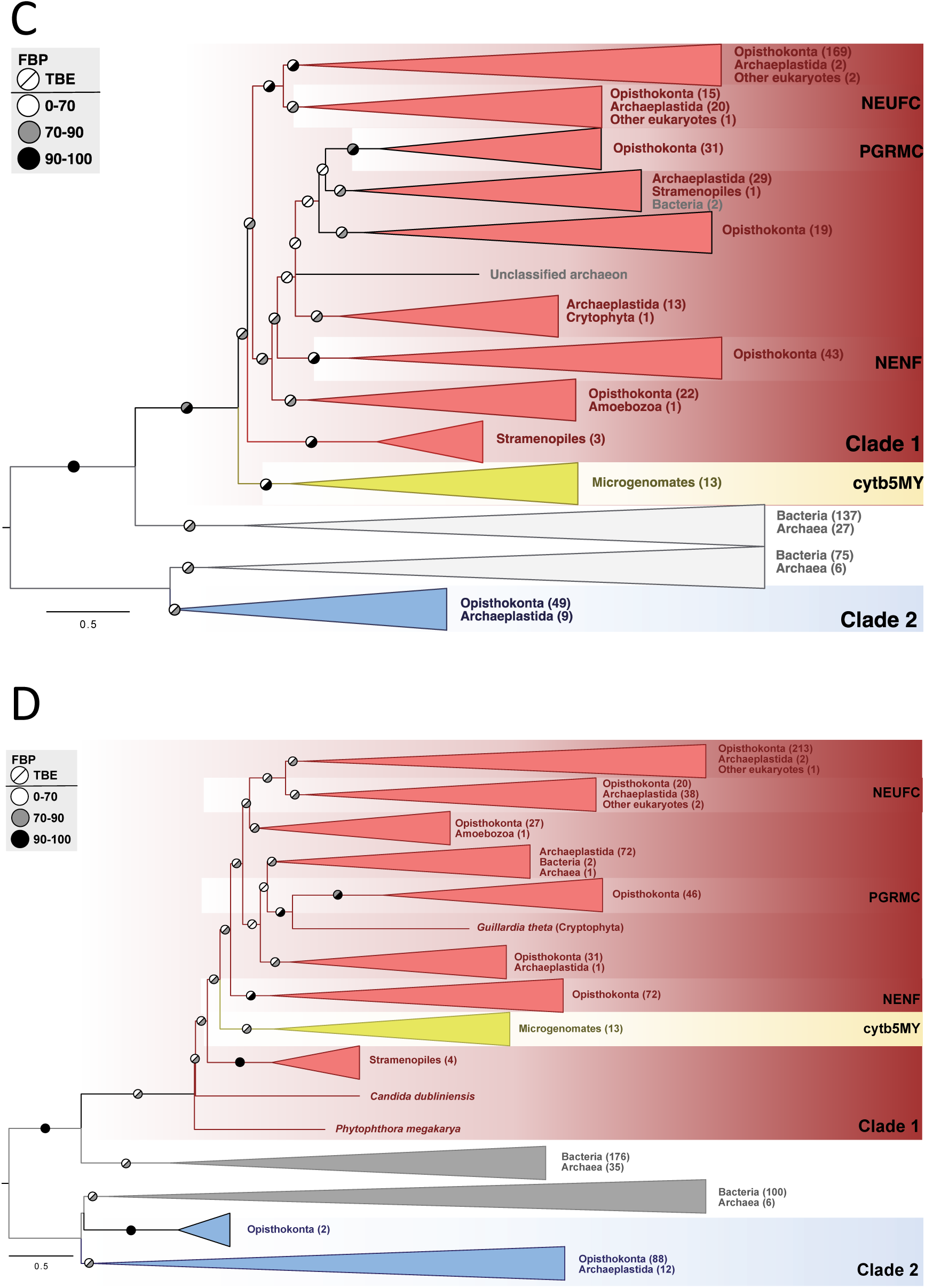
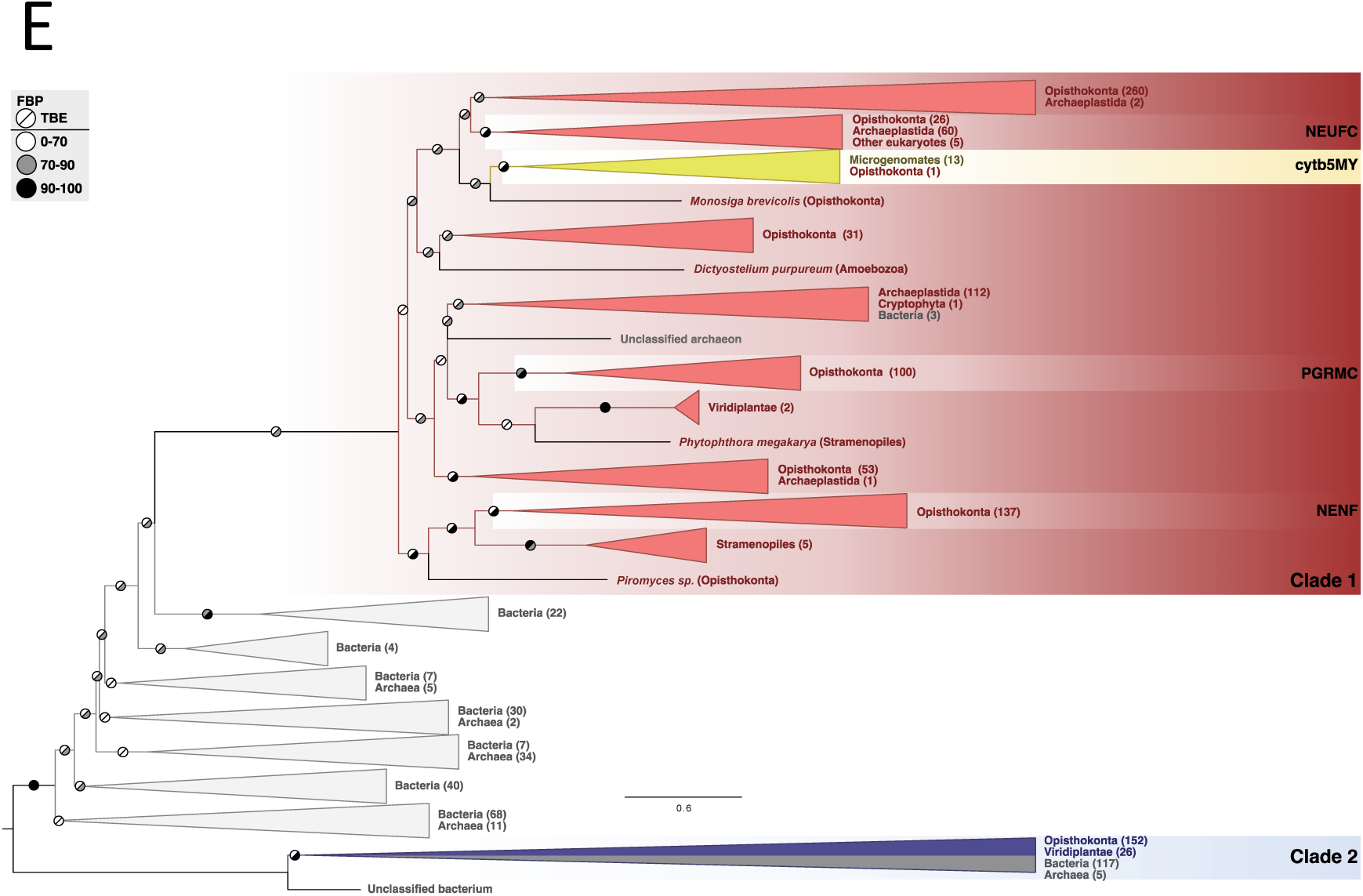
Phylogenetic reconstructions of cytb_5MY_ tree position with alternative sequence datasets. Maximum-likelihood phylogenetic reconstructions using IQ-Tree2 under the PMSF approximation of the LG+C60+R8+F model (A,B), LG+C50+R8+F (C), LG+C40+R8+F (D) and LG+C50+R8+F (E). Input alignments were based on the reduction of the 2470 gathered sequences at different levels of similarity: 50% (A), 65% (B), 70% (C), 80% (D) and 90% (E), resulting in 274, 570, 702, 968 and 1384 sequences, respectively. Branch symbols indicate Felsenstein Bootstrap Proportions (FBP; upper left half) and Transfer Bootstrap Expectation (TBE; lower right half) interpretations of 100 non-parametric bootstrap pseudorreplicates. Sequence details are available as Supplementary Appendix D.

**Fig. A5.**
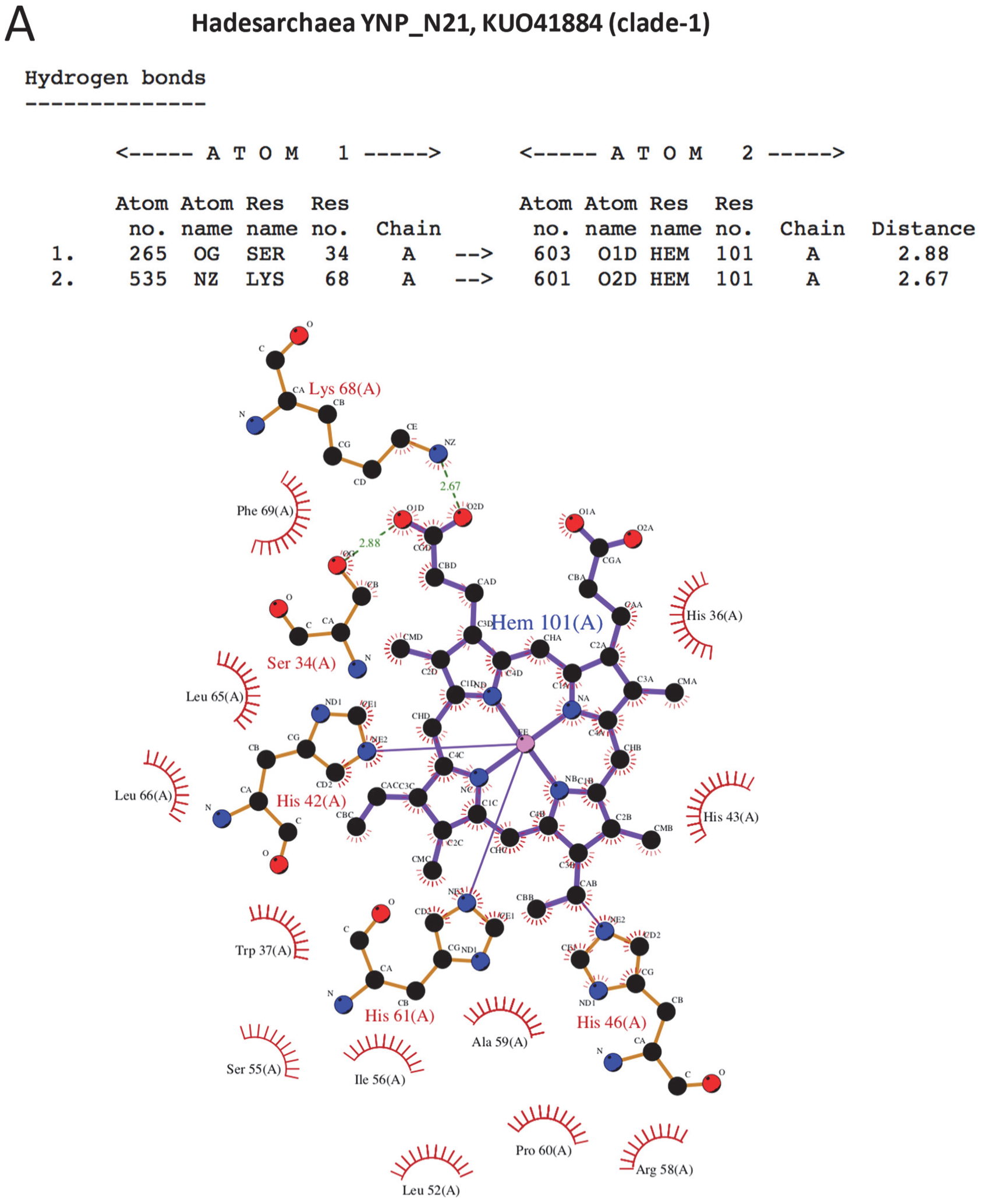
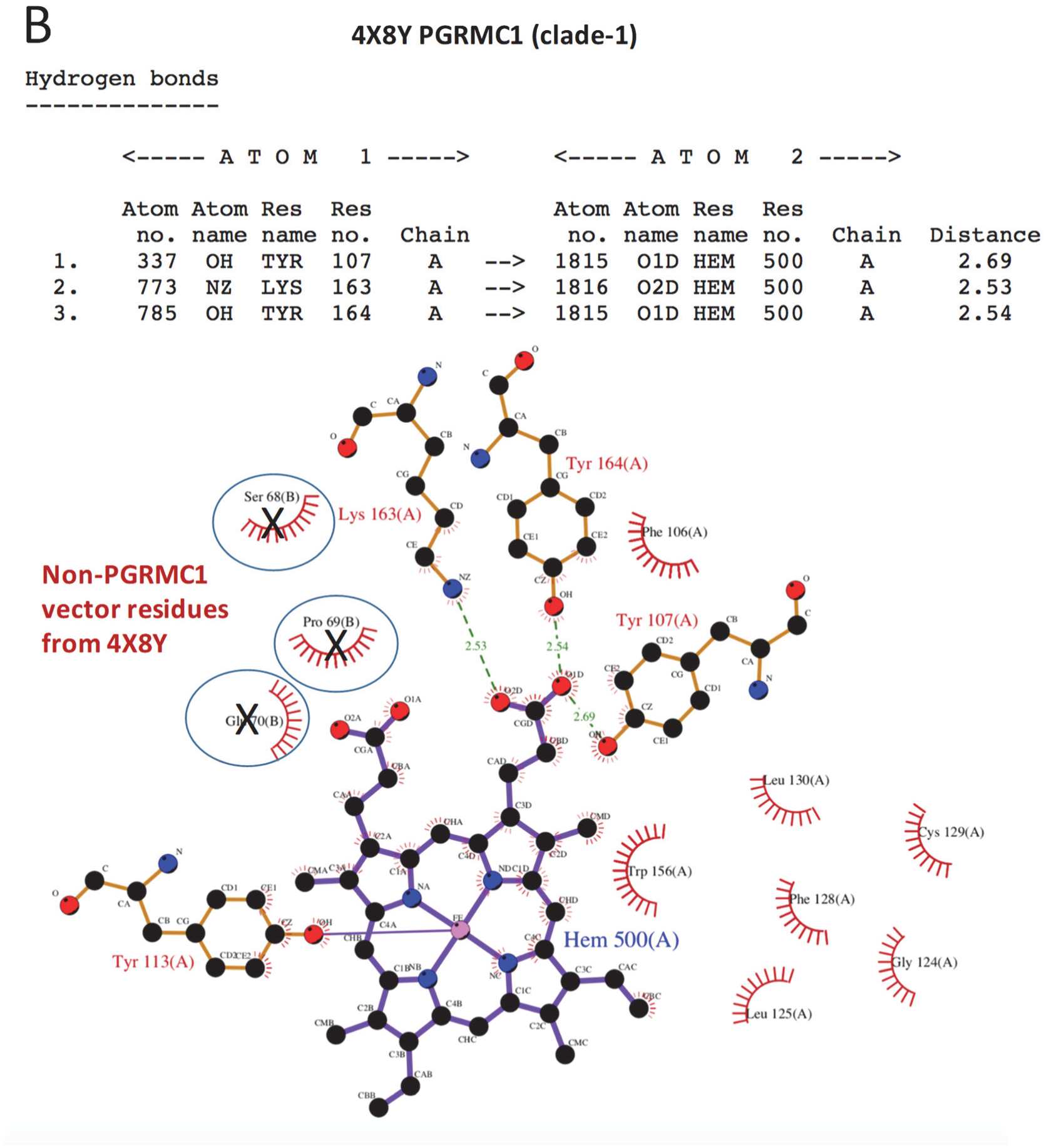
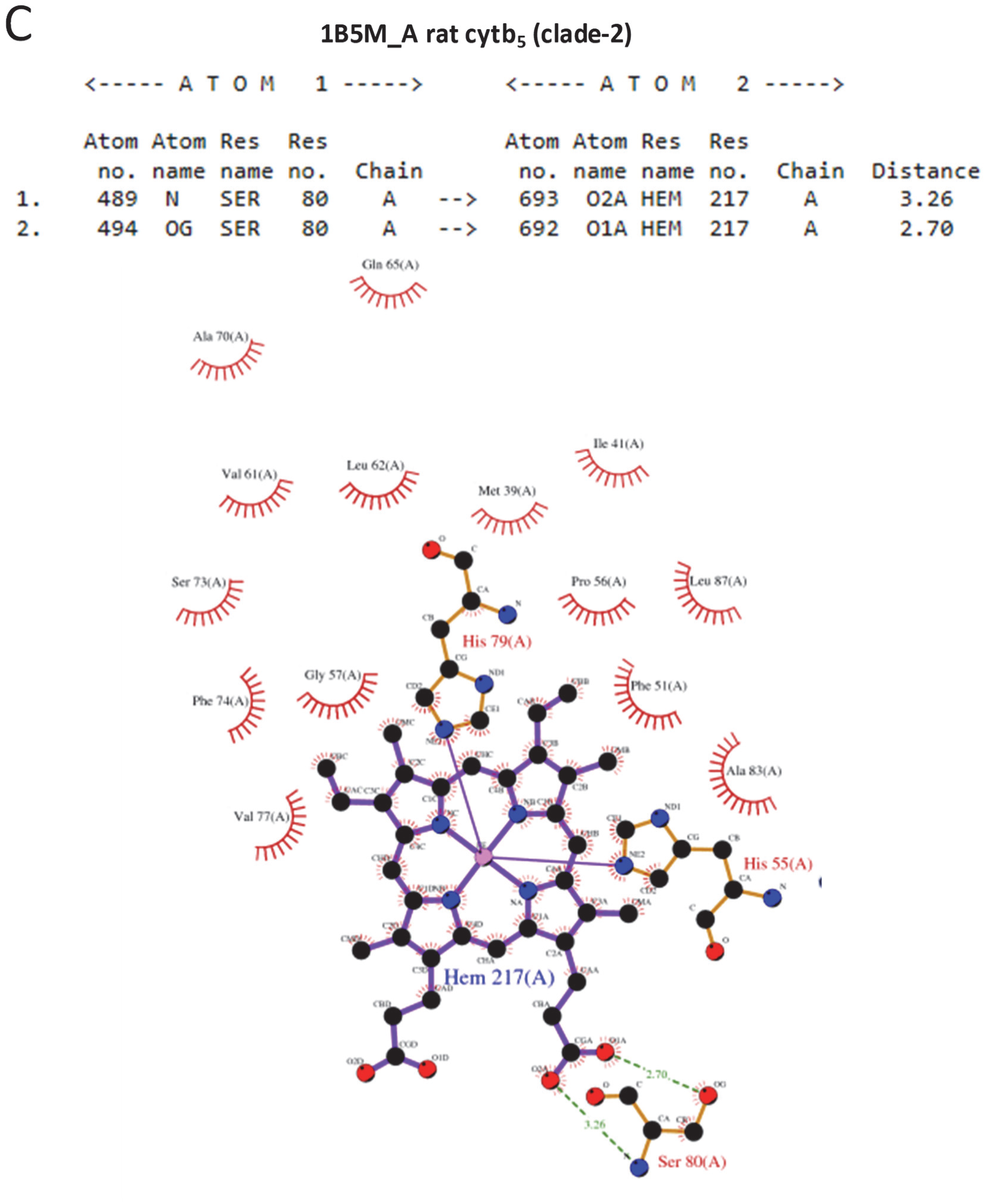
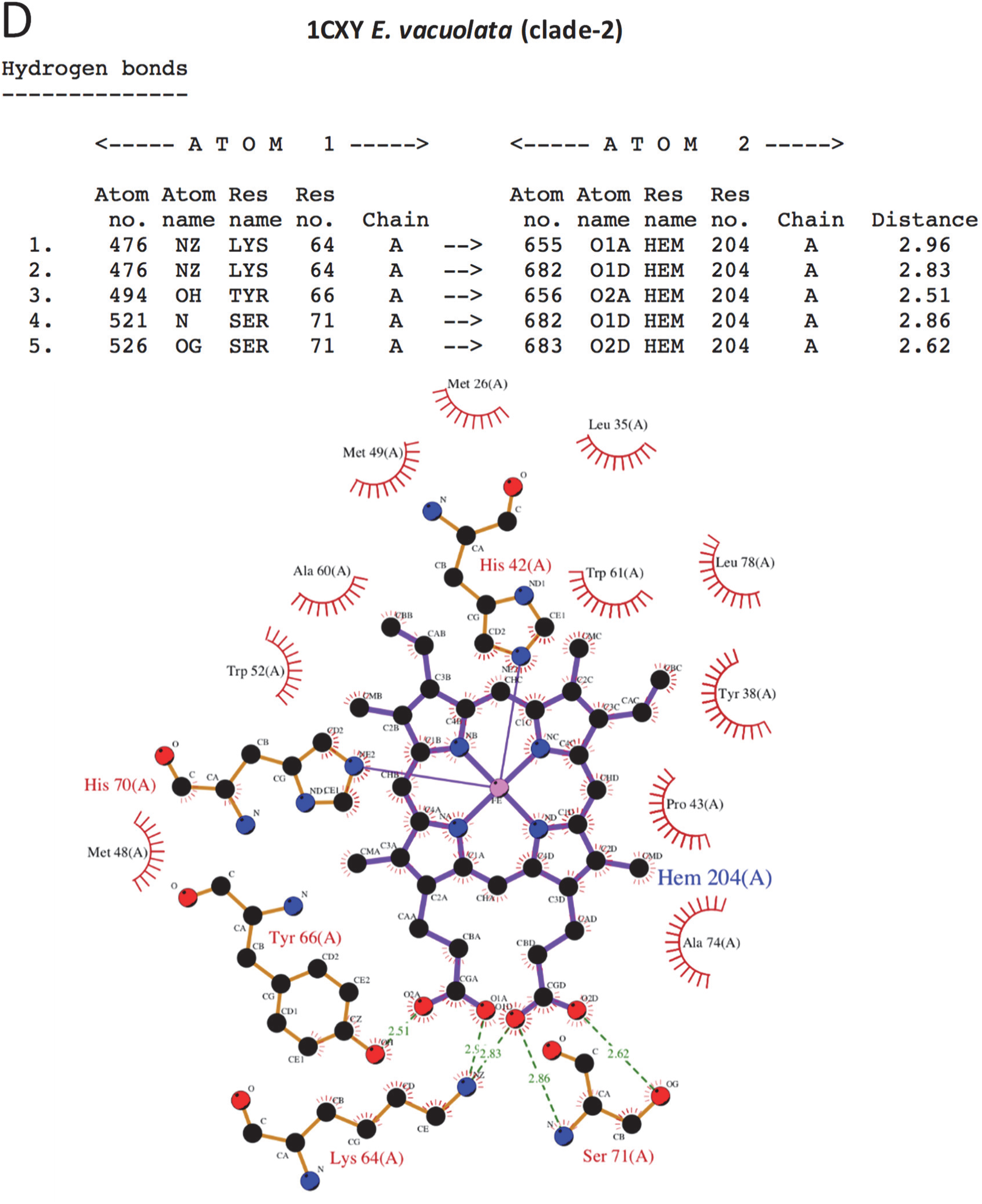
Heme-interacting residues from Fig. 4E. H-Bonds with heme propionate groups are given at the top of respective panels, with LIGPLOTs below. (A) Hadesarchaea YNP_N21, cytb_5M_ KUO41884.1 (clade-1). (B) Human PGRMC1 4X8Y (Clade-1). Non-PGRMC1 N-terminal residues in the 4X8Y structure from the bacterial expression vector that make contact with heme are indicated by a circled X. The N-terminus is not proximal to heme in the dimeric structure (Fig. 3C). These residues make contact from adjacent symmetry units. The PGRMC1 MAPR domain of 4X8Y was also N-terminally truncated relative to Fig. A1B by 2 residues. N-terminal non-PGRMC1 residues (from the expression vector fusion protein) of adjacent crystal symmetry units contacting heme in the PGRMC1 4X8Y structure could artefactually stabilize the heme-dependent dimer. A native PGRMC1 MAPR-domain structure would be desirable to confirm biological heme-dependent PGRMC1 dimerization. Hydrophobic interactions between heme and S68, P69 and E70 from 4X8Y PGRMC1 structure are not shown for Fig. 4E because these represent non-PGRMC1 residues from the bacterial expression vector. (C) Rat mitochondrial cytb_5B_ 1B5M_A (Clade-2). Numbering is according to the corresponding rat cytb_5_ sequence NP_085075.1. (D) *E. vacuolata* 1CXY (clade-2).

**Fig. A6.**
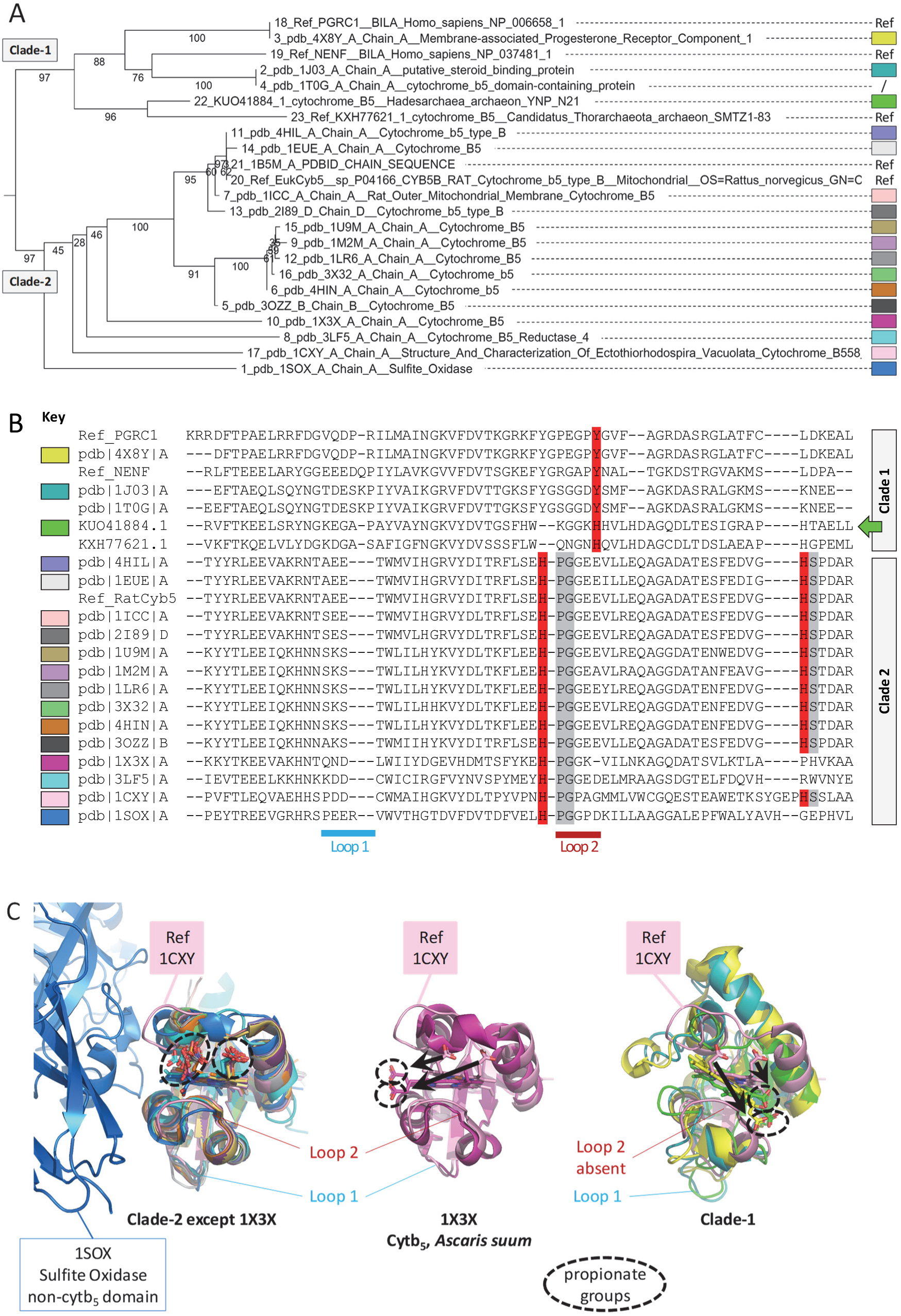
The most similar non-MAPR cytb_5_-domain published structures to Hadesarchaea cytb_5M_ are from clade-2. (A) The thirty PDB structures most similar by DALI to KUO41884.1 (Hadesarchaea cytb_5M_, this study) were from seventeen proteins (colored boxes). MAFFT alignment followed by NJ Poisson putative tree construction (1,000 bootstrap) showed that all except the recognised MAPR proteins (4X8Y, 1J03, 1T0G) belonged to clade-2. Reference sequences include clade-1 proteins NP_006658.1 PGRMC1, NP_037481.1 NENF, and archaean cytb_5M_ KXH77621.1 (no structure available), and clade-2 NP_085075.1 rat cytb_5_. Note the KRR N-terminal PGRMC1 MAPR residues absent from 4X8Y. Other conventions follow Fig. A1. (B) Clade-2 proteins with solved structures most similar to KUO41884.1 all shared the HPG heme iron chelation motif involving loop 2 (Fig. 4E). The HS motif was also present in all but chicken sulfite oxidase (1SOX) (Kisker et al., 1997), the cytb_5_-reductase domain of NADH cytochrome b_5_ oxidoreductase (3LF5) (Deng et al., 2010), and *Ascaris suum* cytb_5_ (1X3X). The latter is known to be an atypical cytb_5_ protein (Yokota et al., 2006), and also exhibited divergent tree clustering in “A”. Red: iron atom chelation. Grey: adjacent conserved heme hydrophobic interaction residue (following Fig. 4E). The color key corresponds to proteins shown in a. (C) All clade-2 proteins except 1X3X exhibit similar heme orientation to reference 1CXY (left panel). In 1X3X the propionates are rotated approximately 90° to the left (centre panel, arrows), whereas in clade-1 proteins the propionates are rotated to the right relative to clade-2 (arrows, right panel), and bind at a tilted plane relatively to clade-2, prominently permitted by the absence of loop 2. Superimposable Arabidopsis MAPR structures 1J03 (Yoshitani et al., 2005) and 1T0G (Song et al., 2004) do not include heme. Only 1J03 is included.

**Fig. A7.**
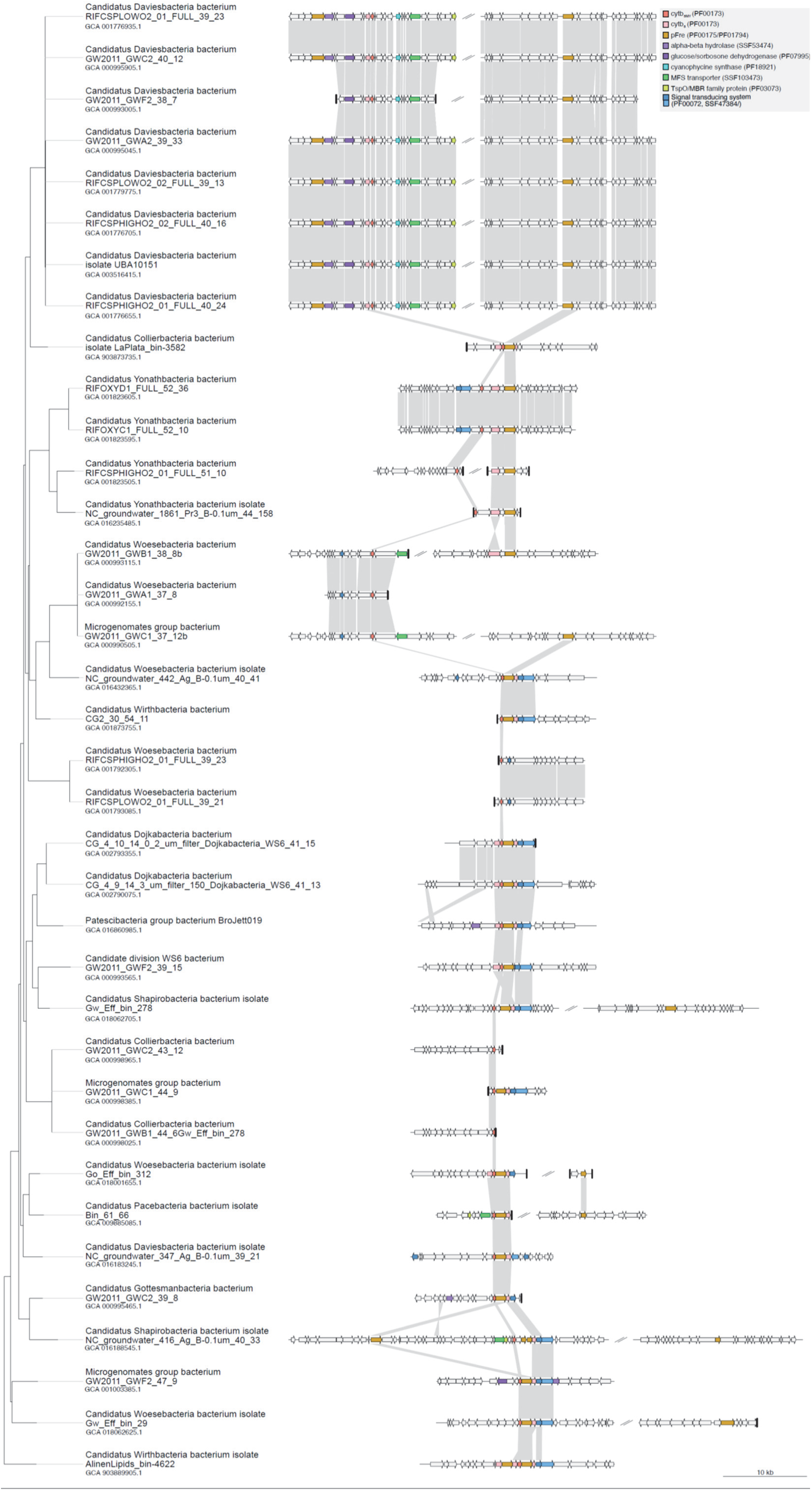
Synteny of cytb_5MY_ and pFre gene clusters. Gene neighbourhoods are shown at a 10-kb distance from cytb_5MY_ and pFre genes. Functional annotation of selected genes is shown according to the legend colors. Arrows represent genes and thick vertical lines, contig boundaries. Connecting lines indicate similarity found as best reciprocal blastp hits with e-values lower than 1e-5. The tree shown at the left side is the phylogeny shown in Fig. A8.

**Fig. A8.**
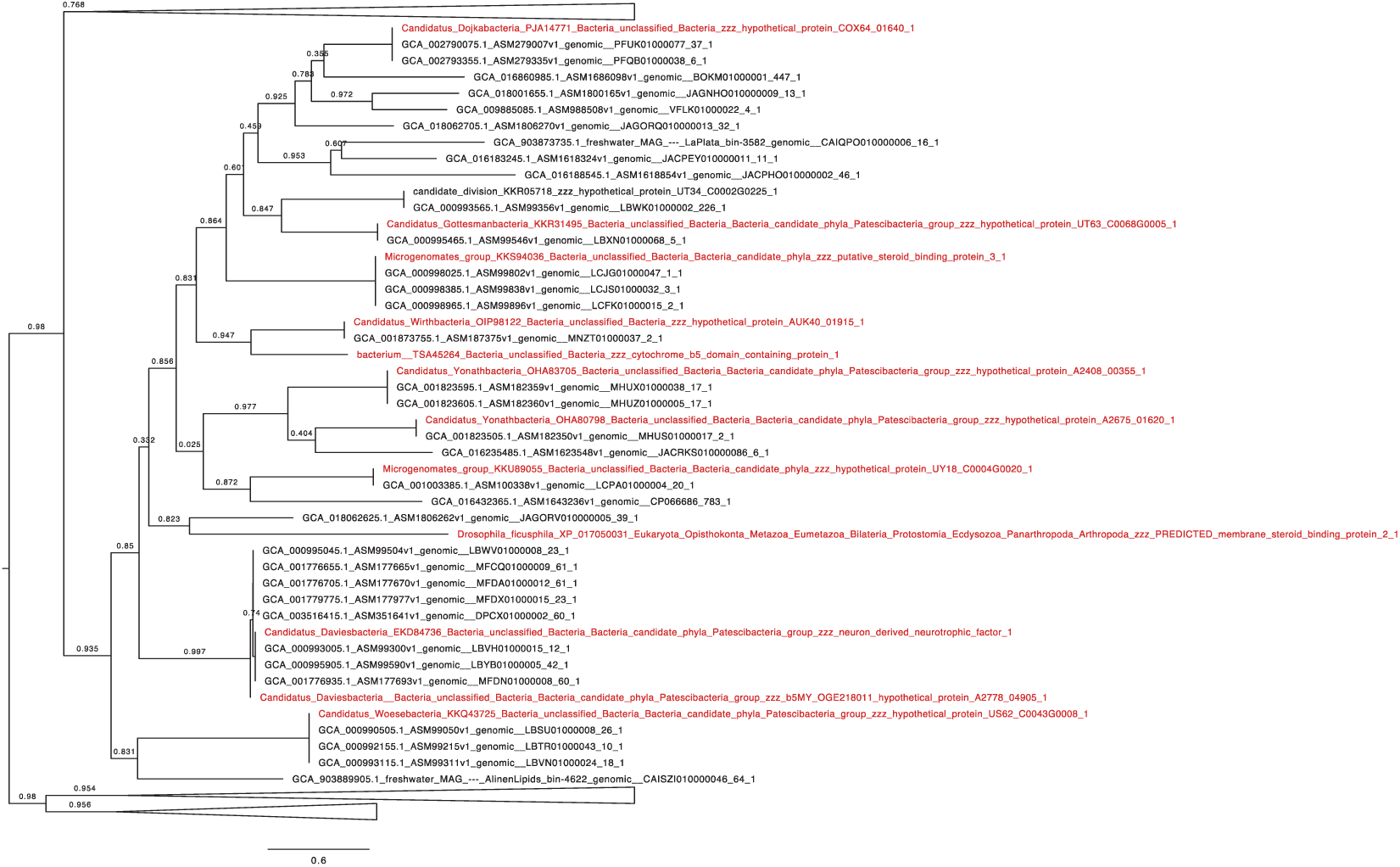
CPR cytb_5MY_ gene similarities. Phylogenetic tree obtained by Fasttree2 including all sequences used for the phylogenetic tree in Fig A3B. Sequences included in the previous analysis are marked in red. Only the cytb_5MY_ clade is shown expanded.

**Fig. A9.**
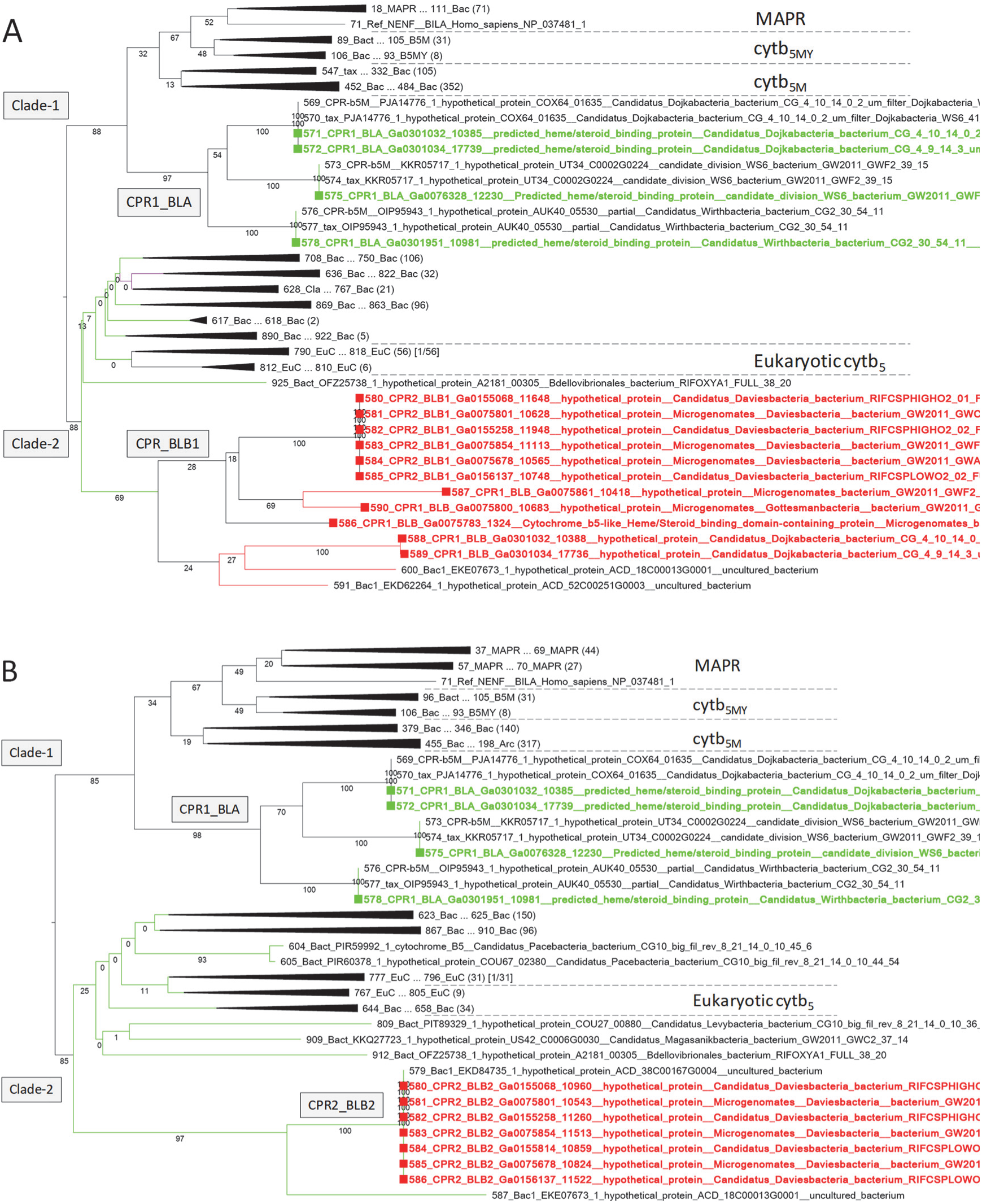
Affiliation of pFre-associated Cytb_5_-L proteins. (A) Neighbour-joining tree under the Poisson model and showing 1,000 bootstrap pseudorreplicates, inferred from 916 sequences, including cytb_5_-like type A (CPR_BLA) and or cytb_5_-like type B1 (CPR1/2_BLB1) proteins (Fig. 5A) as well as the proteins from Fig. A1A. It was impossible to align clades-1 and 2 together with both cytb_5_-L type B1 and B2 proteins and obtain sufficient gap-free aligned sites for tree building. This panel shows the tree with CPR1/2_BLB2 sequences omitted. These sequences include those from Fig. A1A, supplemented with CPR cytb_5_-domain proteins detected in IMG analyses (labelled: “CPRx”, and the results of BLASTp of a cytb_5MY_ search string (KUO41884.1) against the NCBI CPR database (taxid:1783234) (labelled “tax_”). Identical sequences from the three input sources were not removed. (B) All details are identical to “A”, except that CPR1/2_BLB1 sequences were omitted and CPR2_BLB2 sequences included in the alignment to give 912 MAFFT-aligned sequences. The position of the CPR_BLA clade relative to cytb_5M_ is uncertain with only 32% and 34% bootstrap support in both analyses. The corresponding WAG tree for panel “A” positioned the CPR1_BLA clade deep within the cytb_5M_ branch (not shown). Complete tree files and PDF printouts of the expanded trees for this figure are available as Supplementary Appendix E.

**Table A1.** Details of cytb_5MY_ proteins from Fig. A1B. All species are CPR bacteria (Castelle and Banfield, 2018). The details of all cytb_5MY_ and cytb_5M_ proteins detected in CPR bacteria are provided in Supplementary Appendix C.

| Accession | Protein name [species] | Reference |
| --- | --- | --- |
| KKR05718.1 | hypothetical protein UT34_C0002G0225 [candidate division WS6 bacterium GW2011_GWF2_39_15] | (Brown et al., 2015) |
| OHA80798.1 | hypothetical protein A2675_01620 [Candidatus Yonathbacteria bacterium RIFCSPHIGHO2_01_FULL_51_10] | (Ananthar aman et al., 2016) |
| OHA83705.1 | hypothetical protein A2408_00355 [Candidatus Yonathbacteria bacterium RIFOXYC1_FULL_52_10] | (Ananthar aman et al., 2016) |
| KKS94036.1 | putative steroid-binding protein 3 [Candidatus Collierbacteria bacterium GW2011_GWC2_43_12] | (Brown et al., 2015) |
| KKU89055.1 | hypothetical protein UY18_C0004G0020 [Microgenomates group bacterium GW2011_GWF2_47_9] | (Brown et al., 2015) |
| OIP98122.1 | hypothetical protein AUK40_01915 [Candidatus Wirthbacteria bacterium CG2_30_54_11] | (Probst et al., 2017) |
| PJA14771.1 | hypothetical protein COX64_01640 [Candidatus Dojkabacteria bacterium CG_4_10_14_0_2_um_filter_Dojkabacteria_WS6_41_15] | (Probst et al., 2018) |
| EKD84736.1*<br>HCE30597.1* | Predicted heme/steroid binding protein [Candidatus Daviesbacteria bacterium]* | (Wrighton et al., 2012) |
| OGE21801.1 | hypothetical protein A2778_04905 [Candidatus Daviesbacteria bacterium RIFCSPHIGHO2_01_FULL_40_24] | (Ananthar aman et al., 2016) |
| KKQ43725.1 | hypothetical protein US62_C0043G0008 [Candidatus Woesebacteria bacterium GW2011_GWA1_37_8] | (Brown et al., 2015) |
| KKR31495.1 | hypothetical protein UT63_C0068G0005 [Candidatus Gottesmanbacteria bacterium GW2011_GWC2_39_8] | (Brown et al., 2015) |

\* EKD84736.1 and HCE30597.1 are identical. Protein name and species are provided for HCE30597.1.

**Table A2.** Crystallography statistics.

|  |  |
| --- | --- |
| Resolution range | 19.28-1.9 (1.968-1.9) |
| Space group | P 63 |
| Unit cell | 72.17 72.17 22.81 90 90 120 |
| Total reflections | 445959 (41994) |
| Unique reflections | 5544 (519) |
| Multiplicity | 80.4 (80.9) |
| Completeness (%) | 99.87 (100.00) |
| Mean I/sigma(I) | 49.00 (4.27) |
| Wilson B-factor | 25.28 |
| R-pim | 0.09944 (0.3724) |
| CC1/2 | 0.968 (0.643) |
| CC* | 0.992 (0.885) |
| Reflections used in refinement | 5544 (519) |
| Reflections used for R-free | 280 (24) |
| R-work | 0.2228 |
| R-free | 0.2702 |
| Number of non-hydrogen atoms | 663 |
| Macromolecules | 599 |
| Ligands | 43 |
| Solvent | 21 |
| Protein residues | 76 |
| RMS(bonds) | 0.021 |
| RMS(angles) | 1.89 |
| Ramachandran favored (%) | 97.30 |
| Ramachandran allowed (%) | 2.70 |
| Ramachandran outliers (%) | 0.00 |
| Rotamer outliers (%) | 0.00 |
| Clashscore | 4.77 |
| Average B-factor | 29.60 |
| Macromolecules | 29.26 |
| Ligands | 31.56 |
| Solvent | 35.27 |

**Table A3.**
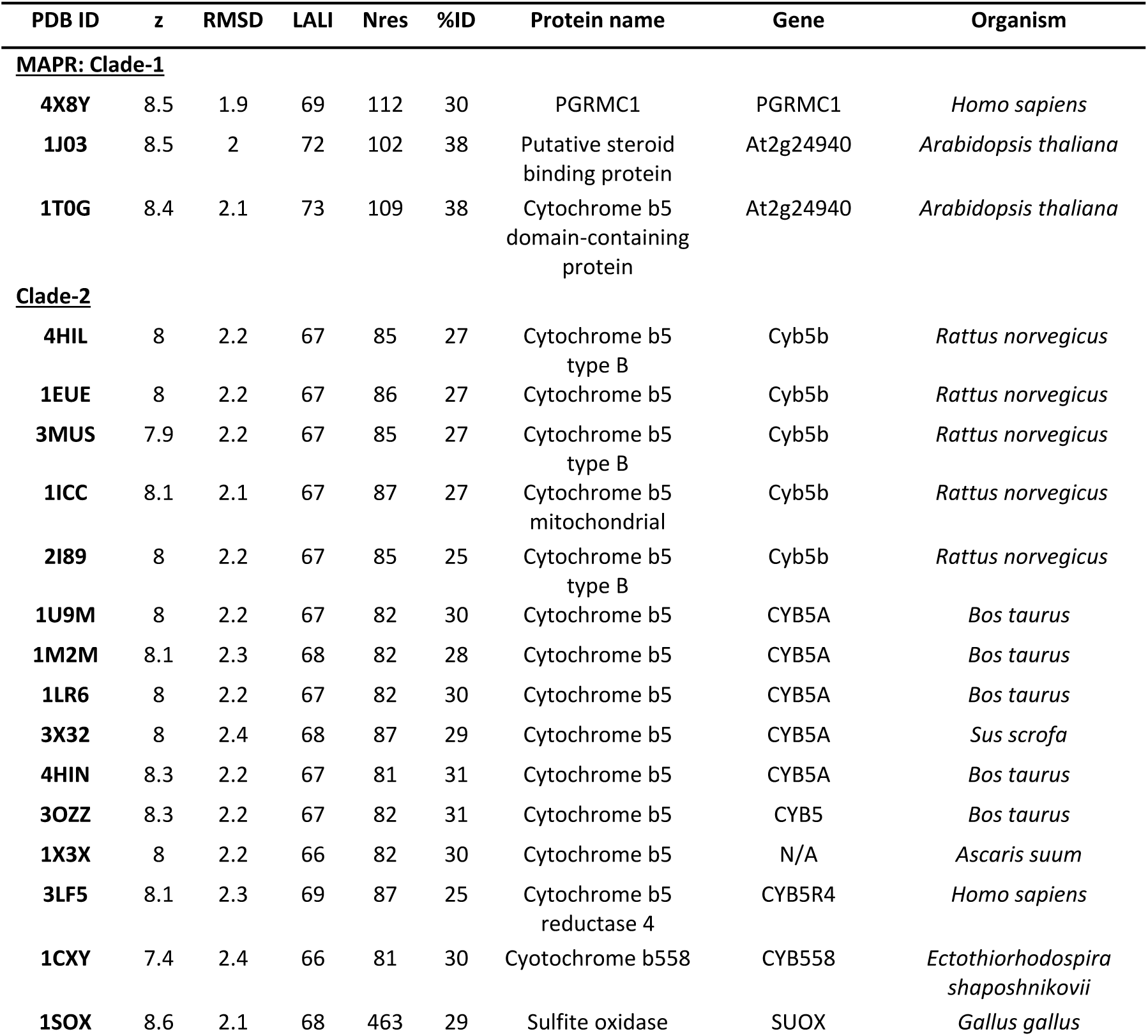
Proteins with the highest structural homology to archaeal Hadesarchaea YNP_N21 cytb_5M_ (KUO41884.1, PDB accession number 6NZX). The protein structures are those from Fig. A6. 3MUS is the Ref_RatCyb5 sequence of Fig. A5. z: z-score confidence in similarity significance, RMSD: root-mean-square deviation of atomic positions across the aligned sequences (Å), LALI: Total number of aligned residues, Nres: Total number of residues, %ID: percentage sequence identity.

Supplementary Appendix A. Zip archive containing full PDF images, FASTA sequence alignments, and xml tree files for preliminary studies leading to and including Fig. A1B.

Supplementary Appendix B. Zip archive containing top BLASTp hits from eukaryotes, using each respective cytb_5MY_ protein from Fig. A1B as query sequence in BLASTp against the eukaryotic data base. See the methods section and the README file in the zip archive for details.

Supplementary Appendix C. Sequences, alignments and phylogenetic trees for the results summarised in Fig. A4, Fig. 2 and Supplementary Appendix D.

Supplementary Appendix D. Summary table of the phylogenetic analysis of Clade 1 topologies for Fig. A4 using TBE branch support values analysis.

Supplementary Appendix E. Zip archive containing full PDF images, FASTA sequence alignments, and xml tree files for Fig. A9.

Supplementary Appendix F. 533 sequences of Cyp51A from Fig. 6.

Supplementary Appendix G. Protein Data Bank validation reports for the structure of Hadesarchaea YNP_N21 cytb5M (KUO41884.1), with PDB accession number 6NZX.

