## Supplementary Appendix A for "MAPR origins reveal a new class of prokaryotic cytochrome b_5_ proteins and possible role in eukaryogenesis": Fig1A_20180702_168seq_Input.docx

>PGRC1__BILA_Homo_sapiens@NP_006658_1

MAAEDVVATGADPSDLESGGLLHEIFTSPLNLLLLGLCIFLLYKIVRGDQPAASGDSDDDEPPPLPRLKRRDFTPAELRRFDGVQDPRILMAINGKVFDVTKGRKFYGPEGPYGVFAGRDASRGLATFCLDKEALKDEYDDLSDLTAAQQETLSDWESQFTFKYHHVGKLLKEGEEPTVYSDEEEPKDESARKND

>PGRC1__BILA_Mus_musculus@NP_058063_2

MAAEDVVATGADPSELEGGGLLHEIFTSPLNLLLLGLCIFLLYKIVRGDQPGASGDNDDDEPPPLPRLKRRDFTPAELRRFDGVQDPRILMAINGKVFDVTKGRKFYGPEGPYGVFAGRDASRGLATFCLDKEALKDEYDDLSDLTPAQQETLSDWDSQFTFKYHHVGKLLKEGEEPTVYSDDEEPKDETARKNE

>PGRC1__BILA_Monodelphis_domestica@XP_001372255_2

MAAEEVAPGEVEGEAGGLLQEIFTSPLNLLLLGLCFFLLYKIVRGEQPPTAGAGDEEPPVLPPLKRRDFTLAQLRRFDGTQDPRILMAINGKVFDVTKGRKFYGPEGPYGVFAGRDASRGLATFCLDKEALKDEYDDLSDLNATQQETLKDWESQFTFKYHYVGKLLKEGEEPTKYSDDEGPKDARERKDD

>PGRC1__BILA_Aplysia_californica@XP_005093157_1

MADSTVNDAADGSLVMRIFSELFGSPLNLALLGVCGFLIYKIIASRRSTDTLPPKEPELPPLKKQDFTLEQLREFDGKGNDGRVLIAVNGKVFDVTKGKRFYGPGGPYGLFAGHDASRALATFSLGEDALKNEYDDLSDLNSLQMESVREWEMQFQEKYVYVGKLLKPGEQPTEYSDTEDEEKTETDKKAD

>PGRC1__BILA_Aedes_aegypti@XP_001648600_1

MADEKVIPETAGTAEEQQIGLLTSVIHEIIYSPLNLILVGVITILVYKIFRSQKQPSAGPPPEPELPRLRRDFTVAELLQYDGKQPDGRVLVAVNGSVYDVTKGKRFYGPGGPYAAFGGRDASRGLATFSVTSNDNAEYDDLSDLTPMEMESVREWEAQFKEKYILVGRLLKPGEKPTNYSDEDEDTPNESTPTGDNATTTTSTSTSTATKEETPAAAPSSPKKTVDSSSSGDAKLIDFTTETSPATTGEVPSKAEDLKTGGDE

>PGRC1__BILA_Drosophila_melanogaster@NP_573087_1

MDSKVGADPSSAYDKEIGNNLNNDDSSFLGNIIREILYSPMNLALLAIICFLVYKIVRDRTEVPSVGVAKPSEPELPKIRRDFTVKELRQYDGTQPDGRVLVAVNGSVYDVSKGRRFYGPGGPYATFAGRDASRNLATFSVVSIDKDEYDDLSDLSAVEMDSVREWEMQFKEKYELVGKLLRKGEEPTNYDDDEDEENVNNDEQERKNLPKSKTETDDTKQEKESHILQSNVTNKNNAQDQATPSTDC

>PGRC1__CNID_Scypho_Periphylla_periphylla__38418

MDNFNSYWKEFSAYFTAPVQICLIVVAAYFIRRILRGRREPEEPVVEEKHEALEPMKKRDFYLEEVKEYDGTKSKRILLAVNGKVFDVTREKSFYGPGGPYGVFAGQDASRGLATFSVDQSILKDEYDDLSDLNAMQMDSLREWEMQFVEKYQQVGKLLRPGQKRTPYELESETDDESTRNAAKKKKVPT

>PGRC1__CNID_Hydro_Liriope_tetraphylla__21877

MAQETATMMSSITQYFSTPVQVALLVIVIYLIRQLWRNKTEDEDEYEENISLEPMERQDFTLEELRPYDGKTNPRILVAVNGKVFDVSPRRNFYGPDGPYSTFAGRDASRSLATFSIDASCIKEEYDDLSDLNSMQMDQVQEWELQFMEKYEQIGKLLKPGEPHTDYKGNQESEDERTDGKSEIKSDAAKKDE

>PGRC1__BILA_Schistosoma_japonicum@CAX73419_1

MGREYPNEGNYIVAMVDAIKCPINLTLFLICVLLGYLSLRPLLRRNKSQKSLVPKMGKRDFTLEELQNFDGSGEHKRILLAVNGKIFDVTNKGQEFYGKGAPYAAFAGKDASRALACFNLETKDEYDDLSDLTADQMKTLREWELQFSERYDHIGRLLKPGEPHRIYEINDGDDDVGVNHVQGTLKEKLA

>PGRC1__PORI_Homo_Oscarella_pearsei__42261

MLESLSELFSLWNIVIFAVSFFVIKKMFLQPALPPSPPSPQEPSLPKMKPRDFTVEQLREFDGSDPNKPLCLAVNDKVFDVSRGRSFYGPGGPYAVFAGRDATRAFANFIADESQLRDEYDPCDDLTPGQRMQVNEWEDRFMMKYDYVGKLLPPSTEAGSDSGGDGEAETEENREALSRKVDALVSEAEKLAEEDMKGTSIGGLIDLISEDAKED

>PGRC1__PORI_Homo_Corticium_candelabrum__219013

MVFDILSDVWESLTVFNAFISLLCFYFLIRLFSSRGKTPQVATQQQEEEKPRMPPRDFTNEELLEFNGSDPEKPICLAVRGKVFDVTRGKDFYGPGGVYGVFAGHDATRGLANWSTDESDIKDTYDNCSDLTPVQWMEVRDWEGHFMSKYDYVGKLLTPEQAAERTTDDASLKERKELFDKVQSLVKQADEEAMKQLENETKPKDKDT

>NENF__BILA_Homo_sapiens@NP_037481_1

MVGPAPRRRLRPLAALALVLALAPGLPTARAGQTPRPAERGPPVRLFTEEELARYGGEEEDQPIYLAVKGVVFDVTSGKEFYGRGAPYNALTGKDSTRGVAKMSLDPADLTHDTTGLTAKELEALDEVFTKVYKAKYPIVGYTARRILNEDGSPNLDFKPEDQPHFDIKDEF

>NENF__BILA_Mus_musculus@NP_079700_1

MARPAPWWRLRLLAALVLALALVPVPSAWAGQTPRPAERGPPVRLFTEEELARYGGEEEDQPIYLAVKGVVFDVTSGKEFYGRGAPYNALAGKDSSRGVAKMSLDPADLTHDTTGLTAKELEALDDVFSKVYKAKYPIVGYTARRILNEDGSPNLDFKPEDQPHFDIKDEF

>NENF__BILA_Monodelphis_domestica@XP_001374471_1

MAAPGLRRRPNLWPFLQALVLALNLAAVRAQQTPRTPERGPPVRLFTEEELARYRGEEEDQPIYLAVKGVVFDVTSGKEFYGRGAPYNALVGKDSTRGVAKMSLDPADLTHDTTGLTEEELKSLDDIFTNVYKAKYPIVGYTARRILNEDGSPNRNFKPEDQPHFDIKDEF

>NENF__CNID_Antho_Acropora_digitifera@XP_015766270_1

MENCFATFALMFSFLAVACTALDVTVESVENHSPFSEQTDRSRSNENTVFTKEELAKYNGEDPSLPVYVAIKGIVFDVSEAREFYGAGGKYNVFAGRDASRALAKWSMSEEDMNDNLDDLTEEELARLDESYDKLFASKHTVVGYLEGHELQDDDNLSRRMFNDNIEL

>NENF__CNID_Antho_Pavona_decussata__61800

MENCFATFALMFSFLAVACTALDVTVESVENHSPFSEQTDGSRSNENTVFTKEELAKYNGEDPSLPVYVAIKGIVFDVSEAREFYGAGGKYNVFAGRDASRALAKWSMSEEDMNDSLDDLTEEELARLDESYDKLFASKHTLVGYLEGHELQDDDNFSRKMFNDNIEL

>NENFlike__FUNGI_Mortierella_verticillata@07310T0

MSSIYAYFESLAKEPYFLPSIVISVSIALLITYLQDKPKNNDDSQSTDNKPTASSDPQQPATATDAAPKPKAFALPTTELPPPNTTRVFTHEELAKHDGKDPEAPIYVAIKGTVFDVSAKRAMYGPGAGYSCFAGKDASKALGKSSLKIEECVADYSDLTEKELKVLNDWYSFFEKRYPIVGKTEA

>pdb_MAPR_1T0G_A Chain A, cytochrome b5 domaincontaining protein (Arabidopsis)

MGHHHHHHLEEFTAEQLSQYNGTDESKPIYVAIKGRVFDVTTGKSFYGSGGDYSMFAGKDASRALGKMSKNEEDVSPSLEGLTEKEINTLNDWETKFEAKYPVVGRVVS

>pdb_MAPR_1J03_A Chain A, putative steroid binding protein (Arabidopsis)

GPMEFTAEQLSQYNGTDESKPIYVAIKGRVFDVTTGKSFYGSGGDYSMFAGKDASRALGKMSKNEEDVSPSLEGLTEKEINTLNDWETKFEAKYPVVGRVVS

>NENFlike__FUNGI_Spizellomyces_punctatus@08863T0

MATPLHKQASPEVVTPFGPFLDISSWNPVVLGAVLGLIVLVLVKSLTRGQSKVPQSPKKPQSPQSVTSSAPVGMMTTDSGPTQPPGDKEFTAAELARYDGTNAELPVYLAVKGTVFDVTKNRDMYSPGKGYHVFAGKDASQALGKSSLKPEECVPDYSSLTAEEMETLNKWEAHYKKKYNIVGKVVD

>NENFlike__FUNGI_Catenaria_anguillulae@56261

MAAVTPLFPDIPLPLLLAVFFGSISALFFLRSRSRSSSSPNPAASATSATQPAKSQASPPNMTTSNDPPATQLTLTPAQLTQFNNADASLPVYVAIKGIIYDVSSNREMYPCGQSYGLFAGKDASRALAKSSLDEKHIRADWQDLPEEEIKVLDDWVKFYEKKYPKVGRVVAE

>NENF__BILA_Acanthaster_planci@XP_022083121_1

MELQNLSNLLPSFIILYCLLVCMIFPLSSSSEKDKDDDKDLNLNFQERPMRAFTDEELSFYDGSFEGRPIYMAVKGVVFDVTSGKRYYGKGAIYNVLAGKDCTRAVAKMSLDPQDLTSDVTGLLEEELASLDEVFREVYMAKYPVVGHMSFSALETDRDSHGRTEL

>NENFlike__ICHTHYOSPOREA_Creolimax_fragrantissima@0714T1

MQAIGLPDFETHWYVVLVPFALAIYFALRVSSNNFTTCPYIITGKPRTSAEKAMEKAEQNAFKAGAVRHFSAEELRHFDGTDENSPVYVAIKGIVYDVSENRDAYKPGKGYALFAGRDASRALALMSLKVDDATANLDGLEPKQMDVLDNWVSFYKKKYKVMGTTGVTYSAHEKNVRDATESAKIK

>NENF__CNID_Antho_Antipathes_caribbeana__52569

PARGGICIGRLVLAIVLVAFSNGEKVSIDDGAHSPFPDDFKENADLLHRIFTKSELATFDGKNAGPIYMAVKGIVFDVTSAREFYGPGHTYHKLTGKDASRAIATWSMEEDELNDNLDGLTTEQLNSLENIFNTIYMAKYPVVGYLQGHEPVEENTTFRRRKVEL

>NENFlike__ICHTHYOSPOREA_Sphaerothecum_destruens@comp20708_c0_seq5_6

FLESVFCEDVAKIYHNWIAFSCIVAWKYFPTDIKDDFFLRLAKVQPNSLLFYLFGLAATSYVCFRVFTHSSKTQTLVFTKDLLAKYDGTNPNLPVYLAIQGVVFDVTRNRDMYAPGNSYSIFAGKDASYALATSSLDSKDAHCNLEQLTDKQRKTLQDWEDYYKKRYPIIGKLEPSGSVQ

>NENFlike__ICHTHYOSPOREA_Amoebidium_parasiticum@34200

MELSWLYIAVPVLVGFILFLVRSSPPPTQHETEEEKPEPPREPDTPERVFTVEELKQYDGSTDLPVYVAVKGWVFDVSPRREMYAPGQGYAIFAGKDASRGLGMSSLTPADALSDFSTLSADHLKVLNDWEVYYRKRYACVGHVEGSHAASELAKGVGVKGEEEKKNE

>PGRC1__ICHTHYOSPOREA_Amoebidium_parasiticum@22593

MESVSELLSPINVFLVVVIAVLVQKILTPAPDYSKTVVLDAPKEKIEPRDFTLEELVKFNGTDDPHIFMGVSGKVYDVTARKQFYGPGGPYELFAGRDASRALAKGDLSGSSLSMQWDELDDLTGDERATLRDWVESFEAKYDVVGQLVKAHQPPAEPESQPETTKLPAPPGRSGSTTPTPPEGADWTMVNKDDIGASCADLTAHPVHQKS

>NENF__PORI_Calc_Clathrina_coriacea__44495

QASFAGAVKMRFDGVLFFSACLAVVSRLVHCDEAPTEPTLTTRWFSEAEIAEYDGSDPDKPIYMSVKGTVFDVTSGKSFYGKGEGYNALVGGDRSVSVARMSLEPEDLAKKHDLSVLSQDERKSLDEIYESLYMVKYPVVGKMDFVRDSEEQPLSSPPAADEHTEL

>PGRC1__PORI_Calc_Sycon_ciliatum@scpid78563

MAVDIIEELSTLLEPFNLFLIAVCGFLLYKIIFAPADPGKDVKKEPAVEQVVGDLTVEQLLAYDGKDPTKPVLLAVCGKIFDVTAGKHFYGPGGPYEVLAGHDATRALALMSNEHSAVKDVDDDLSDLTRGQLSTAREWEQSFQFKYRYVGALVKESDSGEGKDDEQAKTDDEPPSPSATPEPTGNEEASTDGADVSEDTNIPSDP

>NEUFC__AMOEBOZOA_Acanthamoeba_castellanii@470459504

MSEAKGASLRQRPGKKQKAKKDGGIEITRVEKKGAENKQEDIQRPVRPSKKPPSSSSSSTARIYETLAVVCLLLALLVGVAVYLLHYEPQIVKQLDEGFVKAAATIAQLFQPKNGPNATSKPVAKKAKVVSKIFSADELRLHNGSDPAKPILLSVLGKVFDVTDGKRFYAKGGSYSFFAGRDASRSFATGEFEEENLTDDVTDLEPEQVAAIKEWQTQFERQYKYLGKVVGRFYDAAGKATPALKKVKEKLQKAKIVVDQELEEKQQFPECNSRWTEKDGATVWCTESSGGIERSWKGYPRQFKSARAQNPSPDAFECQR

>NEUFC__PORI_Calc_Leucosolenia_complicata@lcpid21432

MSARHRKGAGLSSQLPSYEEDGSKSARSSTKPVLLILVAVIVLAGIGGFVLRKEISAVLDSQQDPDSTKSQEAHTASEAAADSEIKDAGADTPPETVKRSAKNAAEDRLFTVEELKQYDGSDSSKPIYLALVGEVFDVSKGKDYYGKDGGYAFFSGRDAARAFVTGEFNEEGLTDNVTGLSTPNIIGIVEWQAFYRKDYTYAGKLIGTYYDEQGNPTENLKAYTELVDIANGEKLSEEEERKMFPPCNSHWAQGKGGTVWCSKESGGIKRDWVGVPRRYYKAGSDEYRCVCIRTTGPPSLGEDRGADFGDLVNPNMRAFDYCHPESATCAIPEK

>NEUFC__PLAC_Trichoplax_adhaerens__24556

MAVTRRKAAKSDDSKTDKIEKPEETNQESNSKGSKLVPIFILAAAIAAGVFAYKQDVDLLANTAESYQNKYKDESKRPAVGTGSGSSTDLSNVIAPDGQIFTASDLKQYNGDDPQKPIYLAVLGQVFDVTKGKEHYGKGGGYNFFSGRDGTRAFVTGDFSEKGLNDEIKGLSHENFIGIKEWIDFYHKDYSYVGKLIGRFYNAKGEPAPILSKIKTRLSKAYALRKEAENENFKMPGCNTHWTKEEGTVYWCEKKSGGIERDWVGVPRKLFYENGKFKRCACIRTKGPSSFESSKNSNNGDLDYHLLKEYDGCAFASTTCSANDAE

>NEUFC__PLAC_Hoilungia_hongkongensis__2357

MPATKRKEAKTSKPTVSSEAKTNDADQKKDSKESKPKLLPIFILAVTITVGALAYSRNLDLPMNPDEARAEYEAQINNQDKYKDESKRPAAGTGSATNTDLSSIIADGGQIFTTSELKQYNGEDPEKPIYLALLGQVFDVTKGKEHYAKGGGYGFFSGRDGTRAFVTGDFTEKGLIDDIKGLTYENLIGIKEWIDFYRKDYSYVGKLIGRYYDSEGKPAAILSKIKTRLTKAYRLREEAQVENLKMPGCNTRWTKEEGTVYWCENKSGAIERDWAGVPRKLFSAEGKFKRCACIRTSGPSSYGSVKNTNNGDLGFHLLREYEGCNPTSNTCKANDAQ

>NEUFC__CNID_Antho_Balanophyllia_europaea__419706

MSRERNIKSAKKPEKLSTSDDVQSSSREEKEMKKDALASKSSGAQTFVQAYHVLMPVVVVIIAGLLTFIVTRQQTETTMKEATKVPQTATTKDNSKKQVLLTAEELAKHDGSDPNIPVYIVVLGRIYDVEKGRRHYEVGSGYNVFAGRDSTPSFVTGKFVREEATDDVKGLSPEEMMGIKDWLDFYRKDYTYVGKLIGRYYDSEGNPTEALKEAKAVIKEGQRLKKLQEAENKRFPGCNSKWSAGEGSVVWCSENSAGISRDWAGVPRKMFKPGKKDAKCVCVKTTGPSSDTGEGNEGDLNNPNMQLYPDCGKYDVTCKL

>NEUFC__CNID_Antho_Porites_lutea__252363

MSRERNIKSTKKPEKLNVSDYDDKTSAQMEKDVSVSKPSGVQTFAKIYYVLMPVLVVIIAGLLTFAVTRQETATIVKETTKETGTGTGGKTKETSKKEVLLTAKELAKHDGSDPNIPVYLAILGRIYDVEKGRRHYEVGSGYNVFAGRDSTPSFVTGKFAREDATDDVKGLSPEDMMGVKGWLDFYRKDYTYVGKLIGRYYDSEGKPTEALQEAKALIKEGQRLQKLQDAENKRFPGCNSRWSKDEGSLVWCSENSAGINRDWVGVPRRMFKPGKKDPKCVCVKATGPSSDTGEGNEGDLNNPNMKLYPGCGKYDVSCKL

>NEUFC__PORI_Homo_Oscarella_pearsei__41833

MSAKSEDKKEEKRPESKKRESKPSSTSIPRSTTILIVLMSAAIALLAISLFERKSDDSPASVEERKSPPDVGRQERKAEPSIPPKTTEKPPSEAPSQSKKDTGKRLFPVSANDKVFTVEELGEHGPNAKEIYLAIFGRVYDVTKGKKHYGKNGGYWFFAGKDATRSFVSGDFTDEGLTDDLTDLSPSLLLGIEEWVETYDKDYKYIGILAGRFYNSDRSVTQDLLDVEEKIKRGHVEKATENNLMKLFPSCNSHWSAGKGEVSCSKKSGGIERNWVGVPRKLFTAGGKRHRCACVKNFGPGSDGEDKGKKHGDLDDPRLKQYDGCPANANACKVGS

>NEUFC__PORI_Homo_Corticium_candelabrum__153723

MSKAHDVKRRRVDADGSRTESPDAGKPTDRQASHSAAASVETEPWGPGCMLLGVIAVGTLTLLVTFTLYRQQELGYSRSEEKATDVDDSVGADGHAKPLFPVYATDQLFTRDELSSHGSSSKRIHLAILGRVYDVTKGRKYYGPKGGYAFFAGRDATRSFVSGDFSDAGLTDDLEGLSIREYLGVADWVGTYEKEYKYVGRLIGRFYDGDGRPTDSLLDAENKIRLAGEMKSKEDDMKKLFPPCNSRWAAGKGEVWCSNKSGGIHRDWVGVPRKMFVPGGSAYRCACVKDSGAASDGSDTGGRGDLDNPQLNEYQNCNSKSATCAVDD

>NEUFC__PORI_Homo_Plakina_jani__20654

MSYDQNVKHRRSMKEGSREEVPFTSTSPSDEMTQQQAKPQYAGLFASGWYWLFAVATVAGLAVLLVNKQQADIEGHDTDLYSEVEEASKPPSNQERLFAVYPSDRLFTQDELSKLGSSAKRIHLAILGRVYDVTKGHKYYGPKGGYSFFAGRDATRSFVSGDFSDSGLTDDLEGLSTREFLGIADWVGTYEKDYTYVGRLIGRFHDQTGAPTDALLDAEKKIRLARELQAEEEDRNKLYPPCNSRWAAGKGEVWCSDKSGGIQRNWVGVPRKLFVPGGSAYRCACVKDSGSASDGSDTNGRGDLDNPQFKEYDNCNPKSTTCTVDD

>NEUFC__PORI_Demo_Crambe_crambe@TR29691_c0_g2_i1

MGLRGYLARIVFIVLVAIAFSYRNEIMEMISKAVGNRETSPPTTEAKTTGQDKPPPPPPPPTGQDKPTAGQNEPKPSGGHKTLKEPELDYYNPPFDPENPPEPLYSKSGGTRMVTLNELAAHGHSGPLKPIWLAMMGKVFDVDKGAEHYYGPNGGYKFFTGIDGTKAFVTGEFDEEGLTDDITDLTPLQLGEIENWVKFYGDDYTYVGKLIGRYYTRDGSPTKHWYHYQKKLGEKELIKAEQKRQEKQFPGCNSHWTESTKGKVYCSEKSGGITRDWVGYPRRYFAPGTKSWRCACVHEKDLGSPQVRMYPDCDPNSHECKVDPEKIKELKARGE

>NEUFC__PORI_Demo_Stylissa_carteri__92844

MVSRLSLLGVALILVAIGYQYRNDIQRLVSGATRTSPLKKEPPGQPEVSSVQTDIKPGQGTPPRTEPRGPTKNTATETPLQQSSEQSTPTSSKSNVAGSQTAETGAGPQKVDSKAYYNPPFDPEKGVQLFSKSGTRLITLNELAAHGHSGPLKPIWLAIVGKVFDVDKGAEHYYGPKGGYNFFTGRDGTRAFVTGEFDEDGLTDDLEGLSPLQLGEIENWVKFYNDDYTYVGKLIGRYYNKDGSPTKEWYKYNKLLGEQEKIKAEQKAQEKRFPGCNSKWTEQDGGNVYCSEKSGGIQRDWVGYPRRYFAPGSKTWRCACVNENDLDSPQVRLYPDCDPTSINCKINKEKIQQLRESGQ

>NEUFC__PORI_Demo_Mycale_phyllophila__132463

IGLQATGKWGKKKKVSPSNSTMAKPREKDQSEKSSRRPPPPPPARRNKYWGYIARIVFVVVVAICFSKRDVIKASLAKFSNQQAGGDNPEVRQADGDKPAVNPPLKPPEPDYYNPPFDPNNPPETIYSKSGVRMITKNELAAHGHSGPLKPIWLAMLGQVFDVDKGAEHYYGPDGGYNFFTGIDGTKAFVTGEFNEEGLIDDISELSPLQLGELDNWVKFYRSDYTYVGKLIGRYYNRDGSPTREWYAYQKKLGEKDLIKAEMRKMEEKFPPCNSYWDQATEGKVFCSTNSGGISRDWVGYPRRFFAPGTVDWRCACAHEDDLNDPKLKLYPDCDPKSSECKVIAEQIKALRKQRL

>NEUFC__PORI_Demo_Chondrilla_nucula__11553

MAGLSTRKQFALGVILVLFATLYYYHREGRINIGVFKTAENFISKLKDLVFPKSTSTTNRAHKTTRPSDTGAPVNTPPVPKARKLDNTGPDRRVPAPWPQDRAGVRLITKEELKRHGPNGPFSPVWLSIIGQVYNVEKGKEHYYGPNGGYSFFSGIDGTRAFVTGEFNKEGLVENIYDLTARDCKDLISWTVDTYDKDYIFVGKLIGGFYDAEGNPTEMRLELDRQAERAKDTEKLLKEEQLQFPSCNSRWNANDGGWVYCGKKSGGIHRDWEGLPRRYFSPITREERCACVNDKDLNHPKVNLYSNCAPDARECQTRPPGNLH

>NEUFC__PORI_Demo_Halisarca_caerulea__29695

MAGLSTRKQLAVGIVLVLFATLYYYHKEGRINIGIFKSIENFFINLLKPKATSTPRKTPPSTGSSSTASPSSSTSGGRKEPDVKARDLDTTGPDLRTPAPWPQDRRGVRLIAKEELKRHGPNGPFSPVWLSIIGQVYNVEKGKEHYYGPNGGYKFFSGIDGTRAYVTGEFNKEGLVENIYDLTARDCKDLLSWTVDTYDKDYIFVGKLIGGFYDKEGNPTEMRRELDRQAERAKDTERLLQEEQQQFPSCNSRWNPNEGGWVYCGKRSGGIHRDWEGLPRRYFSPITREERCACVHDRDLDHPKVNLYANCAPDARECQTRQPGSLN

>NEUFC__BILA_Caenorhabditis_elegans@NP_497868_1

MDKNRRRTDDAGLMTKTLAGIAALVFFLSFICSSYDITVTHVISTIDVILEESEYYRSAKEWTASSIDATWNEVSIPSRKAEHIQAINPEVDVAAGGKHVFTPEQLHFFDGTRDSKPIYLAILGRVYNVDGKKEYYGPGKSYHHFAGRDATRAFTTGDFQESGLIATTHGLSHDELLSIRDWVSFYDKEYPLVGVVADLYYDSEGQPTPELTDVLARVEKANEYRKAQAVEIEVFPPCNSEYNQNGGRVWCSTKSGGVERQWAGVPRKLIEPTTEKFRCACVKNFGPGVSGAEEVKSSSNRGDLDHPDLELFPDCSPTSNSCKIVS

>PGRC1__ICHTHYOSPOREA_Creolimax_fragrantissima@3431T1

MDEGGTLWDELTSPINIILTIAIVYMVKKLLESEDVYANPTVLDAPQPTIERDFTVSELKPYNGEEKPFILIGLNGNVYDVSARPDFYGLKGPYGLFAGRDASRALATGDLSGEALGDEFDDLTSLTEDERGAMIEWEQTYQMKYKRVGALVREHNPTTVSKSEKDVAEASKQCPESSSPTSTSSAKLAVNKAGLAATPSPSLSREWDIVDTDDLETNSQ

>ArchB _KXH77621.1 cytochrome B5 [Candidatus Thorarchaeota archaeon SMTZ183]

MVKFTKQELVLYDGKDGASAFIGFNGKVYDVSSSFLWQNGNHQVLHDAGCDLTDSLAEAPHGPEMLDRFPVVGILEDD

>ArchB_KUO39518.1 cytochrome B5 [Hadesarchaea archaeon YNP_45]

MRVFSREELSRYNGRNGAPAYVAHDGKVYDVTESFHWKGGRHHVLHDAGQDLTESMGRAPHPAELLKKFPVVGILRD

>ArchB_KUO41884.1 cytochrome B5 [Hadesarchaea archaeon YNP_N21]

MRVFTKEELSRYNGKEGAPAYVAYNGKVYDVTGSFHWKGGKHHVLHDAGQDLTESIGRAPHTAELLEKFPVVGVLRG

>ArchB_SES62969.1 Predicted heme/steroid binding protein [Methanococcoides vulcani]

MGVIEVEEFTKEELAKYNGKDGAKCYIAYQGEVYDVTDSMLWDDGDHQGMHEGGIDLTEEMDDSPHDDDVMEDFTVVGKLID

>ArchB_WP_091689031.1 cytochrome B5 [Methanococcoides vulcani]

MEEFTKEELAKYNGKDGAKCYIAYQGEVYDVTDSMLWDDGDHQGMHEGGIDLTEEMDDSPHDDDVMEDFTVVGKLID

>ArchB_WP_048206030.1 cytochrome b5 [Methanococcoides methylutens]

MEEFTKEQLAKYNGQDGEKCYIAYKGKVYDVTESMLWDEGDHQGMHEAGIDLTEEMDDSPHDDDVMEDFPVVGTLVD

>ArchB_WP_048195578.1 cytochrome b5 [Methanococcoides methylutens]

MEEFTKEELSKYTGKDGSKCYIAYKGKVYDVTDSMLWDDGDHQGMHEGGMDLTEEMDDSPHDDDVMEELPVIGTLIN

>ArchB_ArchBEu_MS_WP_011499504.1 cytochrome b5 [Methanococcoides burtonii]

MEEFTTEELAKYNGKDGEKCYFAYKGKVYDVTESMLWEDGDHQGMHEGGIDLTADHEDAPHDDDVLEDFPVVGTLKA

>ArchB_OBZ34268.1 cytochrome B5 [Methanohalophilus sp. DAL1]

MREFTPDGLAKYNGKDRDEIYVAYNGKVYDVTNSELWMAGDHQGMHEGGIDLTEEMEEAPHEEDVFNEFEIIGIFVNNH

>ArchB_WP_105460305.1 cytochrome B5 [Methanohalophilus euhalobius]

MREFTPDGLAKYNGKDRDEIYVAYNGKVYDVTNSELWMAGDHQGMHEGGIDLTEEMEEAPHEEDVFNEFEIVGIFVNNH

>ArchB_WP_096711997.1 cytochrome B5 [Methanohalophilus euhalobius]

MKEFTPDGLAKYNGKDRDEIYVAYNGKVYDVTNSELWMAGDHQGMHEGGIDLTEEMEEAPHEEDVFNEFEIIGIFVNNH

>ArchB_WP_072360653.1 cytochrome B5 [Methanohalophilus portucalensis]

MREFTPDGLAKYNGKDRDEIYVAYKGNVYDVTNSELWMAGDHQGMHEGGIDLTEEMEEAPHEEDVFNEFEIIGIFVNNH

>ArchB_WP_072561339.1 cytochrome B5 [Methanohalophilus halophilus]

MREFTPDGLAKYNGKDRDEIYVAYKGNVYDVTNSELWMAGDHQGMHEGGIDLTEEMEEAPHEEDVFNEFDIIGIFVNNH

>ArchB_WP_013038189.1 cytochrome b5 [Methanohalophilus mahii]

MREFTPDGLAKYNGKDRDEIYVAYKGNVYDVTNSELWMDGDHQGMHEGGIDLTEEMEEAPHEEDVFNEFEVIGILVNNH

>ArchB_WP_013897393.1 cytochrome b5 [Methanosalsum zhilinae]

MGSDMKEFTAEELAKYNGRDGNRAYVAYNGKVYDVTDSFLWEDGDHQGMHEAGMNFDDELDLEAPHEADVMDDFPVVGTFRENSD

>ArchB_WP_091710816.1 cytochrome B5 [Methanolobus vulcani]

MKEFTLEEVAMYNGTDNEKVYVVYAGQVYDVSESEFWESGEHMGLHEAGTDLTESLDLEAPHEVDALDNFPIVGKIKE

>ArchB_WP_023844018.1 heme/steroid binding protein [Methanolobus tindarius]

MQEFTLEEVAKYNGTDNEKVYVVYAGQVYDVTASEFWESGEHMGLHEAGTDLTESLDLEAPHETDALDNFPVVGKIKE

>ArchB_WP_091934094.1 cytochrome B5 [Methanolobus profundi]

MQEFTIEEVAKFNGKDNEKIYVVYSGNVYDVSESDFWDDGEHMGLHEAGTDLTESLDLEAPHEVDALESYPIVGTIKK

>ArchB_WP_094228323.1 cytochrome B5 [Methanolobus psychrotolerans]

MQEFTLEEVAKFNGKDGEKVYVVYAGQVYDVSGSEFWNGGEHMGLHVAGTDLTEALDMEAPHEIDALENYPVVGTIKK

>ArchB_WP_015054822.1 cytochrome b5 [Methanolobus psychrophilus]

MEEFTLEEVAKYNGKNGQKAYVVYSGKVYDVTDSDFWDSGEHMGLHEAGMDLTEDLDMESPHESDALENFKIVGVIKK

>ArchB_AKB45585.1 hypothetical protein MSVAZ_3316 [Methanosarcina vacuolata Z761]

MAVLFGIFLAIGCAGNKPVTPNETGNSEQAVAPAGAITEAVNETPITEITLAEAVSPTERQNATEMQNVTGVSELKEYTLKELAEYNGKNGTVYVAYQGQVYNVSDSDIWKNGTHNGCNAGTDLTGKMDKTPHGGKILKGYPVVGTLKK

>ArchB_WP_048124311.1 hypothetical protein [Methanosarcina vacuolata]

MRKALIVLAVLFGIFLAIGCAGNKPVTPNETGNSEQAVAPAGAITEAVNETPITEITLAEAVSPTERQNATEMQNVTGVSELKEYTLKELAEYNGKNGTVYVAYQGQVYNVSDSDIWKNGTHNGCNAGTDLTGKMDKTPHGGKILKGYPVVGTLKK

>ArchB_WP_048158449.1 hypothetical protein [Methanosarcina sp. Kolksee]

MRKALIVLVVLFGIFLAIGCAGNKPVTPNETGTSERAVTSAEVVTGTEVVKETPVTEISPAEAVTSTLKNTTERQNATEMQNVTGVSELKEYTLKELAEYNGKNGTVYVAYQGQVYNVSDSDIWKNGTHKGCDAGTDLTGKMDKTPHGAKILKGYPVVGTLKK

>ArchB_WP_011306307.1 hypothetical protein [Methanosarcina barkeri]

MRKALIVLVVLFGIFLAIGCAGNKPVTSNETGTPGQAVTPAEDITEAVKETPVTEISPSEAVTPTKIQTATEMQNVTGIAGLKEYTLKELAEYNGENGTAYVAYQGQVYNVSDSNIWKDGTHKGCNAGTDLTGKMDKAPHGAAILKGYPVVGTLKK

>ArchB_WP_048120212.1 hypothetical protein [Methanosarcina barkeri]

MRKALIILVVLFGIFLATGCAGNKQVTPNETGTPEQAVTPAEGITEAVKETPVTEISPVEEGTPTIKNTTERQTAIEMQNMTRKSELKEYTLKELTEYNGENGTAYVAYQSQVYNVSDSDIWKNGTHKGCNAGTDLTGKMDKTPHGAAILKGYPVVGTLKK

>ArchB_WP_048154356.1 hypothetical protein [Methanosarcina barkeri]

MRKALIILVVLFGIFLATGCAGNKQVTPNKTGTPEQAVTPAEGITEAVKETPVTEISPVEEGTPTIKNTTERQTAIEMQNMTRKSELKEYTLKELTEYNGENGTAYVAYQSQVYNVSDSDIWKNGTHKGCNAGTDLTGKMDKTPHGAAILKGYPVVGTLKK

>ArchB_AKB82764.1 hypothetical protein MSBR3_2186 [Methanosarcina barkeri 3]

MGEIVRKALIILVLLFSTFLAIGCSGNEPVVPNETGTPNETGTPEEVVTPIEVSTPTGEKAIERQNTTETQNVTDKSKTKEYTLEELAEYNGKNGVAYVAYEGQVYDVSDDDLWKNGSHKGCNAGTDLTGKMDKTPHGAKILKGYPVVGTLKK

>ArchB_WP_048110418.1 hypothetical protein [Methanosarcina barkeri]

MRKALIILVLLFSTFLAIGCSGNEPVVPNETGTPNETGTPEEVVTPIEVSTPTGEKAIERQNTTETQNVTDKSKTKEYTLEELAEYNGKNGVAYVAYEGQVYDVSDDDLWKNGSHKGCNAGTDLTGKMDKTPHGAKILKGYPVVGTLKK

>ArchB_WP_095645751.1 hypothetical protein [Methanosarcina spelaei]

MRKALIILVLLFSIFLAIGCSGNEPGVPNETGTPGEVVTPIEVSTPAGETAIEIQSTTETQNVTDKSKTKEYTLEELAEYNGKNGIAYIAYEGQVYDVSDNDLWKNGSHKGCNAGTDLTGKMDKTPHGAKILKGYPVVGTFKN

>ArchB_WP_011023631.1 cytochrome b5 [Methanosarcina acetivorans]

MKEYTLEELSEYNGKNGKIYIAYDGQVYDVSDSYMWEDGTHQGLHDSGQDLSDAMDNEAPHGPEVFKDYPVVGTLKK

>ArchB_WP_048173635.1 cytochrome b5 [Methanosarcina siciliae]

MKEYTLEELSEYNGKNGKTYIAYAGQVYDVSDSYMWEDGTHQGLHDSGQDLSDAMDNEAPHGPEVFKDYPVVGTLKK

>ArchB_WP_048135733.1 MULTISPECIES: cytochrome b5 [Methanosarcina]

MKEYTLEEISEYNGKNGKVYIAYAGQVYDVSNSYLWEDGTHQGLHDSGQDLSEAMDNEAPHGPEVFKDYPVVGTLKK

>ArchB_WP_048137110.1 MULTISPECIES: cytochrome b5 [Methanosarcina]

MKEYTLEELSEYNGKNGKVYIAYAGQVYDVSNSYLWEDGTHQGLHDSGQDLSEAMDNEAPHGPEVFKDYPVVGTLKK

>ArchB_WP_048173339.1 cytochrome b5 [Methanosarcina sp. 2.H.A.1B.4]

MKEYTLEELSEYNEKNGKVYIAYAGQVYDVSNSYLWEDGTHQGLHDSGQDLSEAMDNEAPHGPEVFKDYPVVGTLKK

>ArchB_WP_048126521.1 MULTISPECIES: cytochrome b5 [Methanosarcina]

MKEYTLEELSEYNGKNGKVYIAYAGQVYDVSDSNLWEGGTHQGLHDSGQELSEAMDNEAPHGPEVFKDCPIVGTLKK

>ArchB_WP_015411227.1 MULTISPECIES: hypothetical protein [Methanosarcina]

MKEYTLEELSEFNGKNGKTYVVYDGQVYDVSNSYLWEDGTHQGLHESGKDLTEDMDEAPHGPEVFKDYPVVGTLKK

>ArchB_WP_048041697.1 cytochrome b5 [Methanosarcina mazei]

MKEYTLEELSEFNGKNGKVYIAYDGQVYDVSDSYLWEDGTHQGLHESGEDLTEAMDEAPHGPETFKDFPVVGTLKK

>ArchB_WP_048127057.1 cytochrome b5 [Methanosarcina lacustris]

MKEYTLKEISELNGKEGKVYIAYDGQVYDVSDSYLWEDGTHQGLHDSGQDLTEAMDEAPHGSEVVKDYPVVGTLKK

>ArchB_WP_048177937.1 cytochrome b5 [Methanosarcina sp. MTP4]

MKEFTPEELSEYNGKDGKMYVAYEGKVYDVSDSYLWEDGNHQGLHDAGQDLSESMNDEAPHGPEVFEDYPVVGTLKE

>ArchB_WP_011033516.1 hypothetical protein [Methanosarcina mazei]

MKKALIILIVLFGVFLAIGCAGEEERAPNETRAPEQTVTPAEAVTPTGEEETEIQTVTEMKEYTLEELAEYDGRNGKTYVAYQGQVYDVSNSDLWENGTHKGTHNAGKNLTEEMDDAPHGPEELKDYPVVGTLRE

>ArchB_AKB56454.1 hypothetical protein MSBRM_3456 [Methanosarcina barkeri MS]

MGKIMRKKLIILIVLFCMFLATGCVGNKLVTPEKAGTQADADTPNGEEATETQNLTEKTEIKEYTLEELAKYDGTNETIYIAYQGQVYDVSSDSYLWKDGNHEGCPAGKDITEELDRTPHGAEILKKYPIVGTLKE

>ArchB_WP_011306210.1 hypothetical protein [Methanosarcina barkeri]

MRKKLIILIVLFCMFLATGCVGNKLATPEKAGTQDDADTPNGEEATETQNLTEKTEIKEYTLEELAKYDGTNETIYVAYQGQVYDVSSDSYLWKDGNHEGCPAGKDITEELDRTPHGAEILKKYPVVGTLKE

>ArchB_WP_048124350.1 hypothetical protein [Methanosarcina vacuolata]

MRKKLIILLVLCCMFLATGCVGNELVTPEKAVTQADADNPNGEDTTETQNVTEPGIKEYTVEELAKYDGTNETIYVAYQGQVYDVSSDSYLWKDDNHEGCPAGKDITEELDRTPHGAEILKKYPVVGTLKE

>ArchB_WP_048158468.1 hypothetical protein [Methanosarcina sp. Kolksee]

MRKKLIILLVLCCMFLATGCVGNELVTPEKAVTQADADTPNGEDTTETQNVTETGIKEYTVEELAKYDGTNETIYVAYQGQVYDVSSDSYLWKDGNHEGCPAGKDITEELDRTPHGAEILKKYPVVGTLKE

>ArchB_OEU41989.1 hypothetical protein BGV40_11960 [Methanosarcina sp. Ant1]

MRKKLIILIVLFCIFLAIGCVGNKTVTPEEAVTQADVATPTGEEATETLNMTEKTEIKEYTLEELAKYNGKNGTIYVAYQGKVYDVSSDSYLWKDGSHEGCTAGKDITEEMDKTPHGAEILKKYPVVGTLKK

>Bact_OGP58440.1 hypothetical protein A2162_07725 [Deltaproteobacteria bacterium RBG_13_52_11b]

MRELTYQELATFDGKEGRPVYIAFEGRVYDVSNSPLWETGLHMNRHPSGKDLTADISAAPHGPEVLERYPQIGSVARGASEELNHLPAVLQKIIGAFPVARRHPHPVFVHYPIALLMATSFFVLLHLLFQKPSFGLTAYYLLILGAVSTPFAVATGLLTWWVNYRLKLTFFIRRKIQLSIVLLAFGIILLVWRSSSPDVSHPLYYIMAFLLTPIVFLLGYYGGQMTFPIEKGKV

>Bact_OGP67081.1 hypothetical protein A2169_08180 [Deltaproteobacteria bacterium RBG_13_47_9]

MKEFSSEELSSFNGEDGSPLYIAFGGKVYDVSKSPLWSKGHHMNRHPSGKDLSGAISAAPHGPEILERYPQVGILKKEVPEELKHLPSLLQRLLRQFPMARRHPHPMLVHFPIAFLTAASLFTLLSFFFQNFYFEITGFYLLILGAIASPFAMATGFLTWWVNYRMKLTLYVKRKIQFSILLLIFEIILITWRSLNPELSNLLYLILMLMLTPVVTLLGYYGGRMSFPPET

>Bact_OGP72750.1 hypothetical protein A2V86_03695 [Deltaproteobacteria bacterium RBG_16_49_23]

MKEFTPEELLSFNGKEGKPVFIAFEGKVYDVSKSSLWSKGLHMNRHPPGKDLSGEISAAPHGPEVFERYPQIGILKKGQPEELKHLPPLLRNLLLKFPVARRHPHPMLVHFPIAFLMASSLFLLLYLLFNHAPFEQTGFYLFLLGAVSSPFAIGTGLLTWWVNYRLKLTHFVKRKIQLSIILLIFEIILILWRTSGPEVSGPVYFILIFSLAPLVMLLGYYGGQMTFPLEKL

>Bact_OGP96884.1 hypothetical protein A2157_00600 [Deltaproteobacteria bacterium RBG_16_47_11]

MKEFTSEELVTFNGKDGKPVYIAFERKVYDVSKSPLWSSGLHMNRHPSGKDLTGEISAAPHGPEVFERYPQVGILKKGPPEELKHLPPVLQDLLQRFPMARRHPHPMIVHFPLAFLMGSSLFILLHLLFRKPAFEITSFYLFILGAISSPFAMVTGLLTWWVNYQLKLTLFVKRKIQLSILLLILEIILVLLRGSNPAMTNPLYFILMILLTPIVGLLGYYGGQMTFPAEK

>Bact_OGP87057.1 hypothetical protein A2156_09250 [Deltaproteobacteria bacterium RBG_16_48_10]

MREITPEELLSFDGKEGRPVYVSFQGKVYDVTRSRLWASGSHMKRHPSGKDLTGEIAAAPHGSEVLEQYPQVGVLQGGSPEELKHLPPWLRSVLKRFPFARRHPHPMVVHYPIAFLMASSLFMLLYLLFKKTSFELTGYYLLLLGTISSPFAIGTGLLTWWINYRLKPNYFVKRKIQLSFVLLPLEIFLILWRGLNPYPPLQQVHPAYIVLMLMLIPVVGLLGHYGGQLTFPSEK

>Bact_AFV97178.1 hypothetical protein B649_04320 [Candidatus Sulfuricurvum sp. RIFRC1]

MQQFSEEELRKYNGKDGMPAYIAFKNQVYDVTSSKFWQEGTHFKKHFAGCDLTDAMAHAPHSDEVFENYPCIGQFVSPCSLTPENKKDRYRQWYSKYHPHPLIIHFPIALHYFSAFVDILFLDNPSAGYETAVFLSFLIATIAGFFALISGVFSWWINYDFSISKPFVIKLIGALFTLIVGLIPLGQKLLNPNVAFSTGVDGIIYHAVIFMTVISITIVGYYGGKITWGAKK

>Bact_OHD88731.1 hypothetical protein A3G19_09145 [Sulfuricurvum sp. RIFCSPLOWO2_12_FULL_43_24]

MQQFSEEELRKYNGKDGMPAYIAFKNQVYDVTSSKFWQEGSHFKKHFAGCDLTDAMAHAPHSDEVFENYPCIGQFVSPCSLTPENKKDRYRQWYSKYHPHPLIIHFPIALHYFSAFVDILFLDNPSAGYETAVFLSFLIATIAGFFALISGVFSWWINYDFSISKPFVIKLIGALFTLIVGLIPLGQKLLNPNVAFSTGVDGIIYHAVIFMTVISITIVGYYGGKITWGAKK

>Bact_OHD86007.1 hypothetical protein A2Y52_02065 [Sulfuricurvum sp. RIFCSPLOWO2_02_43_6]

MQQFSEEELRKYNGKDGMPAYIAFKNQVYDVTSSKFWQEGTHFKKHFAGCDLTDAMADAPHSDEVFENYHCIGQFVSPCSLTPENKKDRYRQWYSKYHPHPLIIHFPIALHYFSAFVDILFLDNPSAGYETAVFLSFLIATIAGFFALISGVFSWWINYDFSISKPFVIKLIGALFTLIVGLIPLGQKLLNPNVAFSTGVDGIIYHAVIFMTVISITIVGYYGGKITWGAKK

>Bact_DAB38524.1 TPA: hypothetical protein CFH83_05500 [Sulfuricurvum kujiense]

MQQFSEEELRQYNGKDGMPAYIAYKGQVYDVTSSKFWKEGTHFKKHFAGCDLSAEMENAPHSDEVFANYPCVGQFIPSFIKPPETKKERYRLWYAKYHPHPAAVHFPIALHYFSAFVDILFLANPSREYDTAVFLSFLIATVMGFLALLSGVFSWWINYDFSTSKPFVIKLIGALFTLIIGFVPLVQKLQNPDVAFSMGADGITYHAVIFITVISITIVGYYGGKITWGAKQ

>Bact_KPJ78889.1 cytochrome b5 [Deltaproteobacteria bacterium SG8_13]

MKEFDSKELAKYNGENGSPAYIAHKGKVYDVTGSKLWAAGTHMRRHRAGTNLTTDIQAAPHGPEILERYSQVGTLKEETAETAVALPKILAILLESNPFFRRHPHPMTVHFPIVFMLSTPFFNVLYLITGIKSFELTALHCLAGGILFTLVAMSTGVLTWWYNYLGKMLRPVAVKIPLTIVMLITGIVAFVWRINDPGVLDPLQGTGIVYLLLILSLAPLTAVIGWYGASMTFPIEKQ

>Bact_OGP92792.1 cytochrome b5 [Deltaproteobacteria bacterium RBG_16_54_18]

MKKITTKELSEFNGKEGKAVYIAHEGKVFDVSASKLWQGGIHMQRHHAGSDLTTDLQSAPHGPEVLERYPHIAGLEKEAVAEQRMPAMLARLLKRSPMLRRHPHPMTVHFPIVFMFATTMFTLLYLITGMQTFERTALHCLGAGLLFTPVAMATGYYTWWLNYLSRPMRAVTIKKWLAAILLCIEIIAFIWRISNPAVLAPLSIATAMYLILILLLFPLVVILGWLGASLTFPVKKE

>Bact_OHE16742.1 cytochrome b5 [Syntrophobacterales bacterium GWC2_56_13]

MKEFDPESLSRFSGKDGQPAYISHKGRIIDVSASKLWKTGLHMKLHAAGRDLTADISAAPHGPEVLDGYPQVGTLKKERADRSLPKPLETLLERFPVLRRHPHPMMVHFPIVFAISPALFYLLFRITAVNAFETSAFHCLGAGILFSVPAILTGYFTWWLNYQARPLLPVRIKILFSTLLVAVLLAAFLLRLLFPAAVASFAGAGILYILLLSALIPIVTVIGWYGASLTFPLERK

>Bact_OGR27611.1 hypothetical protein A2139_13265 [Desulfobacca sp. RBG_16_60_12]

MTEKPDQEREFTPAELAAGSGADGAPVLIAFRGKVYDVTGSGLWEGGGHMDLHRAGHDLTGEFPDAPHGEEVFQRYPQVGVLKEEAPIPEAPAPKPSAAREFWRRVVHRVPLLRRHPHPMVVHFPIVFMIAAPVFTLLSLITGVKSFEVTGFHCLGGGLLFTPVAMVTGWFSWWLNYESRWLRPVVVKLILSPVLLLVGAGAFLWRYQNPEILAQFPGGPSLVYLGLVCALFPLVGVIGACGAELTFPVNHE

>Bact_EKO39054.1 putative heme/steroid binding protein [Desulfovibrio magneticus str. Maddingley MBC34]

MSQNPEKRFTPQELAAFDGADGKPTYLAYNGVVYDVSASRLWKAGKHMNRHHAGGDMGLELSQAPHNPDVLERFPRVGLLDAPALKPEPAADRVPPWLAKCIKRFPMLKRHPHPMTVHFPIAFCTAAPLCLLLALVTGKQAFAAALPVLLGLALVFTPVAIVTGLFTWWLNYASARVTPIMIKLAATPVLFLALLWTFIESVKTPDILAEAGQHVGFVLVVLALMPIVSVIGWFGAALTFPPHDD

>ArchB_WP_052368481.1 hypothetical protein [Candidatus Methanoperedens nitroreducens]

MEHVNVLLIEDNPGDARLIKEMLIEAKNISFDIEWKDRLSSGLERITMGGVDVVLLDLILPDSPRGFDTFTRTQAQAPEVPIVVLTGLDDETFAINAVRRGAQDYLIKGKVDSNLLTRTIRYSIARKLGEERHFTAMELREFDGKEGRPAYAAFKGKVYDVSNSSLWRDGIHAGSHFAGTDLTENMLRATHGEEVLVKFHIVGELSPEKTFRQRLVQRIEGLHLHPILVHFSIAYSVAIPLLAFLYIFTEEITFEIASYYILVLGLLTAPLATLSGLFSWKVTYAEKMTKVFARKIIFAIALIVVTTVSFMLRTLYTDILTEVDLNYYTYLALLVSLAPIVTVLGYDGGKIVYS

>ArchB_WP_052368516.1 hypothetical protein [Candidatus Methanoperedens nitroreducens]

MDRIKVLLIEDNPGDARLIHEMLAEEKKILFDLEWRDRLSSGLERLAEGGIDVILLDLMLPDSRGFDTFTRTQAQAPEIPVVVLTGLSDEALAIRAVRKGAQDYLVKGKVDSNLLVRSILYAIARRLGEEKYFAIDDLKRFDGKEGRPAYIAFKGRVYDVSNSRLWINGLHLGAHHAGSDLTENMMGAPHGEEVFIKFHVVGELSHEEPFRDRLVRRIKHRVIKRKKGL

>Bact_CZQ97721.1 Hypothetical protein Tpal_2168 [Trichococcus palustris]

MKNNNKAKMITLLLGAGLLFGACGTTQATTTDKASTSAAASSAVPASKAASSSSTASSSSAQSNAKTFTLDELAKYDGKNGNDAYVAVAGIVYDVTHAQKWQNGNHYGVQAGTDLTTAINKSPHGSSVLEGLPIVGTLVN

>Bact_GBD34318.1 Soluble cytochrome b558 [bacterium HR35]

MGKKSIYIGLITVMIIVVGGLVFFKFTNKKSQINHNQVSNQFSLPSAISNNDLLETFGSTSTEILNNQQLEQNSAQPQEKLYTLEEVAQHNSKESCWTVIRDRVYDLTQWIDKHPGGSDKISALCGKDGTQAFENKHGGEEKPEKTLEQFEIGKLKQ

>Bact_SFI24987.1 nitrate reductase (NAD(P)H) [Bradyrhizobium sp. Gha]

MAGRTKQGHDGSSHSPVVQLWTPAAPAKGTRPVTRAEVVLHNTKDDCWVIIRGKVYDITAWAPHHPGGAGIARMYAGKEATAEFGDYHSAEAVAHMAHFCIGELVEAPAALP

>Bact_WP_018458797.1 hypothetical protein [Bradyrhizobium sp. WSM4349]

MRAVIVGAGMGGLMTALALRQSGAFASVYVYEQTKVPSTAGAGLNIPPNGARICRWLGVDLDGGDSKGPDGVIDGGRAAILESTRQFNADGSVTKRPFDHVTAAGDGAGFHHMHRLDLLMCLYKRVSDFGIDSGAPCPIAVHMDCRLTQLRQTAGEVLATFSNGRTATGELLVGADGINSATLQLAWPNSRPKRWTEVTCFRGLIPRTGVASLRKANGNPLDHNPINSFSMDRHRADRSGATTYWVRGGELLNVWIAHYEPESAAFEQEEGDWFPVSQQEIVREVGEAFAGHPSRDDLIALSGAIVRPTKWGLYDRDALETWVQGRICLLGDAAHPMLPTFGQGAAQSFEDAAALASAFALHQRDVPTALLHYERVRHYRATRFQLGSKFAFDHLRAKDTAEQKALLERLDERVSPAFAHDKRGGEDDSWIYAYDARKIGSELPAKRLGPWDFRRTAKVRYGGIKLWMPANPAKGTRRVTREEIALHNTQNDCWIIISGKVYDITEWAPHHPGGAGIARMYAGREATAEFGDYHSTEAVAHMANFCVGALVEN

>Bact_WP_085350873.1 monooxygenase [Bradyrhizobium canariense]

MRAVIVGAGMGGLMTALALRQSGVFASVDVYEQTKVPSTAGAGLNIPPNGARICRWLGVDLDGGDSKGPDGVIDGGRAAILESTRQFNADGSVTKRPFDHVTAAGDGAGFHHMHRLDLLMCLYKRVSDFGIDSGAPCPIAVHMDCRLTQLRQTAGEVIATFSNGRTATGELLVGADGINSATLQLAWPNSRPKRWTEVTCFRGLIPRTGVASLRKANGNPLDHNPINSFSMDRHRADRSGATTYWVRGGELLNVWIAHYEPESAAFEQEEGDWFPVSQQEIVREVGEAFAGHPSRDDLIALSGAIVRPTKWGLYDRDALETWVQGRICLLGDAAHPMLPTFGQGAAQSFEDAAALASAFALHQRDVPTALLHYERVRHYRATRFQLGSKFAFDHLRAKDTAEQKALLERLDERVSPAFAHDKRGGEDDSWIYAYDARKIGSELPAKRLGPWDFRRTAKVRYGGIKLWMPANPAKGTRRVTREEVALHNTQNDCWIIISGKVYDITEWAPHHPGGAGIARMYAGKEATAEFGDYHSTEAVAHMANFCVGALVEN

>Bact_WP_085385823.1 monooxygenase [Bradyrhizobium canariense]

MRAVIVGAGMGGLMTALALRQSGVFASVDVYEQTKVPSTAGAGLNIPPNGARICRWLGVDLDGGDSKGPDGVIDGGRAAILESTRQFNADGSVTKRPFDHVTAAGDGAGFHHMHRLDLLMCLYKRVSDFGMDSGAPCPIAVHMDCRLTQLRQTAGEVIATFSNGRTATGELLVGADGINSATLQLAWPNSRPKRWTEVTCFRGLIPRTGVASLRKANGNPLDHNPINSFSMDRHRADRSGATTYWVRGGELLNVWIAHYEPESAAFEQEEGDWFPVSQQEIVREVGEAFAGHPSRDDLIALSGAIVRPTKWGLYDRDALETWVQGRICLLGDAAHPMLPTFGQGAAQSFEDAAALASAFALHQRDVPTALLHYERVRHYRATRFQLGSKFAFDHLRAKDTAEQKALLERLDERVSPAFAHDKRGGEDDSWIYAYDARKIGSELPAKRLGPWDFRRTAKVRYGGIKLWMPANPAKGTRRVTREEVALHNTQNDCWIIISGKVYDITEWAPHHPGGAGIARMYAGKEATAEFGDYHSTEAVAHMANFCVGALVEN

>Bact_WP_085409789.1 monooxygenase [Bradyrhizobium canariense]

MRAVIVGAGMGGLMTALALRQSGVFASVDVYEQTKVPSTAGAGLNIPPNGARICRWLGVDLDGGDSKGPDGVIDGGRAAILESTRQFNADGGVTKRPFDHVTAAGDGAGFHHMHRLDLLMCLYKRVSDFGIDSGAPCPIAVHMDCRLTQLRQTAGEVIATFSNGRTATGELLVGADGINSATLQLAWPNSRPKRWTEVTCFRGLIPRTGVASLRKANGNPLDHNPINSFSMDRHRADRSGATTYWVRGGELLNVWIAHYEPESAAFEQEEGDWFPVSQQEIVREVGEAFAGHPSRDDLIALSGAIVRPTKWGLYDRDALETWVQGRICLLGDAAHPMLPTFGQGAAQSFEDAAALASAFALHQRDVPTALLHYERVRHYRATRFQLGSKFAFDHLRAKDTAEQKALLERLDERVSPAFAHDKRGGEDDSWIYAYDARKIGSELPAKRLGPWDFRRTAKVRYGGIKLWMPANPAKGTRRVTREEVALHNTQNDCWIIISGKVYDITEWAPHHPGGAGIARMYAGKEATAEFGDYHSTEAVAHMANFCVGALVEN

>Bact_WP_085394750.1 monooxygenase [Bradyrhizobium canariense]

MRAVIVGAGMGGLMTALALRQSGVFASVDVYEQTKVPSTAGAGLNIPPNGARICRWLGVDLDGGDSKGPDGVIDGGRAAILESTRQFNADGSVTKRPFDHVTAAGDGAGFHHMHRLDLLMCLYKRVSDFGMDSGAPCPIAVHMDCRLTQLRQTAGEVIATFSNGRTATGELLVGADGINSATLQLAWPNSRPKRWTEVTCFRGLIPRTGVASLRKANGNPLDHNPINSFSMDRHRADRSGATTYWVRGGELLNVWIAHYEPESAAFEQEEGDWFPVSQQEIVREVGEAFAGHPSRDDLIALSGAIVRPTKWGLYDRDALETWVQGRICLLGDAAHPMLPTFGQGAAQSFEDAAALASAFALHQRDVPTAMLHYERVRHYRATRFQLGSKFAFDHLRAKDTAEQKALLERLDERVSPAFAHDKRGGEDDSWIYAYDARKIGSELPAKRLGPWDFRRTAKVRYGGIKLWMPANPAKGTRRVTREGVALHNTQNDCWIIISGKVYDITEWAPHHPGGAGIARMYAGKEATAEFGDYHSTEAVAHMANFCVGALVEN

>Bact_WP_092226384.1 monooxygenase [Bradyrhizobium sp. Gha]

MRAVIIGAGMGGLMTALALRQSGVFTSVDVYEQTKVPSTAGAGLNVPPNGARLCRWLGVDLDGGDPNGPDGVIDGGRAAILESTRQFNADGTMTQRRFDHVTAAGDGAGFHHMHRLDLLMCLYKRVFEFGPDTGAACPITVHMDCRLTQLRQTAREVTATISNGRSTTGEVLVGADGINSATLQLAWPNPRPKRWTEVTCFRGLVPRAVVASLRKADGSPLDHNPIDSFSMDRHRTDRSGATTYWVRGGELLNVWIARYEPDSAAFEKEEGDWFPVSREEILHEVGEAFTGSASRDDLLALAGAIVRPTKWGLYDRDALETWVQGRICLLGDAAHPMLPTFGQGAAQSFEDAAALASAFALHKRDVATALLHYERVRHYRATRFQLGSKFAFDHLRARDTAEQKALLEKLDERVSPAFAHDKRGGEDDSWIYAYDARKIGAELPRKRLGPWDFRRAAKASYATITSKLWTPAAPPKAPRPVTREEVALHNTKEDCWIIVRGKVYDITEWAPHHPGGAGIARMYAGKEATAEFGDYHSAEAVAHMAHFCIGDLVDAPTTLP

>Bact_WP_092298662.1 monooxygenase [Bradyrhizobium sp. Ghvi]

MRAVIVGAGMGGLMTALALRQSGVFASVDVYEQTKVPSTAGAGLNIPPNGARICRWLGVDLDGGDPKGPDGAIDGGRAAILESTRQFNADGSVTKRPFDHVTAAGDGAGFHHMHRLDLLMCLYKRVFAFGPDSGEPCPITVHMDSRLTQLRQTDGEVAATFSNGQTATAEVLVGADGINSATLQLAWPNPRPKRWTEVTCFRGLIPRTVVASLRQADGSPLDNNPIHSFSMDRHRNDKSGATTYWVRGGELLNVWIARYEPDSAAFEKEEGDWFPVSREEIVREVGEAFTGHPSRNDLLALAGAIVRPTKWGLYDRDALETWVQGRICLLGDAAHPMLPTFGQGAAQSFEDAAALSCAFALHKSDVATALLHYERVRHYRATRFQLGSKFAFDHLRARDTAEQKALLEKLDERVSPAFAHDKRGGEDDSWIYAYDARTIGTELPPKRLGPWDFRRAAKASYAAITRTLWTPTGSAREPRPVTRAEVARHNTRDDCWVIIRGKVYDITEWAPHHPGGAGIARMYAGKEATAEFGDYHSAEAVAHMAHFCIGDLVEAAGA

>Bact_WP_011048843.1 monooxygenase [Ruegeria pomeroyi]

MRAIVIGAGMGGLMTALALRQSGVFSAIEVYEQTRQPSTAGAGLNIPPNGARICRWLGVDLDGGDPKGPDGAIDGGRAAILKSTRQFNTDGSVTERPFDHVTAVGDGAGFHHMHRLDLLMCLYKRVFEFGPDSGTPCPISVHMDSRLTGLEQSQGGVTAAFANGHTATGDILVGADGINSATLSLAWPSPRPRRWTEVTCFRGLIPRETVAALRKPDGSPLDYNPIHSFSMDRHKTDRSGATTYWVRGGELLNVWIARYEPDSTAFEQEEGDWFPISREEITRDVGAAFAELPNRDDLVALAGAMVRPTKWGLYDRDALDSWVQGRICLLGDAAHPMLPTFGQGAAQAFEDAAALGCAFALHRRDVATALLHYERVRHYRASRFQLGSKFAFDHLRPKDTEAQKALLEGLDERVTPAFSHDKRGGENDAWIYAFDARNIGPTLPAKKWGPWDYREASKEDHAKATQSLWRPEVPADGARRVTRAEVARHNTRDDCWIIVSGKVYDITAWAPHHPGGAGIARIYAGKEATAEFGDYHSAQAVAHMAHFCIGELVETSDDIQPPA

>Bact_OUW27504.1 monooxygenase [Rhodospirillaceae bacterium TMED167]

MRAIIVGAGMGGLFTALALRQSNAFSSIEVFEQTKEPATAGAGLNIPPNGARLCRWLGIDLDGGDPKGPHGAIDGGRAAILDSSRQFNEDGSTSERLFDHETSLGDGAGFHHMHRLDLLMCLHKRVAMFGPDSGKPCPITVHMDNRLTEFQQTAEDVTATFSSGLVATGDVLIGSDGINSATLELAWPDQRPRRWTGVICFRGLIARDDVAVLRKADGRPLDNNPIDSFSMDRHKTERSAVTSYWVRSGELLNVWIAHYEPDSKEFKQEEGDWFPVSQQEVADQVREAFAESPNRDDLIALAGAIDRPTKWGLYDRDALDTWVQGRVCLLGDAAHPMLPTFGQGAAQSFEDGAALSSAFERHGEDVGSALLHYERVRHYRATRFQLSSKFAFDHLRARDTLEQKALLEGLNERVSPAFAHDKRGGESDDWIYAYNAREIGTELPSKKWGPWDYRETNKEEFTAARQRLWKPASPADGTRSVTAEEVAQHNTKEDCWVIILGKVYNITEWAPHHPGGAGVARMYAGKDATAEFGDYHSKAAVAHMAHFCIGDLQPPARKQTR

>Bact_KND51202.1 hypothetical protein ABA06_02740 [Parcubacteria bacterium C7867001]

MKLFYPLLLTFVFAFVAGTAFGEYLAISQGKIVVEGASVQNGATALPTSAPSVSATPVATPTSPSVTPSAPAKTTTNKTSTPSGITKSTVAAHSTSASCWTIVSGNVYDLTSWINQHPGGSSAIRSMCGRDGTEDFLDQHGGDRRATAELASFKIGALTN

>Bact_KUG60468.1 hypothetical protein AVL63_08830 [Nesterenkonia jeotgali]

MTDSAPSAALPKTLLASLALPALLLLTSCGEGSQEQSPAEDQEAEQAETTELDEADSGEDPVEEDPVEEDQAEEEQPEGTDSEGAEQAISMAEVESNDSPDSCWAVLDETVYDLTTWIEEHPGGEARIEQLCGTDATEDFGAQHGGDSAPESQLAELEIGELEG

>Bact_OHA81475.1 hypothetical protein A2675_03315 [Candidatus Yonathbacteria bacterium RIFCSPHIGHO2_01_FULL_51_10]

MRSKAPIAGITIGIVTLLGMGAQAVYATTYTSAQVASHNTASDCWEIINGKVYNLTAFIPQHPGGQSAVIAQCGKDATTVFNNGPHSASTLSALSAYFLGDLNAASVVTPTTPPTPTPGTNMPSQVAPAPVANPTQTSQKFMNRDDDENDDEDDSDHSDISETRHEVDDEDSEYTLSTSRTSQEQRGRGDSSGDDD

>EukCyb5_XP_003857898.1 cytochrome bdomain protein, putative [Leishmania donovani]

MPAVYTRDQVAEHNSKESGWLIINNGVYDVSDFYDDHPGGRDILLAHIGTDATEGFEAVNHSRGAMRRLEKLKVGELPENERRRYITLEQVAAKKSAAGAWLVIHNKVYDVTPFLDLHPGGRDILLYSAGGDATQAFTGNGHSDTAYQMMGKYVVGDLEPSERKTLVNRKATGAKQAATTQMLHVKNENASLLLHIQEQLRLLMALALFVIAGVFLLS

>EukCyb5_XP_003721610.1 putative cytochrome bdomain protein [Leishmania major strain Friedlin]

MPTFYTRDQVAEHNKKKSGWLIINNGVYDVSDFYDDHPGGRDILLAHIGTDATEGFEAVNHSKGAVRKLDKLKVGELPENERRRYISMEQVAAKKSADSAWFVINNKVYDVTPFLDLHPGGRDILLYNAGGDATQAFTDNGHSDTAYEMMGKYVVGDVEPSERKTLVNRKATRTTQAATTEMVCVKSENASLLAHIQEQLKLLMALALFVIAGVFLLS

>EukCyb5_XP_003871608.1 putative cytochrome bdomain protein [Leishmania mexicana MHOM/GT/2001/U1103]

MPTVYTKDQVAEHSHKESGWLIIQNGVYDVIDFYDDHPGGRDILLAHIGTDATEAFEAVNHSRGAMRKLEKLKVGELPENERHRYISMEQAAAKKSADGAWLVINNRVYDVTPFLDLHPGGRDILLYNAGGDATQAFTDNGHSDAAYHMMGKYVIGDLGMSERKTFVNRKSTGATQMAHVRNKYASLLAHIQAQLRLFLALALLVIAGVFLLS

>EukCyb5_XP_001561521.1 putative cytochrome bdomain protein [Leishmania braziliensis MHOM/BR/75/M2904]

MPTLYTKDEVAAHNVKENGWLIINNSVYDVSKFYDDHPGGRDPLLAHIGTDATEAFEAVNHSRGAKYKLEELKVGELSENERRHYISLEQVAAKKSANGAWFVINNKVYDVTKFLDLHPGGRDILLCNAGGDATQAFTDNGHSPAAYKMMSTYAIGDLEPSERKVFVTQKATGERSGATTAMVGVKSGNESLLIQIQQQLKFLIIVALFIIAGVFFLT

>EukCyb5_KPI85339.1 Cytochrome b5like protein [Leptomonas seymouri]

MTAIYTKAQVAEHNKKESGWFIINNSVYDVSKFYDDHPGGRDVLLANIGRDATEEFEAVHHSKGAMRKLEDLKVGELPPSERRRFISMKEVSAKKSAEGAWFVINNKVYDVTKFLDLHPGGRDILLYNAGGDATEAFTDNGHSDNAYRMMARYCIGDLELDERKKFVNRKVTGEQHNSTVKAVRVKSEDESLLGRIQEQLQLFIVLALFVMAGVFLLS

>EukCyb5_XP_015653265.1 putative mitochondrial Cytochrome b5like protein [Leptomonas pyrrhocoris]

MTATYTKAEVAQHNKKENGWFIINNSIYDVTKFYDDHPGGRDVLLANIGRDATEEFEAVHHSKGAMRKLDDLKVGELPESERRHFISMSEVATKKSADGAWFVINNKVYDVTKFLDLHPGGRDILLYNAGGDATEAFTDNGHSDNAYKMMGKYCIGDLELDERKKYMNRKVTGEQHASTTQVVRVKSEDESLLARIQEQLQLFIVLALFVMAGVFLLS

>EukCyb5_EPY29873.1 cytochrome bdomain protein [Angomonas deanei]

MSTTYSRADVAKHNKKEDGWFIVNNDVYNVTKFYDDHPGGRDVLLAKIGTDATEDFEAINHSNGALKKMKELQIGTLPEAERRKYISMKEVETKMTADAAWFVINNKVYDVTKFLDNHPGGRDIILCNAGQDATESFINNNHSEGAYKMLAGLVIGDLEPSERRRVMRRKGEAGHGQQEVAPTSSHGKRGDESLLERLKEQLTLFIGFVVFIMAGIILLR

>EukCyb5_XP_005537823.1 similar to cytochrome B5 [Cyanidioschyzon merolae strain 10D]

MAMGRTTQFTLDEVAKHADKDSCWLVIDGKVYAVEKFLNEHPGGEDVLLETAGRDATREFEDVGHSKSAREQLKEFYIGDVREPTAEELAAKRAVQASAAAERRDGDGVRANGLGKERLFAGGAFGAEPLSGGSSTWTLLKRLFVPVCIAVLVYLVREYSKQVQTGG

>EukCyb5__XP_004997136.1 cytochrome b5 type B [Salpingoeca rosetta]

MVWSIKFPGKPTICVTAQKAQLVRAKHQQAKQAKEEEGATSSNHKTPRVDTSQHIQAAKHTNQHQPPPSLPTTTPATMSKTISYAEVAKHNTNESCYMVIHDKVYDVTKFLIEHPGGEEVMMDYAGKDASEGFEDVGHSEDAREQLSDFVIGELPADEKGQAATESNIAPQAPSRAQASSDGSWFLYVALAAVVAGAGVAYFYFNK

>EukCyb5__sp|P04166|CYB5B_RAT Cytochrome b5 type B (Mitochondrial) OS=Rattus norvegicus GN=Cyb5b PE=1 SV=2

MATPEASGSGRNGQGSDPAVTYYRLEEVAKRNTAEETWMVIHGRVYDITRFLSEHPGGEEVLLEQAGADATESFEDVGHSPDAREMLKQYYIGDVHPNDLKPKDGDKDPSKNNSCQSSWAYWIVPIVGAILIGFLYRHFWADSKSS

>pdb_3MUS_EukCyb5_A Chain A, Cytochrome B5 Type B

DPAVTYYRLEEVAKRNTAEETWMVIHGRVYDITRFLSEHPGGEEVLLEQAGADATESFEDVGHSPDAREMLKQYYIGDVHPNDLKPA

>EukCyb5_XP_002904260.1 cytochrome b5 [Phytophthora infestans T304]

MAETAAPETPAPAAAVKEFTMEDVAPHNTTEDCWMVIREDGVRKVYDVTAFLDDHPGGPEIMVDVAGQDATDEFEDIGHSNDARAQLKQFEIGKIKGDAKKEATATTSAGGSKVSASGRQDISGDNGPLYAVLAAAVVFTALYFKYQ

>EukCyb5_XP_008909162.1 hypothetical protein PPTG_14330 [Phytophthora parasitica INRA310]

MAETAAPETPAPAAAVKEFTLEDVASHNTAEDCWMVIRDDGIRKVYDVTAFLDDHPGGPEIMVDVAGQDATDEFEDIGHSNDARNQLKQFEIGKIKGDVKKEAAAKTSAGGSKVSASGRQDISGDNGPLYAVLAAVVAFAALYFKYQ

>EukCyb5_POM78817.1 Cytochrome b5 [Phytophthora palmivora var. palmivora]

MAETAAPETPAAVKEFTLEEVAPHNTAEDCWMVIRDEGIRKVYDVTAFLDDHPGGPEIMVDVAGQDATDEFEDIGHSNDARAQLKQYEIGKIKGDVKKEASAKTSASIGGKASASGRQDISGQNGPLYAVLAAVVAFAALYFKYQ

>EukCyb5_XP_642935.1 cytochrome b5 B [Dictyostelium discoideum AX4]

MSEDKQYTMEEVSKHDKVDDLWMVINQKVYDVTSFVNDHPGGGDYLIQNAGKEATNEFLDVGHSQKAVDMLKDYYIGVCTDSKPLQNPLSSNPIPISTESPVIKSTESSSNNNNNEDKNKDVTTTIALVGGIVVAVASIAFFTFAKIKK

>EukCyb5_XP_020437449.1 hypothetical protein PPL_02343 [Heterostelium album PN500]

MDVQQSYTLEEVAQHNKRDDLWMVIDGKVYNCTEFVDEHPGGGDYIIDNAGRDATLEFIDAGHSEKAIALLKDFYIGECSNAKHIGKLNAPATAATSNATTTEKTESESVSSSQTTDNSTNNISTRKSSSSTSEPTNSYLIPLIIIGAVVVVSAVIANNNINSIDIYSGSFHRGLWTVEDLKQTGWDFNSMNADEKQKFNLLVLQRILDQMAQPSQLRKPVSKIPYYGPYTSFVNDPYKTDPYTGAYQINPVLGGDNFISGLFGTSGGPLGGRFGASGFNGGGLIAGGYPGQFVLSAIFPNQYMANGPIAGLLTGDIRNPIYDATRHPPPYISTAYGPLAPYLQKRQGGDGWQN

>EukCyb5_EWM28245.1 cytochrome b5 [Nannochloropsis gaditana]

MAPQQFTLEEVAGHNSEKDIWLIIGNEKTGGPKVYDVTKYLEEHPGGSEVLLDHAGKYADEMFEDIGHSGDAKEKLKTLMVGELTPEDVEKLASEKVSRASELPSKGGLNPFAVVVLLVAIVLGLYLSKQRTEELGSYNRRDLQMLLWAERGGDQNRACFHGMCQERDEHCVAKATGTRPWRNMTGTERVQRRGAGAVDMGDYGRASRSRWLTQIL

>EukCyb5_GBG32565.1 Cytochrome b5 [Aurantiochytrium sp. FCC1311]

MSNTEDSKAAATSADPAAEPAKSAPADDGVKVFTRGEVAGHTNEENDLMIIIMGKVYKVDKYLEEHPGGPEIIADCAGQDATEEFLDTGHSAEAQETLKKYFVGNVEGGDTSSSSATTKSGSEGPSMTMMIAVLAILFAIFAAVKLQA

>EukCyb5_XP_022587929.1 putative cytochrome b5 [Cyclospora cayetanensis]

MEDLMKLPELSWEEIRKHTSAESCWCVFHGLVYDLTKFLNKHPGGSHIIIETAGRDATDAFEDIGHSLEARIMADEYIIGRVEGATNIRRCKPEAKAERCVVSAAKGSPSTGVFIAIFLVALIGIWAYFQFQDEDSTFHHRISAAPQHPIE

>EukCyb5_PFH31116.1 cytochrome b5 family heme/steroid binding domaincontaining protein [Besnoitia besnoiti]

MATRETETGREPWRTRVVSAEEVRKHNSEKDFWCIIHGVVYDLTPVLDKHPGGVEVLMDYAGEDASDAFEDIGHSFSARRMTVGLEVGVLAGAEDNAPGALSKTAKAATSAAREKADCSTCAGGLLSGRTAAGALIVLAAAATVFYILGIS

>EukCyb5_XP_012194396.1 hypothetical protein SPRG_00789 [Saprolegnia parasitica CBS 223.65]

MRHGRGLKTAFVDTLWKRRQRVLLHMSDLHEFTAAEVLRHASTDDCWLILGDDGRQKVYDITAFLETHPGGPEILMDLAGQDAHEEFKEVGHSKAAQDMVEQLCIGRLRIDGRRKPKRSRVVLPVAVNDETSPRNDRLVALMGVLMALLFGYLLVPNSVL

>EukCyb5_XP_008605243.1 hypothetical protein SDRG_01494 [Saprolegnia diclina VS20]

MRHGRGFKTTFVDHLRKRRQLANMSELHEYTAAEVLRHASTDDCWLILGDDGRQKIYDITAFLESHPGGPEILMDLAGQDAHEEFKEVGHSKAAQDMVEQLCIGRLRVDGRRKAKRGRVVLPVAVDDEKRPRHDRLVALMILLMALCFGYLLVPNTAL

>EukCyb5_XP_001459012.1 hypothetical protein [Paramecium tetraurelia strain d42]

MNGEKRIVGWDELAEHSNRTSLWVVIEGQVFDVTTYLAEHPGGDDILIKYGGLDGTQKFLEVNHSNYARSLRNARLVGTLTSDPQPNDYLKAVKSKKQKNNFNPNRQITWEELGQHNKKEDLWIVIEGKVYDVTDFQDDHPGGPAILLGKAGDDATAAFHDANHSQSAYKQLEKLQVGVITGVKPNLSGSGSSTNLIFVILLILAIGAGIFVITK

>BactPrG_AHK78592.1 cytochrome b5 [Halorhodospira halochloris str. A]

MADEADEALPVITQEELARHDQPEDCWKAIHGKVYDITDYLPRHPGPPALVLHWCGQESTEAWETKGYGMPHSDAAGILLEDYLIGILEEEEASADRQQD

>Bact_WP_044414127.1 cytochrome b5 domaincontaining protein [Rhodopseudomonas palustris]

MMRKLYLATTSLFWVMVLAFWAGDVLSPHAEQPAAISASRDITSAELAKHATPQDCWMAIRGQVYDLSAYLPDHPSRPQIIEPWCGKEATQAYDTKTKGRPHSKEADDLLPKYRVGRFVPGAD

>Bact_WP_104522120.1 cytochrome b5 domaincontaining protein [Rhodopila globiformis]

MMRTLYVASTTAFWALVIAFWAAGTWAPHAEQPPAAPADRAITAAELARHARPEDCWMAIRGSVYDLTPYLPDHPSRPQVIEPWCGKEATDAYNTKTKGRPHSKEADSLLPQYRIGRFVPDAK

>Bact_WP_011472391.1 cytochrome b5 domaincontaining protein [Rhodopseudomonas palustris]

MMRKLFIATTATFWAMVLAFWGGSLWSPDAEQPLAAAPDRAIAAAELAKHTTPETCWMAIRGDVYDLAAYLPDHPSRPQIIEPWCGKDATEAYNTKTKNRPHSKEADEMLPKYRIGRLLPER

>Bact_WP_011156558.1 cytochrome b5 domaincontaining protein [Rhodopseudomonas palustris]

MMRKLFYVSTAAFWIAVAGFWIANLLVPGTDTTATAAEREIGAAELAKHAVPQDCWMAIRGNVYDITAYLPDHPSRPSIIEPWCGKEATEAYDTKTKGRKHSSEADALLPKYRIGRFIGGS

>Bact_WP_012494763.1 cytochrome b5 domaincontaining protein [Rhodopseudomonas palustris]

MMRKLFYVSTAAFWIAVAGFWIANLLVPGADTMATAAEREIGAAELAKHAVPQDCWMAIRGNVYDITAYLPDHPSRPSIIEPWCGKEATEAYDTKTKGRKHSSEADALLPKYRIGRFIGGS

>Bact_WP_107343201.1 cytochrome b5 domaincontaining protein [Rhodopseudomonas palustris]

MMRKLFYVSTAAFWIAVAGFWIGNLLVPGTDTSATAAEREIGSAELARHAVPQDCWMAIRGNVYDITAYLPDHPSRPSIIEPWCGKEASEAYDTKTKGRKHSPEADALLPKYRIGRFVGGG

>Bact_WP_107354441.1 cytochrome b5 domaincontaining protein [Rhodopseudomonas palustris]

MMRKLFYVSTAAFWIAVAGFWIGNLLVPGTDTSATAAEREIGGAELARHAVPQDCWMAIRGNVYDITAYLPDHPSRPSIIEPWCGKEASDAYDTKTKGRKHSPEADALLPKYRIGRFVGGG

>Bact_WP_022721647.1 cytochrome b5 domaincontaining protein [Rhodopseudomonas sp. B29]

MMRKVFFSSTVLFWIAVLGLWIGNVITPGTSGPAAAADREIDAAELARHASPEDCWMAIHGAVFDITAYLPDHPSRPSIIEPWCGKEASSAYDTKTKGRSHSRDADALLPKYRIGVFKP

>Bact_WP_041798956.1 cytochrome b5 domaincontaining protein [Rhodopseudomonas palustris]

MMRRLYFAATATFWIAVLGFWAGSALTPVTRQPAVAADRDITAAELARHATPADCWMAIRGSVYDLTGYLPDHPSRPSIIEPWCGKEATDAYNTKSKGRAHSGEADAMLPKYRIGRFVTGSAQ

>Bact_WP_092680997.1 cytochrome b5 domaincontaining protein [Rhodopseudomonas pseudopalustris]

MMRKLFLASTATFWVIVIGFWAGSALTPVAQQPAVAAEREITAAELASHAKPDDCWMAIRGGVYDLTSYLPDHPSRPTIIEPWCGREASDAYNTKTKGRAHTGEADAMLPKYRIGRFLPGG

>Bact_WP_027277557.1 cytochrome b5 domaincontaining protein [Rhodopseudomonas palustris]

MMQRLFLTSTAVFWLAVLGFWIGSLSAPIAQPPAAPSVADRSISTAELAQHATPQSCWMAINGAVYDITGYLPDHPSRPQIIEVWCGKEASDAYATKTRGRPHSHEADQLLPKYRIGRFAP

>Bact_WP_103012903.1 cytochrome b5 domaincontaining protein [Rhodopseudomonas palustris]

MMQRLFLTSTAVFWLAALGFWIGSLSAPIAQPPAAPSMPDRAISAAELAQHATPQSCWMAINGAVYDITGYLPDHPSRPQIIEAWCGKEASDAYATKTRGRPHSREADQLLPKYRIGRFAP

>Bact_KKR12926.1 hypothetical protein UT41_C0001G0470 [Candidatus Wolfebacteria bacterium GW2011_GWC2_39_22]

MKKLTLISLFIFWAFVTSLFTAGFALQEKTNNPTPQNNPAQLIAQGAILTKEELAKHASASSCWLLIDKKIYDVTSYLNQHPGDADTILPTCGTDATRAYSTKGRTTSPSPHSQNAHELLKAYYIGDLGQEAIAVTPTASPTTKPQSNTPATQPTAPSTQPNTSVALSIQEIAKHNTAQDCWMIVNNNVYNVTSYIPRHPGGASKILTYCGKDGGAVFEGLPHSTNAHQLLASFFVGAVGQVVNTQTVQQTTSPTVPVGTSNNRGDDDEDDD

>Bact_KKU37079.1 hypothetical protein UX49_C0002G0004 [Candidatus Wolfebacteria bacterium GW2011_GWC2_46_275]

MKKLTAVSLLIFWALVTALLTSGLVFYQNKQQAVEIPPEQQSASVVAPGALLSMREILTHNTAASCWLLVSGKVYDVTTYLDKHPGDASTILPTCGTDATQAYATKGRTSSPKSHSQNATEMLKAYYIGDFGQKPVAQNAATQKPTTATAQPTTSTAKPTTLSTTVTNPSVTLSIQEIAKHSTTDDCWMIVNGNVYNVTSYIPRHPGGVGAIRPYCGRDGTAAFEGLPHSMNAHQMLANY

>Bact_KKQ25926.1 hypothetical protein US39_C0003G0026 [Microgenomates group bacterium GW2011_GWC1_37_12b]

MKIKVVVGIALVLFTIIIGSIGVAGLVLYDQKKNVGPAQKLASTDLSISDQNPSVQSNSELGGISAEVVSSHSSSSDCWMIIDGNVYDFTKFLNIHPGTSATMLPYCGRDGSNAFATKDKNPGTAHTSIARDLLGQYYLGTLGNGSGSNLAQNPTIPGTNSKLTPTTRPGTTIPTGSTTNVPASSSSNILTTATVSAHSSSSDCWIIISGNVYAVSGYLNSHPGGAGVIANYCGRDASNAFATKDRNPGSSHSSFAVSQLSGLLVGRIGTPVTGSGNTGTNTAATPTPTQRINTPTPTTIPGGGGGGGGGSVTLTTTQVSAHNTLQDCWMIISNRVYNLTGYIAQHPGGQNAILNYCGRDGTNAFDTRGGSGTHSNNARNLLNGFYIGDLGTSVPVGSTPTSTPLPGSATSTPIQNTSQVPSVVLQKYPDATLRGEIEYEDDGRMELKITTSGQCRDIKINSSGVISEDKSC

>Bact_KKR28269.1 hypothetical protein UT61_C0053G0005 [Candidatus Woesebacteria bacterium GW2011_GWA1_39_8]

MAKNKRNVRTEEAIAKHNKAEDCWLIVDGNVYDVTVFVAQHPGGSDLIVGYCGGDATAAFNTRDKNPPEKHSSFAASLLRSYYIGPVGRLPENILYPDPNDPTKVVIKPFGSVGPIPTLTASQAGGGSSALIDAAEPYCGKDGTQAYQSKGGQGGNHSAYAYSLLGNYFIANLGSSVALNNGSPLVPTVAASANPTATPVVSGGGGASASNLTLSVSEIATHNILQNCWLIISGSVYNVTSFISQHPGGVTQITNWCGKDGTSAFQTKGGKGSNHSSYAYSLLNPYLIGTVGSSVSVNATPTTASSGGGSTTQTSAGIPSVIIDKYPEATLIKGEYEDDGRWEGKVNTNSGCRSIKINSSGNITNDSSC

>Bact_KKW01684.1 hypothetical protein UY34_C0009G0009 [Parcubacteria group bacterium GW2011_GWA2_48_9]

MKTETIIGIIGTIILVVFIGITWNTYTITPSAQLNEVTLSNQQVQNQLQEAVNSGSFTLDTALISQHSNADSCWLLISGKVYDVTQYLQLHPGGRAIILPFCGKEATAAFDTKAGGGSHSNNAISQLGAFLIGDLNMLLTELPVNSPALSNTAGTVNTNQEPSPAPVTPVPTSITLDAAAVARHDNSRDCWLIINNSVYAVTSYLARHPGGQAIIVPFCGKDATAAFATQAGSGSHSNSAVNQLAAFKIGVLGSTTTVQDVQQIQQNTNNLPAQGNGEDESEEENEEEEEDD

>Bact_OGF28161.1 hypothetical protein A2227_06710 [Candidatus Falkowbacteria bacterium RIFOXYA2_FULL_47_19]

MKKTVLFVLIFFGISVSLMIAGGSFRPTSGVKIDLGGNKTSGITVADNVPEKPAILAAVEVAKHNSISDCWMIINGKIYDLTAFFSQHPGGAEAISRHCGTDGTEAYGTKDKNETDGHSSYAKSLLADYYLGDLNGALPEEKIRMAENKDAEGNTGIKSAIENEKTPIGDSVAGAGGITSAALAGHNSPSDCWMALSGKVYDVTAYLRRHPGGDAMLAYCGRDGTTAFNGRGHSDYARSLLPAYYIGDFGDSAGPAPEISGGQTRSGGLSAVEKEYPGAIILEENIEDDGRREIKLLYEGRQYEVRVGPGGDILRTEDH

>Bact_OGG92595.1 hypothetical protein A3G63_00260, partial [Candidatus Kaiserbacteria bacterium RIFCSPLOWO2_12_FULL_52_8]

MVTAIFAAGLLSANSAGGGIAVGQNSGVAGQVDGVVPAQSLDASGTALVLNAVELAKHNSSSSCWLLISGKMYDVTTFLNQHPGNAGTMLPYCGKDATTAYANKGTAGGSPHSSTASAMLAGYYIGTLNQSIAVSSGKTTASTETNSTVQVAPSPAPVIQTPVVPAPVAPSPTAPRPAPSSTV

>Bact_OHA60684.1 hypothetical protein A2556_02565 [Candidatus Vogelbacteria bacterium RIFOXYD2_FULL_44_9]

MKNYVSISLFIFWAVVVAVMTAGLVFSDQNKTPTQITGDQIATSPSNSSANSNQKTTLTMTEVAKHNRATDCWQVINNNVYNFTSFLSQHPAGADAMIPYCGQDSTIAYNTQGGRGQDHSAQARSLLASYLIGPLGQTVTAREITPIPAPTNNNQTVRENKREQEDD

>Bact_OGL88469.1 hypothetical protein A3I42_03275 [Candidatus Uhrbacteria bacterium RIFCSPLOWO2_02_FULL_49_11]

MKKELLIGIVGSLLVLAGAGYYTMHYRAQTAQLDALMKRAAPSTNIQNSSPPVSVITLTKEEIAKHNSASDCWIIVQGSAYAVTGFLNIHPGGAAAIAPYCGADATQAFLTQGGQGTHSAVADQQLATLLIGKVGAQVAPTAIQKADINAATIPFSGEREDENERDDD

>Bact_OGZ32431.1 hypothetical protein A2V69_02320 [Candidatus Portnoybacteria bacterium RBG_13_40_8]

MIYLYKILRITAWLLVIATVLSLFSGYLSIKYFSSKGIDYRNLHVSIVPWIFIPLFYIHSAIGFLNLLTSHKSTNKKSVKIFVEIVWATVFILLILVIVAKPSAAITPNNNNSSGSGNFSLTIEEIAKHNSTSDCWLIINNRVYDLTSYLRAHPSGVYTISPYCGKDGSRGFATKDIGTSHSATANNLLNSFYIGDVAKD

>Bact_PCJ24955.1 hypothetical protein COA94_06930 [Rickettsiales bacterium]

MSSKLTIFTLKDIRDMPESRVVIILNDQVYDVTDYLVCHPGGREVLLENNRKDATEAFSAIGHSDRAEEILKEYKIGSLAEKERSGGPPLQSNALSSSPEPLKRVMHIKNKLVTEEDPYFFHKIFGLIVLLHMAIRFIVLGLDAGEVIIQRDIIGFNSDNVFSDGTKLFFAFCHGILSVSSFIFAVPKHSSQNKPMIHQMFRGHSVCFASRAVLCMVVDVLVQSPELKRFLISCIVLSCLIAADQITKYLATEDDRYKTTNSMPYWRGCSVEREHLHKTFYAFAQFLASIICLFGSYTTIFFTLPAIQGAALLMTLTRKNIITSHSYHQIYTFLLFYPIPLFLVLWPAKTIFAIILSGVLYYLRSRNINKYLLWIPVLIMANMIELTPENYPNGLIIGAAIFLLTLYIVKSNAKIIETERIDGNNRVVSHKQITPDAYELIIRTQTPIKLEIGQHVLIQINDSLNRKYTPIWTKYLPETDQTLLCLRIKEYKSTERLTASSYLARCGEGSVLALHGPYGNKFYCPKNDAIIDKINKCQYNLEDYQVYLFSAGSGITPIYQLAKNICKPGQKRGDSKNGGEKGDGKEKGREKGKKNEKLTLITCDQKHENQMMKDELTQLKEDFPNELDWLCFLSREGKNSPDIPATIFKERLTPGKLMNIVKCDSPSLIIICGPDKWQEMIQQTINLIDFDLDKPPPKCKVLAW
