## Supplementary figures and images for "MAPR origins reveal a new class of prokaryotic cytochrome b_5_ proteins and possible role in eukaryogenesis"

### Fig1A_168sx61gf-tree-1000xPoiss-full.pdf

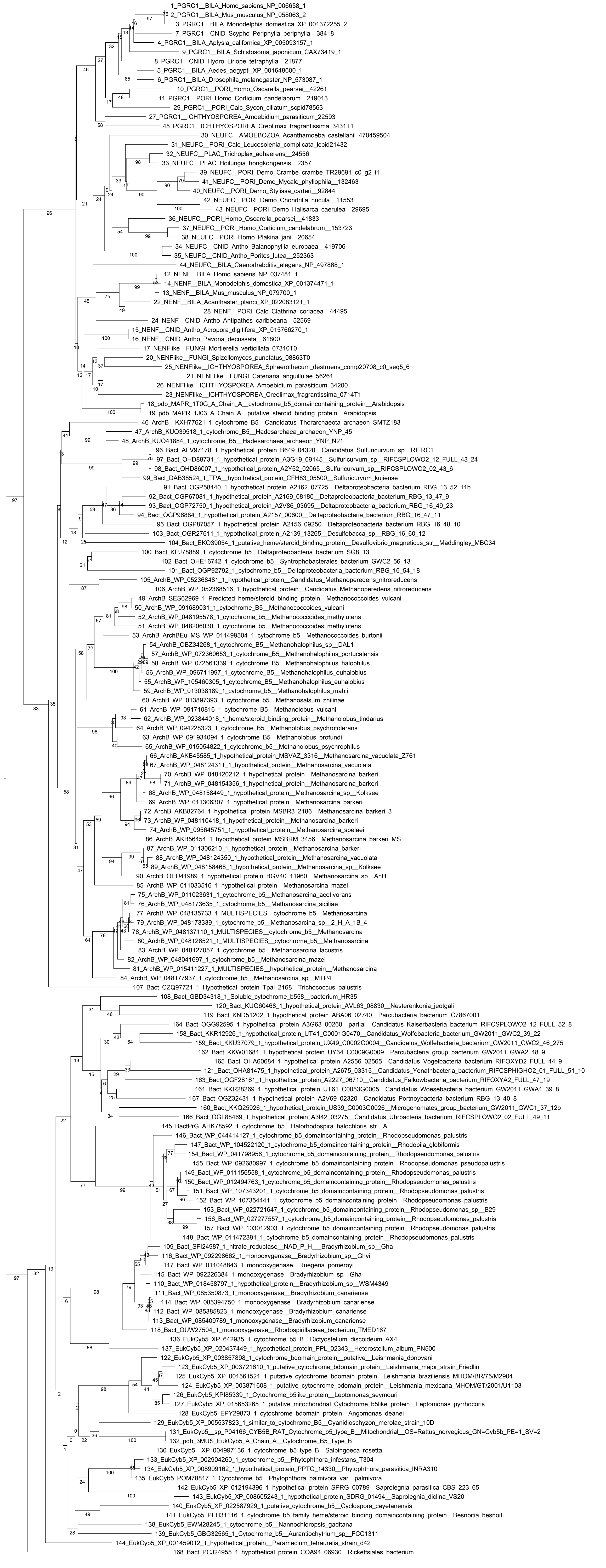

### Fig1B-tree_1000xPoiss.pdf

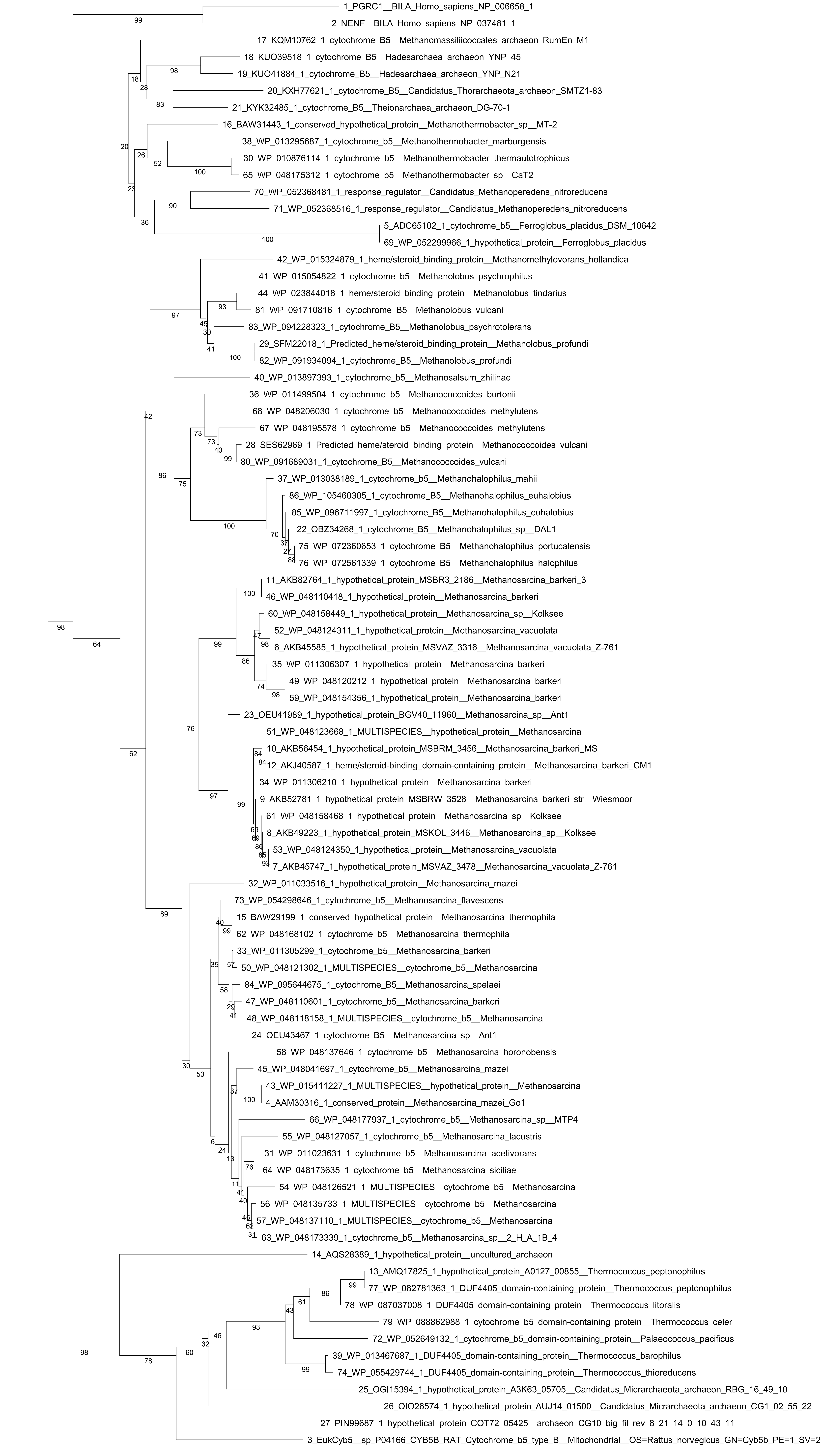

### Fig1c_tree_1000xPoisson.pdf

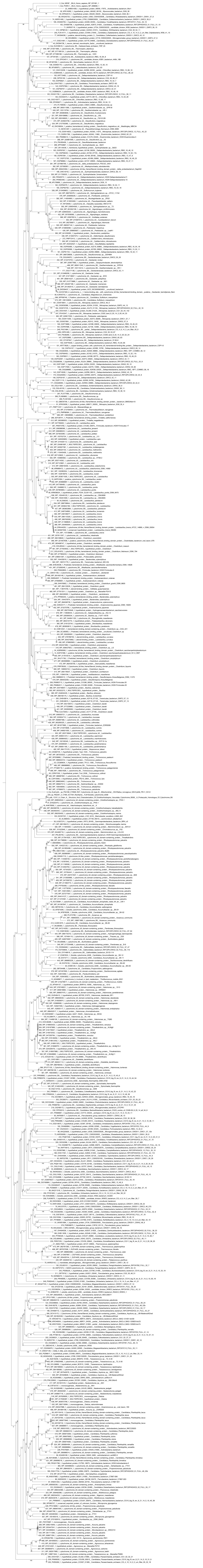

### Fig1d_tree_1000xPoiss-full.pdf

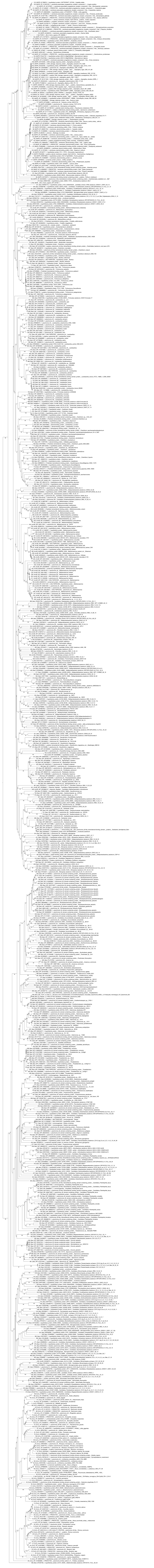

### Panel-A_CPRcytb5-affin_Clade1&2_Poiss.pdf

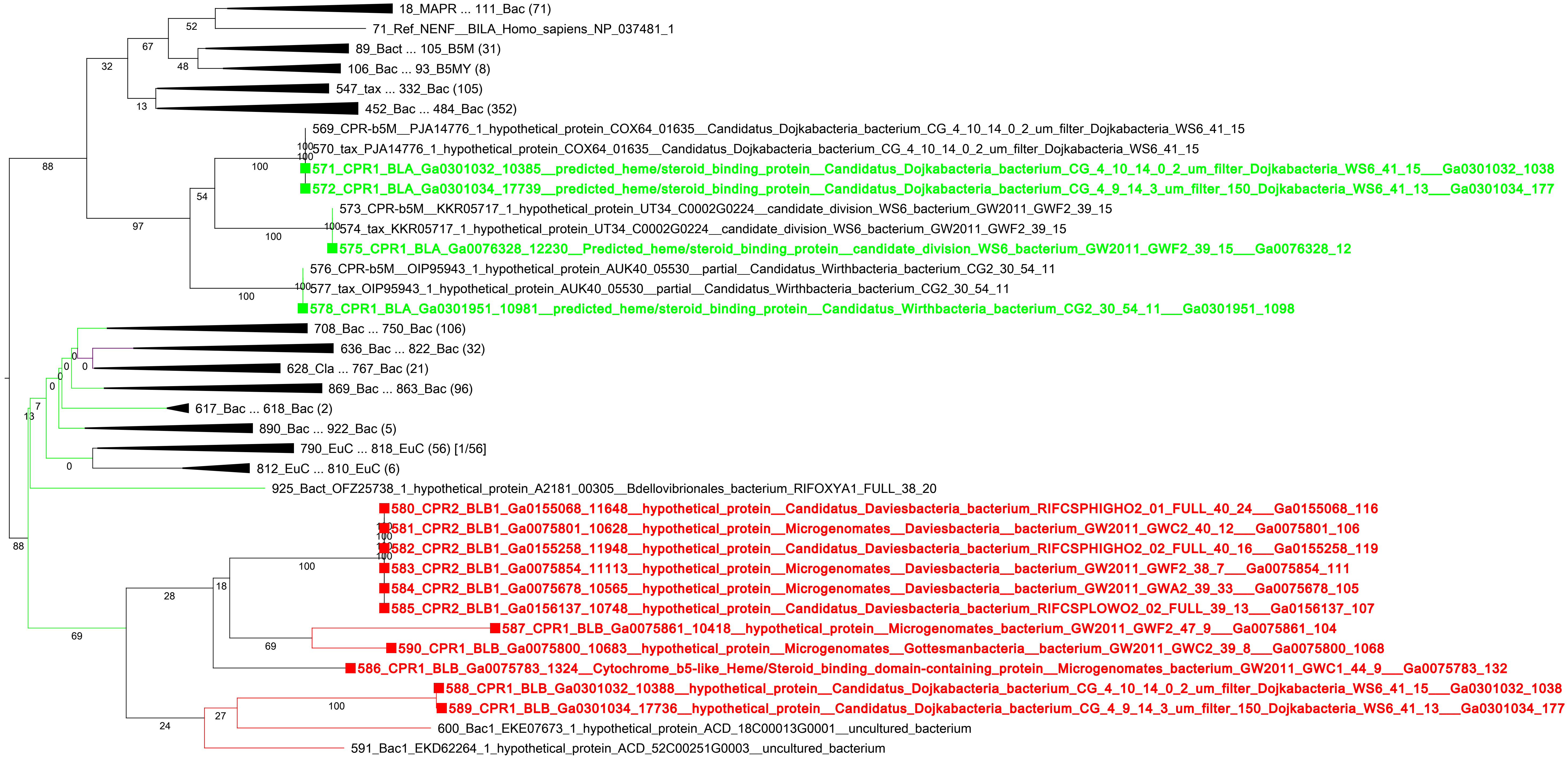

### Panel-A_CPRcytb5-affin_Clade1&2_Poiss_full.pdf

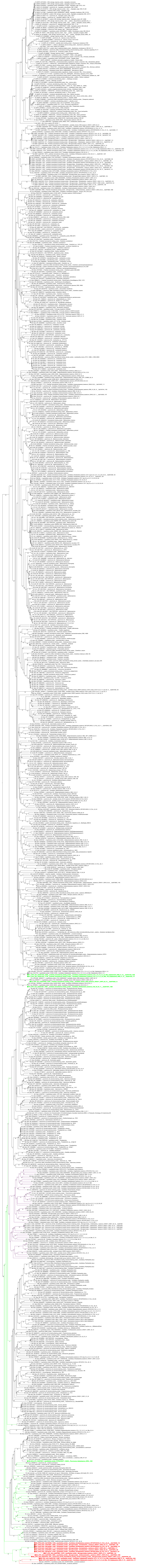

### Panel-B_CPRcytb5-affin_Cl1&2_BLB1_Poiss.pdf

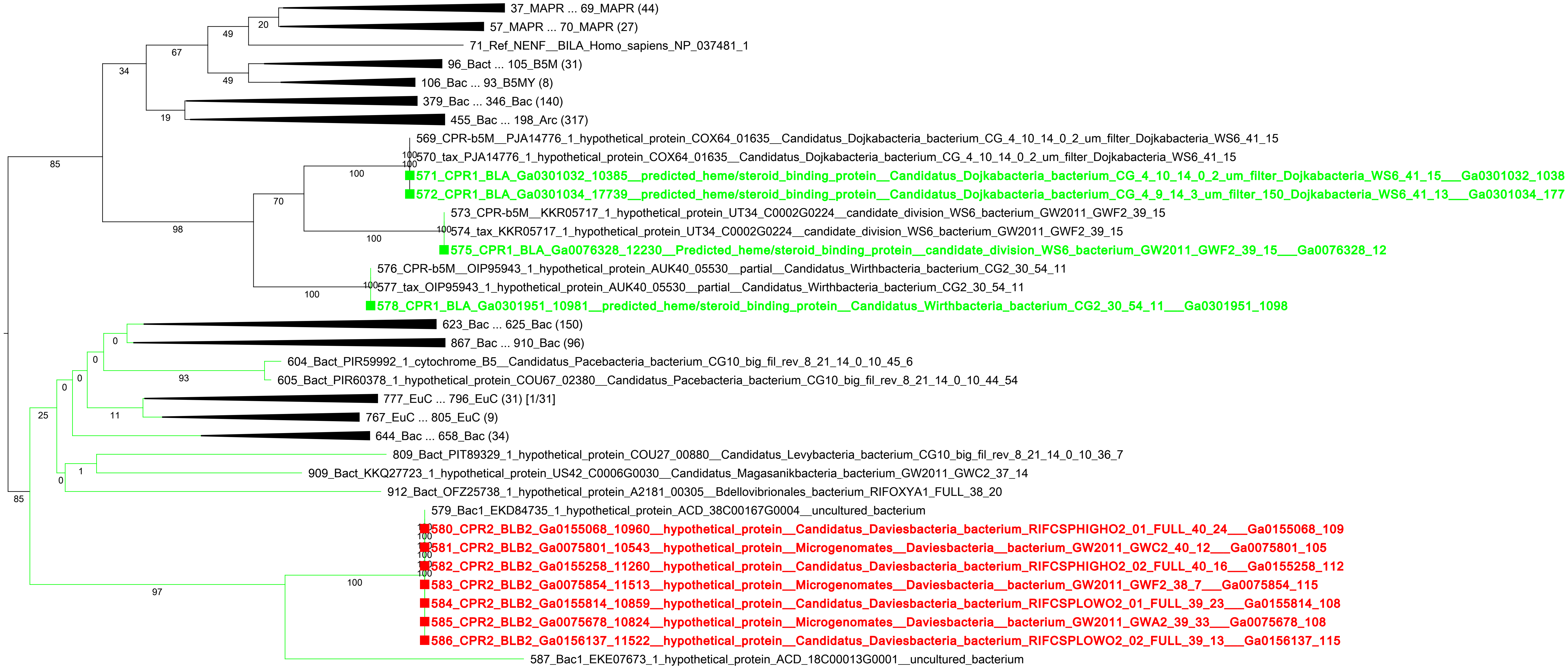

### Panel-B_CPRcytb5-affin_Cl1&2_BLB1_Poiss_full.pdf

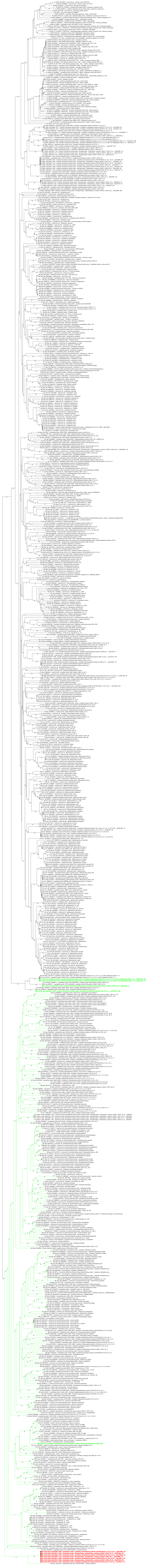
