## Supplementary Appendix B for "MAPR origins reveal a new class of prokaryotic cytochrome b_5_ proteins and possible role in eukaryogenesis": README_cytb5M_BLAST_Euk.txt

Method

The top 100 BLAST hits after using each respective cytb5MY sequence as BLAST query agianst the eukaryotic sequence data base (NCBI BLASTp) were added the the following 16 control sequences, where >PGRC1 and >NENF are MAPR proteins, >b5MY correspond to cytb5MY proteins, >C1 are clade-1 control proteins, and >C2 are clade 2 control proteins. These were subjected to L-INS-i MAFFT, and the MAFFT-aligned sequences are provided in fast files saved by the AliView program. The procedures are described in detail in the Methods. All eukaryotic proteins contained MIHIR residues at residues aligned with the MIHIR of Fig.1e.

Reference proteins:

>PGRC1__BILA_Homo_sapiens@NP_006658_1
MAAEDVVATGADPSDLESGGLLHEIFTSPLNLLLLGLCIFLLYKIVRGDQPAASGDSDDDEPPPLPRLKRRDFTPAELRRFDGVQDPRILMAINGKVFDVTKGRKFYGPEGPYGVFAGRDASRGLATFCLDKEALKDEYDDLSDLTAAQQETLSDWESQFTFKYHHVGKLLKEGEEPTVYSDEEEPKDESARKND
>NENF__BILA_Homo_sapiens@NP_037481_1
MVGPAPRRRLRPLAALALVLALAPGLPTARAGQTPRPAERGPPVRLFTEEELARYGGEEEDQPIYLAVKGVVFDVTSGKEFYGRGAPYNALTGKDSTRGVAKMSLDPADLTHDTTGLTAKELEALDEVFTKVYKAKYPIVGYTARRILNEDGSPNLDFKPEDQPHFDIKDEF
>b5MY_KKR05718.1 hypothetical protein UT34_C0002G0225 [candidate division WS6 bacterium GW2011_GWF2_39_15]
MKKKIVILLILALVVFVVMLGGTYVTRMIQVGKTRQVAGIKKEGLSVYKRSDLSKYNGAIKGRPILIGLN
GYVYDVTEGKKFYEPGGTYHYLAGKDSSSELNLIGGDIIRRKYPIVGVIVD
>b5MY_OHA80798.1 hypothetical protein A2675_01620 [Candidatus Yonathbacteria bacterium RIFCSPHIGHO2_01_FULL_51_10]
MSRSFIKSLGSPVLLFAAGVILLGLTYVLVQAGRVPAPVEVPSGLPVFSNADLTRYNGDDATLPIYVGLD
GYVYDVTPGKEFYAPGGAYHSIAGTDASSELRFFGGTIIREKYKIVGTIAP
>b5MY_OHA83705.1 hypothetical protein A2408_00355 [Candidatus Yonathbacteria bacterium RIFOXYC1_FULL_52_10]
MNSFRHHPRTRLIFSGIAILALMVGIFVILQLNANPVVPQGQAEDLPFFTKTSLAQYNGEIPDLPIYLAL
DGYVYDVTPGKEFYVPGGTYHHIAGKDASQELKVFGGNLIREKYSIVGKFTPE
>b5MY_KKS94036.1 putative steroid-binding protein 3 [Candidatus Collierbacteria bacterium GW2011_GWC2_43_12]
MNKRIKLYTALIKKLSLITSLVCLLALGLHFSNSKSPSPSYLPDFTQKYLATFKGVDSKKPIYIALNGLV
YDVSEGREFYIAGGPYHDLAGKDSSKELNLFGGEIIKRKYPIVGRLVN
>b5MY_KKU89055.1 hypothetical protein UY18_C0004G0020 [Microgenomates group bacterium GW2011_GWF2_47_9]
MKVKRARTGLVITAATILLAALLTRYYWPARINESIDQPLPEYDSEALMRFDGTDPNKPIYLALDGLVYD
VSMGREMYYGPGESYHYLVGRDASQPLHVFGAEIIKKKYQVVGRYR
>b5MY_OIP98122.1 hypothetical protein AUK40_01915 [Candidatus Wirthbacteria bacterium CG2_30_54_11]
MKKKILFALIMIVLIASSAFTVYHVFTERQARAKISALESSPSISFFTAADLARYDGTDPILPIYLALDG
YVYDVTAGSSFYAVGGDYHYLAGRDSTTDLQIFGGDIIKRKYPIIGKYVN
>b5MY_PJA14771.1 hypothetical protein COX64_01640 [Candidatus Dojkabacteria bacterium CG_4_10_14_0_2_um_filter_Dojkabacteria_WS6_41_15]
MKGRKIPILIGVAIGLLILGTFGYAKWHSAKSASNTPTTAETASLKEFTVAVLAQYDGTDSSKPIYLAMN
GLVYDVTTGKQYYEKGGTYHWLAGKDSSTDLNLIGGDIIKRKYPVIGKLVQ
>b5MY_EKD84736.1 neuron derived neurotrophic factor [uncultured bacterium]
MKRIIFSAGLIFLFLLVFLVLVNFYINSRYKPDVSLQPTQTNEEAAKTFTQDELKKYNGTDADLPIYLAL
DGVVYDVTAGREFYASGGAYHYLAGKDASIPLHLVGADIIKNKYPVVGKLVK
>b5MY_OGE21801.1 hypothetical protein A2778_04905 [Candidatus Daviesbacteria bacterium RIFCSPHIGHO2_01_FULL_40_24]
MNSRYKPDVSLQPTQTNEEAAKTFTQDELKKYNGTDADLPIYLALDGVVYDVTAGREFYASGGAYHYLAG
KDASIPLHLVGADIIKNKYPVVGKLVK
>b5MY_KKQ43725.1 hypothetical protein US62_C0043G0008 [Candidatus Woesebacteria bacterium GW2011_GWA1_37_8]
MKHKKPPKNSKLVKLVTIISISSVVVVFLFSFAVKLYLFPKNKPANSSDWESRYFSGDELKKYNGTNPKL
PIYMGYEGKVYDVSAGAGFYAAGKTYNYLTGRDATAELNEVGVGEIIIRKYPVIGKIKN
>b5MY_KKR31495.1 hypothetical protein UT63_C0068G0005 [Candidatus Gottesmanbacteria bacterium GW2011_GWC2_39_8]
MINNVVFLVVLVFAFSSQFTRTQTQKSEETKVFKVSELSKFNGTDPNKPIYIAYNGEVYDVSGGRNFYKP
GGTYHFLAGRDSTSELNIAGGAIIKRKYPVIGVLRDNYER
>C1_b5M_KUO41884.1 cytochrome B5 [Hadesarchaea archaeon YNP_N21]
MRVFTKEELSRYNGKEGAPAYVAYNGKVYDVTGSFHWKGGKHHVLHDAGQDLTESIGRAPHTAELLEKFP
VVGVLRG
>C2_P04166.2|CYB5B_RAT RecName: Full=Cytochrome b5 type B; AltName: Full=Cytochrome b5 outer mitochondrial membrane isoform; Flags: Precursor
MATPEASGSGRNGQGSDPAVTYYRLEEVAKRNTAEETWMVIHGRVYDITRFLSEHPGGEEVLLEQAGADA
TESFEDVGHSPDAREMLKQYYIGDVHPNDLKPKDGDKDPSKNNSCQSSWAYWIVPIVGAILIGFLYRHFW
ADSKSS
>C2_pdb|1CXY|A Chain A, Structure And Characterization Of Ectothiorhodospira Vacuolata Cytochrome B558, A Prokaryotic Homologue Of Cytochrome B5
NETEATLPVFTLEQVAEHHSPDDCWMAIHGKVYDLTPYVPNHPGPAGMMLVWCGQESTEAWETKSYGEPH
SSLAARLLQRYLIGTLEEIT
